# A pivotal role for Nrf2 in endothelial detachment-implications for endothelial erosion of stenotic plaques

**DOI:** 10.1101/537852

**Authors:** Sandro Satta, Michael McElroy, Alex Langford Smith, Glenn R Ferris, Jack Teasdale, Yongcheol Kim, Giampaolo Niccoli, Tom Tanjeko Ajime, Jef Serré, Georgina Hazell, Graciela Sala Newby, Ping Wang, Jason L Johnson, Martin J Humphries, Ghislaine Gayan-Ramirez, Peter Libby, Filippo Crea, Hans Degens, Frank Gijsen, Thomas Johnson, Amir Keshmiri, Yvonne Alexander, Andrew C Newby, Stephen J White

**Affiliations:** Department of Life Sciences, Manchester Metropolitan University, Manchester M1 5GD UK; School of Mechanical, Aerospace and Civil Engineering (MACE), The University of Manchester, Manchester, M13 9PL, UK; Bristol Heart Institute, University of Bristol, Bristol BS2 8HW, UK; Department of Cardiovascular Medicine, Chonnam National University Hospital, 42 Jebongro, Dong-gu Gwangju, 61469 Republic of Korea; Fondazione Policlinico Universitario A. Gemelli IRCCS, Roma, Italia; Universita’ Cattolica del Sacro Cuore, Roma, Italia; Laboratory of Pneumology, Department of Chronic Diseases, Metabolism and Ageing, KULeuven, Leuven, Belgium; Bioinformatics Core Facility, Faculty of Biology, Medical and Health, University of Manchester; Wellcome Centre for Cell-Matrix Research, Faculty of Biology, Medicine & Health, University of Manchester, Manchester, M13 9PT, UK; Brigham and Women’s Hospital, Harvard Medical School, Boston, MA 02115 USA; Department of Cardiology, Biomedical Engineering, Erasmus MC, Rotterdam, The Netherlands; Department of Cardiology, Bristol Heart Institute Upper Maudlin St. Bristol BS2 8HW

## Abstract

Endothelial erosion of atherosclerotic plaques and resulting thrombosis causes approximately 30% of acute coronary syndromes (ACS). As changes in the haemodynamic environment strongly influence endothelial function and contribute to plaque development, we reconstructed the coronary artery geometries of plaques with thrombi overlying intact fibrous caps from 17 ACS patients and performed computational fluid dynamic analysis. The results demonstrated that erosions frequently occur within areas of stenosis exposed to elevated flow. We recapitulated this flow environment *in vitro*, exposing human coronary artery endothelial cells to elevated flow and modelled smoking (a risk factor for erosion) by exposure to a combination of aqueous cigarette smoke extract and TNFα. This treatment induced endothelial detachment, which increased with pharmacological activation of the antioxidant system controlled by transcription factor Nrf2 (encoded by NFE2L2). The expression of Oxidative Stress Growth INhibitor genes OSGIN1 and OSGIN2 increased under these conditions and also in the aortas of mice exposed to cigarette smoke. Sustained high level expression of OSGIN1+2 resulted in cell cycle arrest, induction of senescence, loss of focal adhesions and actin stress fibres, and dysregulation of autophagy. Overexpression of either Nrf2 or OSGIN1+2 induced cell detachment, which did not depend on apoptosis, and could be partially rescued by inhibition of HSP70 using VER-155008, or AMP kinase activation using metformin. These findings demonstrate that under elevated flow, smoking-induced hyperactivation of Nrf2 can trigger endothelial cell detachment, highlighting a novel mechanism that could contribute to ACS involving endothelial erosion overlying stenotic plaques.

## Introduction

Rupture of a thin-capped fibroatheroma (plaque rupture) provoke some two thirds of ACS, whereas local erosion of the intimal endothelium that lines the arterial lumen causes most of the remainder (reviewed [1–4]). Both events promote thrombus formation, restricting blood flow, potentially triggering arrhythmia and infarction. Yet plaque rupture and erosion occur on atherosclerotic plaques with markedly different histological features, suggesting substantial differences in the underlying mechanisms, and implying that distinct risk factors contribute to each process. For example, plaque rupture occurs on inflamed and lipid-rich plaques with a greater degree of calcification. In contrast, sites of erosion generally occur on smooth muscle cell- and proteoglycan-rich plaques, specifically containing high levels of versican and the glycosaminoglycan hyaluronan and low levels of biglycan, containing few inflammatory leucocytes [1, 5–9]. In addition, the thrombi overlying eroded plaques display more organisation, indicative of a longer existence, potentially up to 7 days before presentation of clinical symptoms [10–12].

While the understanding of plaque rupture has progressed considerably, plaque erosion remains enigmatic. Histopathology studies have identified altered sub-endothelial matrix composition, which may impair endothelial adhesion and alter endothelial behaviour. Moreover, these studies have identified smoking, a well-known cause of endothelial dysfunction, as a risk factor for endothelial erosion [5, 13–15]. Long-term, heavy smokers have elevated circulating mediators of inflammation, such as tumour necrosis factor-alpha (TNFα), that exacerbate dysfunction of endothelial cells (ECs) [16]. In addition, we have demonstrated that fresh aqueous cigarette smoke extract synergises with TNFα to increase the expression of Nrf2-target genes in human coronary artery endothelial cells, in a shear-dependent manner [17–19]. In summary, altered matrix composition, inflammation and the effects of smoking-related factors are all implicated in endothelial erosion.

Recent advances in intravascular imaging, particularly optical coherence tomography (OCT)-based methods, allowed rupture to be distinguished from erosion in patients with ACS [20, 21]. OCT-defined – acute coronary syndrome with an intact fibrous cap (ACS-IFC) - erosion is a diagnosis by exclusion of rupture or the presence of calcified nodules and is not without controversy [22]. However, OCT-defined ACS-IFC has a similar frequency to that defined by histological studies of erosion, supporting its relevance [3, 21]. Moreover, the largest study to date that utilised OCT-diagnosis of plaque rupture or erosion in 822 STEMI patients [21], identified a propensity for plaque erosion in smokers and women <50yr compared to plaque rupture patients, consistent with post mortem observations [7, 23, 24]. Yet, other traditional risk factors for acute coronary syndromes (for example, diabetes, hyperlipidaemia and hypertension) segregated with plaque rupture, highlighting that the mechanisms of regional intimal endothelial detachment in patients differ from those of plaque rupture and require better definition.

The haemodynamic environment exquisitely regulates the behaviour of endothelial cells: it modulates metabolism, cell shape, cytoskeleton arrangement, proliferation, permeability, response to inflammatory stimuli and apoptosis [25–29]. Thus, haemodynamics strongly influences plaque development, progression and features of lesions that determine their propensity to rupture [30–33]. Emerging evidence also supports the involvement of haemodynamics in endothelial erosion. Studies of human lesions show heightened endothelial apoptosis in regions of disturbed flow, distal to the point of maximal plaque stenosis [34]. Recent experimental work has implicated disturbed flow in promoting endothelial cell death, amplified by neutrophil extracellular trap (NET) formation in regions of endothelial denudation [35, 36]. Yet, the relationship of the site of clinically-relevant endothelial erosion to plaque architecture and haemodynamics remains ill-defined. Here, we studied in a series of patients the geometry and haemodynamic environment permissive for OCT-defined plaque erosion, identifying that while some erosions occurred in regions of disturbed flow that would promote apoptosis, they also frequently occur in regions of elevated flow, suggesting two separate mechanisms of endothelial erosion. In this study, we created a cell culture approach to investigate elevated-flow-dependent detachment, observing an Nrf2-amplifed loss of adhesion in primary human coronary artery ECs (HCAECs) when also exposed to TNFα and the water-soluble components of cigarette smoke. We describe the molecular mechanisms responsible for endothelial cell detachment under these conditions, identifying roles for OSGIN1 and OSGIN2 in mediating detachment. These observations suggest novel targets for pharmacological intervention to limit superficial erosion.

## Methods

Please see supplementary information for full description of the methods.

## Results

### Establishing the Plaque Luminal Geometry and Haemodynamic Environment Permissive for Endothelial Erosion

To gain insights into mechanisms responsible for endothelial erosion from plaques, we first sought to define the luminal geometry and compute the local haemodynamic environment at the site of OCT-defined erosions in ACS patients. We reconstructed the coronary artery luminal architecture from 17 OCT-defined erosion patients containing thrombus overlying an intact fibrous cap. This analysis used a combination of geometrical information derived from bi-plane cineangiography and from the high-definition lumen profile derived from OCT, determined beneath the thrombus to establish the structure prior to ACS (Figure 1A and supplementary data). Computational fluid dynamic (CFD) assessment of haemodynamic environments under both resting and stress conditions were simulated using artery-specific waveforms (Tables S5). Fourteen of the 17 culprit plaques had an average ~40% diameter (60% area) stenosis (Table S6), in agreement with histologically defined data [7, 10]. The predominant flow characteristic at the sites of adherent thrombi was elevated flow, with ~6.5-fold elevation of the time averages wall shear stress (TAWSS) (Figure 1C, 1E), reflecting the stenotic nature of these plaques. Nine of these 14 sites of adherent thrombi showed no increase in oscillatory shear index (OSI), while 5 showed a modest elevation of OSI, correlating with an extension of the thrombi past the point of minimum lumen area (e.g. Figure 1B). The three remaining sites of adherent thrombi displayed clear differences in haemodynamic features, with modest or no stenosis, modest changes in TAWSS values and a pronounced elevation of OSI. These observations suggest that at least two distinct hemodynamic environments can predispose to plaque erosion.

**Figure 1.**
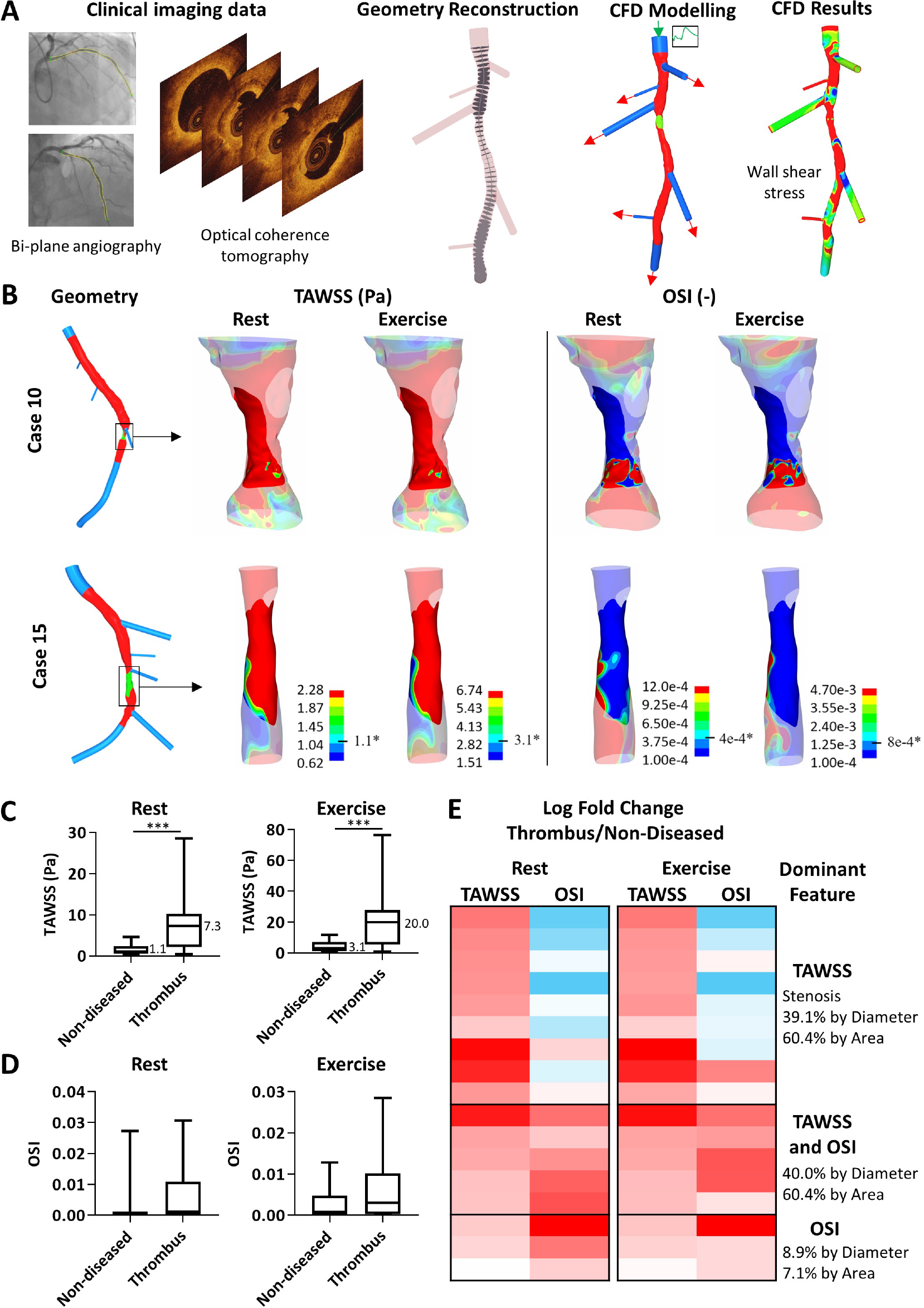
Probing the haemodynamic conditions permissive for plaque erosion. A) optical coherence tomography (OCT) and bi-plane angiography imaging data of coronary arteries were collected and used to reconstruct lumen geometries (n=17). Red sections are reconstructed using a hybrid OCT/bi-plane angiography method to produce a lumen profile to a high accuracy. The blue sections represent the geometry reconstructed using bi-plane angiography. The adhered thrombus is delineated by the green surface, extracted from OCT data. Side branch diameters were measured using OCT imaging, flow extensions were then added and the geometry discretised. Both at ‘rest’ and ‘exercise’ pulsatile flow conditions are simulated for 4 cardiac cycles, with separate waveforms used for the right, left/left anterior descending and circumflex arteries, depending on the location of the culprit lesion. Haemodynamic flow results were post-processed to quantify additional wall shear-based haemodynamic metrics of interest. B) Representative images of vessels with severe (case 10) and moderate (case 15) levels of stenosis (see Table S2). The thrombus is the opaque portion of the metrics, whilst the remainder of the lumen is semi-transparent. Two metrics presented here are time-averaged wall shear stress (TAWSS) and oscillatory shear index (OSI). Minimum and maximum values for the legends are the lower and upper quartiles of the respective metrics averaged across the rest and exercise cases separately. Proximal end at top of image, full data presented (Figure S4, Tables S8-S9) C) CFD was used to calculate the TAWSS within a non-diseased area and under the thrombus (n=17) for both rest and exercise conditions. There was a significant elevation by the Wilcoxon signed rank test between non-diseased and thrombosed in both models. The median value is displayed on the graph, *** p<0.001. D) The OSI was also calculated by CFD, however overall there was no significant difference between groups. E) The log_2_ fold change between the non-diseased and thrombus areas were compared for each individual model at rest and exercise. The change was displayed as a heat map, each row depicts the results from one patient, with red representing an increase in TAWSS or OSI and blue indicating a decrease. In 14 cases a substantial increase in TAWSS appears to be the dominant feature, with 5 of these also showing an increase in OSI reflecting adhered thrombus extending distal to the point of maximum stenosis; increased OSI is the dominant feature the 3 remaining cases with little change in TAWSS.

### Nrf2 Participates in Elevated Flow Endothelial Cell Detachment

To investigate potential mechanisms, we created *in vitro* conditions to simulate exposure of ECs with known risk factors for endothelial erosion. Confluent monolayers of HCAECs were exposed to oscillatory (OSS), normal laminar (LSS) or elevated laminar (ESS) shear stress and treated with 10% aqueous Cigarette Smoke Extract (CSE) [17, 18] and/or 5ng/ml TNFα. Cell detachment was observed at elevated shear stress (ESS) only after the addition of both CSE and TNFα (~30% cell loss, Figure 2A, 2B). Treatment with an apoptosis inhibitor (Z-VAD-FMK), a broad spectrum matrix metalloproteinase inhibitor (GM6001), rosuvastatin, or necrostatin-1 did not prevent cell loss (Figure S5-S8). To determine the role of Nrf2 in cell detachment, pharmacological activators of Nrf2 (2.5μM sulforaphane or 10μM isoliquiritigenin) were added to the cultures with CSE and TNFα. Further activation of Nrf2 triggered almost complete cell detachment (Figure 2B), indicating that chronic hyperactivation of Nrf2 contributes to, rather than protects from cell loss. To investigate further the actions of Nrf2 on HCAEC function we overexpressed Nrf2 in HCAEC and observed a profound inhibition of proliferation (Figure 2C), implicating Nrf2-dependent surveillance of free radical and reactive oxygen species stress in regulating HCAEC replication, with consequences for the ability of the endothelium to repair. In addition, when we overexpressed Nrf2 in HCAEC and subsequently exposed the cells to shear stress using an orbital shaker [39, 40], significant cell detachment was observed (Figure 2D), again implicating Nrf2-regulated gene expression in HCAEC detachment.

**Figure 2.**
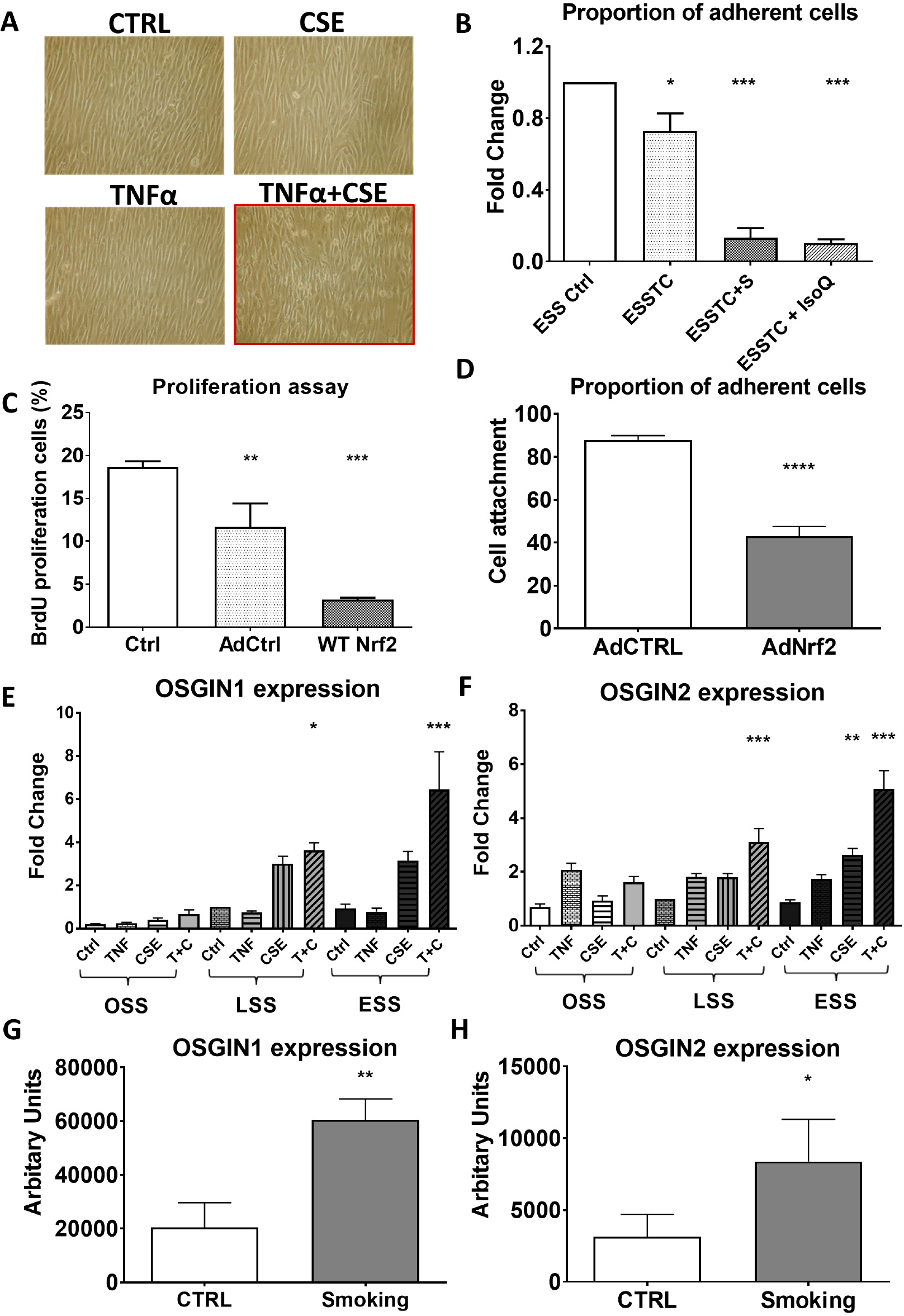
A) Photomicrographs of human coronary artery endothelial cells (HCAEC), cultured under elevated flow post-treatment. Neither 5ng/ml TNFα nor 10% CSE alone had any effect on cell adhesion versus control in ESS conditions. However, the combination of TNFα and CSE resulted in a significant loss of cell adhesion, (framed in red) B) Quantification of cell number in elevated laminar shear stress control (ESS), or with addition of TNFα and CSE (ESSTC) One-way ANOVA Bonferroni *P<0.05, n=3). Further activation of Nrf2 using sulforaphane (Sulf-2.5μM) (*5-fold reduced adhesion vs ESSTC P<0.05, n=3), or isoliquiritigenin (IsoLiq-10μM) (*9-fold reduced adhesion vs ESSTC P<0.05, n=3), increased cell loss (One-way ANOVA Bonferroni ***P<0.001, n=3). C) BrdU proliferation assay. HCAEC cells treated with AdNrf2 virus for 16 h. The percentage of BrdU positive cells from the images was calculated. Each value represents the average obtained from n=6 individual experiments using HCAEC from different donors. Data are presented as the mean ± standard deviation. One-way ANOVA Bonferroni *P<0.05 and **P<0.01, compared with control. D) Adenoviral overexpression of Nrf2 (200pfu/cell and 200pfu/cell AdCTRL combined to match later experiments) promotes 50% of cell detachment compared to AdCTRL (400 pfu/cell) (T-test ****P<0.0001). E-F) OSGIN1 and OSGIN2 mRNA expression in HCAEC cultured under oscillatory (OSS), normal laminar (LSS) or elevated laminar shear stress (ESS) under control, 5ng/ml TNFα or 10% CSE or both (Two-way ANOVA *P=0.05, **P<0.01, ***P<0.001). G-H) Immunohistochemical staining was carried out on 8μm sections of aortas from mice exposed to cigarette smoke for 3 months vs control (CTRL). Expression of OSGIN1 and 2 was significantly increased in mice exposed to smoke (**P<0.01, n=8 OSGIN1 vs 7 CTRL; *P<0.05 n=7 OSGIN2 vs 7 CTRL; T-test).

Under elevated flow, CSE and TNF treatment, we observed a concomitant increased expression of two Nrf2-regulated genes OSGIN1 and OSGIN2 (Figure 2E). This result suggests a complex shear-dependent pattern of regulation, with the highest level of expression at elevated flow with the addition of CSE and TNFα. In addition, we observed augmented OSGIN1 and OSGIN2 expression in the aortas of mice subjected to chronic tobacco smoke. Mice exposed to cigarette smoke for 3 months demonstrated significantly higher expression of OSGIN1 and OSGIN2 in their aortas, compared to control mice (Figure 2G, 2H, S6).

### OSGIN1 and OSGIN2 Function in Human Coronary Artery Endothelial Cells

OSGIN 1 and OSGIN2 have a similar predicted structure with 40.8% amino acid sequence identity and demonstrate amino acid sequence conservation across species (Figure S10-S13). To dissect the potential role of OSGIN1 and OSGIN2 in elevated flow-dependent endothelial cell desquamation, the effect of overexpression of both genes was studied in HCAEC. OSGIN1 and OSGIN2 localised to the nucleus (Figure 3A), despite the lack of an obvious nuclear localisation sequences using cNLS Mapper [41], or SeqNLS [42]. Both OSGIN1 and OSGIN2 inhibit proliferation of HCAEC, with cells accumulating in S-phase (OSGIN1 early S-phase, OSGIN2 post S-phase, early telophase) and not proceeding to cytokinesis (Figure 3B, 3C and Figures S14A-B). Large multi-nucleated cells accumulated (Fig S14C-D, S15, S16 and 4A-C), corresponding with a significant increase in senescence-associated β-galactosidase staining and increase in expression of p-16 and p-21 (Figure 3D, 3E, 3F, S17); however no signature of apoptosis was observed (Figure 3G, Figure S18).

**Figure 3:**
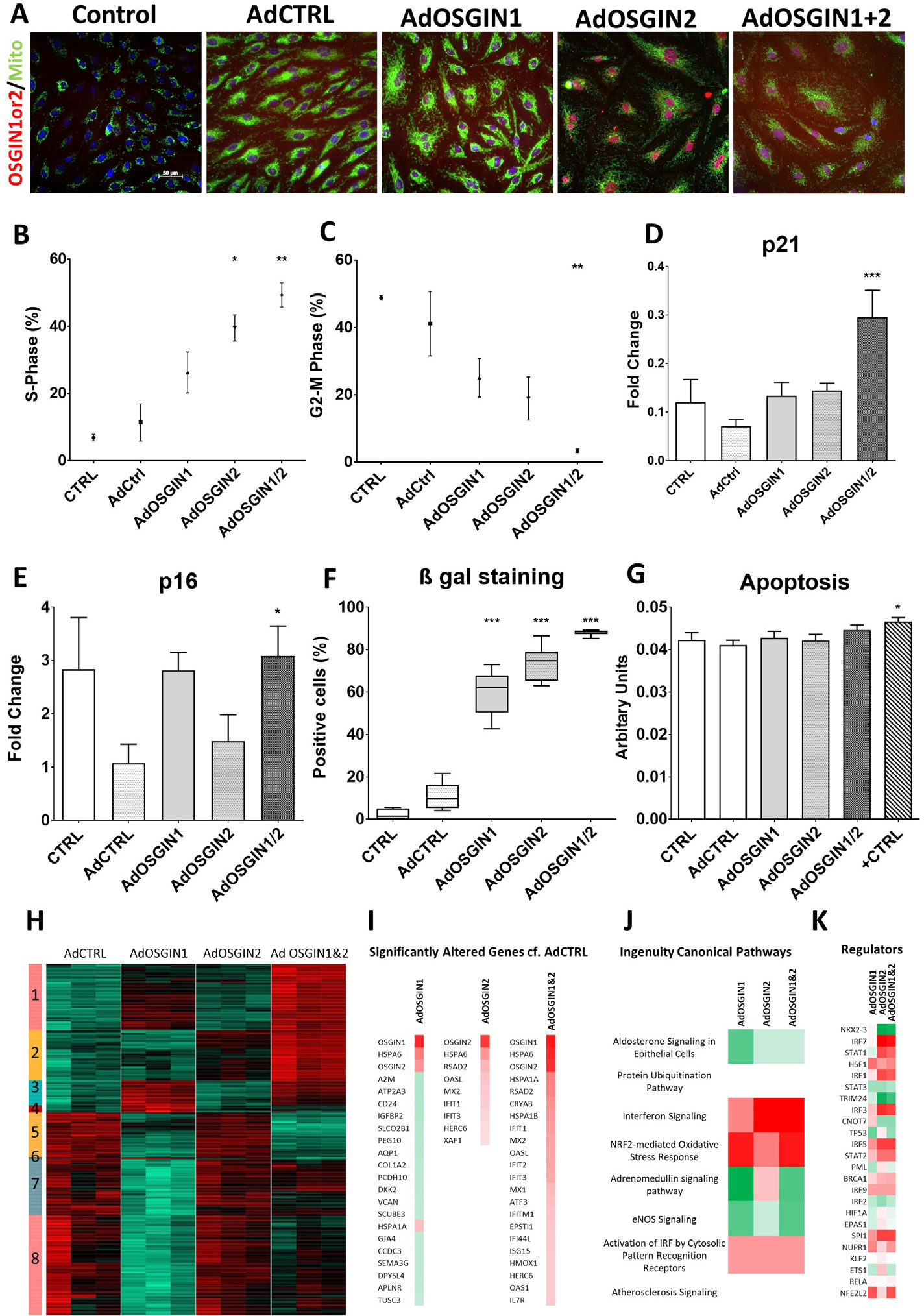
A) Immunocytochemical staining for OSGIN1 and OSGIN2 (in red) shows their nuclear localisation. Mitochondrial staining (in green) confirmed no translocation of OSGIN1 and OSGIN2. B-C) Flow cytometry analysis after adenoviral-mediated overexpression of OSGIN1 and OSGIN2 showed inhibition of cell cycle in human coronary artery endothelial cells, with cells accumulating in S-phase, without proceeding to division (*P<0.05, and ***P<0.001, n=3). D-E) Changes in mRNA expression of p21 and p16 with OSGIN overexpression (***P<0.001; *P<0.05 n=6). F) Increased staining for senescence-associated β-galactosidase and upregulation of P16 and p21 demonstrates the induction of the senescence pathway and lysosomal accumulation by OSGIN1+2 (***P<0.001 n=3) images (Figure S17). G) Apoptotic activity was evaluated using Apotoxglow assay (Promega). AdOSGINs overexpression didn’t show any apoptotic activity. Positive control (0.2mM H_2_O_2_). Data is presented as mean ± SEM (*p<0.05, n = 3). Additional data Figure S18. RNASeq transcriptomics was performed on three different HCAECs batches transfected with adenoviral vector control or overexpressing OSGIN1, 2 or 1+2. H) Heat map of gene expression significantly changed between adenoviral control and adenoviral overexpression of OSGIN1, OSGIN2 or OSGIN1+2; green indicated decreased, black no change and red increased gene expression. The cluster analysis was carried out on the differentially expressed genes identified with DESeq2 using a p-adjusted cut off of 0.05, absolute log2 fold change cut off of 0.5, and base Mean cut off of 50. The Pearson distance was clustered with the hclust function and plotted with an R package of gplots. Clustering analysis revealed 8 clusters. I) The genes with the most significant/largest fold change by overexpression of OSGIN1, OSGIN2 or OSGIN1+2 compared to control are shown. Red indicates an increase in expression and green a decrease. J) Ingenuity pathway analysis predicted upstream transcriptional regulators for genes changing within the heat map, red indicates a predicted activation, white no change and green a decrease. K) Ingenuity pathway analysis of the top predicted canonical pathways, red indicates a predicted activation and green a decrease. White indicates significant changes in genes in this pathway but do not consistently indicate an increase or decrease in the pathway, suggesting dysregulation.

RNASeq transcriptomics were performed on HCAEC, transduced with adenoviral control (AdCTRL), AdOSGIN1, AdOSGIN2, or AdOSGIN1+2 (Figure 3H-K). The relative expression of 360 different genes changed significantly over the three conditions. Clustering analysis identified eight clusters (Figure 3H), cluster 1 being significantly enriched in genes associated with the activation of the Ingenuity IPA canonical pathway NRF2-mediated Oxidative Stress Response, inhibition in eNOS Signalling and alteration in protein ubiquitination Pathway, unfolded protein response (full list, Table S10). These changes are predicted to be driven by the activation of the transcription factor genes HSF1 and NFE2L2 (Nrf2) amongst other predicted regulators (Table S10). Cluster 2 associates with the activation of Interferon Signalling (predominantly type I) and driven by the activation of STAT1 and 2, IRF1, 3, 5, 7 and 9, and inhibition of TRIM24 (Table S14). Cluster 3 was dominated by the Nrf2-mediated Oxidative Stress Response pathway driven by Nrf2 activation (Table S15). Full detail of all clusters is in Tables S10-S20. With AdOSGIN1, 235 genes changed significantly (p adjusted <0.05), 9 with AdOSGIN2 and 169 with AdOSGIN1+2 compared to control (Figure 3I). Eleven of the 20 most upregulated genes participate in interferon signalling or are down stream of interferon. The significant increase in genes associated with the interferon signalling Ingenuity canonical pathway highlights the importance of interferon in regulating the genes altered by OSGIN1, 2 or 1 and 2 (Figure 3J) and the predicted activation of upstream regulator genes IRF7, STAT1, IRF1, IRF3, IRF5, STAT2, BRACA1, IRF9 and SPI1 (Figure 3K). Six of the 20 genes with the largest differences in expression associate with proteostasis: 5 of which interact with the key proteostasis regulator BAG3 (HSPA6, HSPA1A, CRYAB, HSPA1B and ISG15; Ingenuity IPA database), which is itself upregulated (Figure 4H). Furthermore, other interaction partners of BAG3: STIP1, HSPB1, HSPB8, NQO1, HSP90AA1, HSP90AB1, DNAJB1, DNAJB6 HSPA4, P4HA2, SQSTM1 (p62), HSPA8, TRIM69 all increase (Table S20). In summary, the transcriptomic analysis suggests alterations in interferon, Nrf2 and HSF1-regulated gene expression and pronounced changes in BAG3-regulated proteostasis.

**Figure 4:**
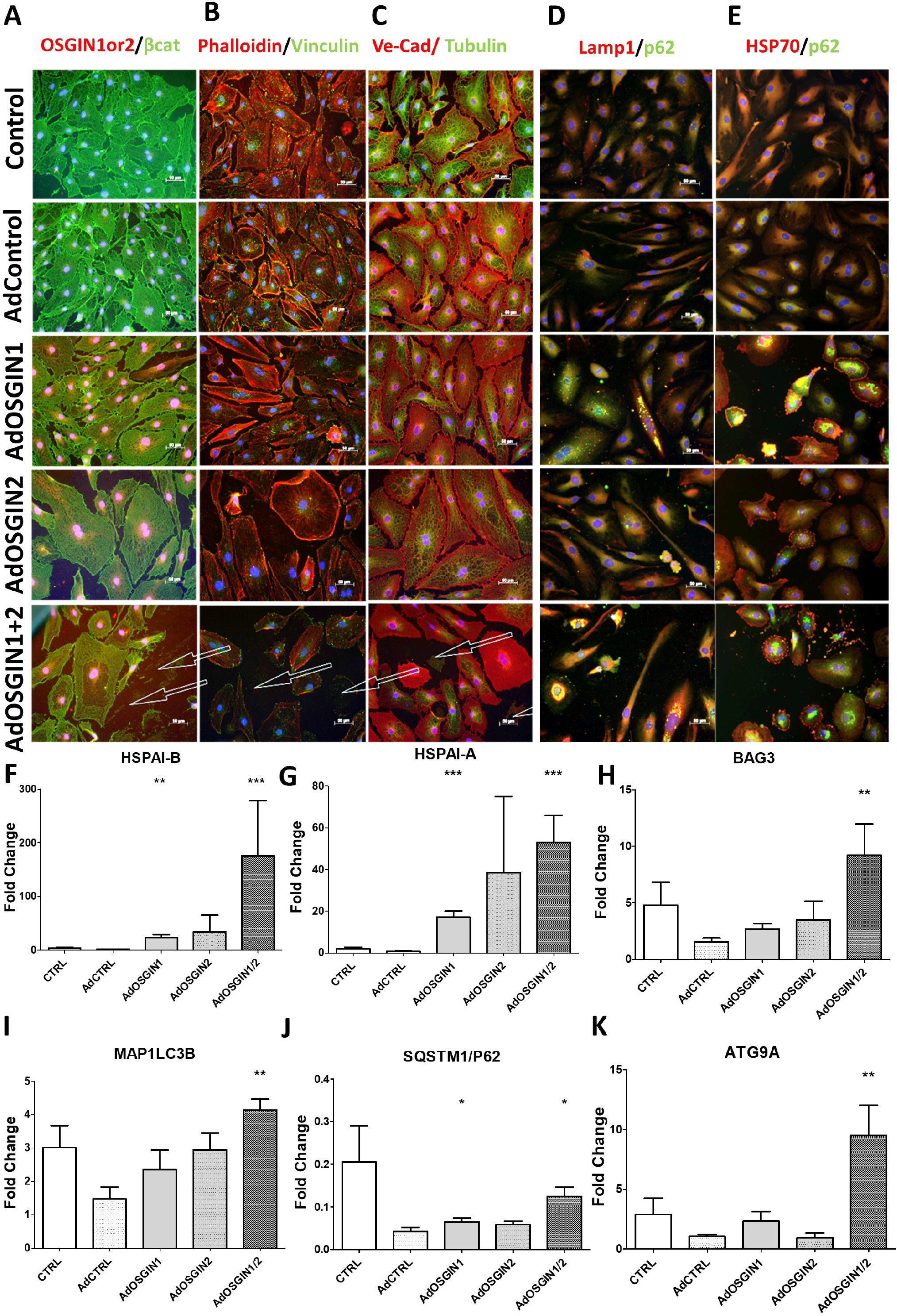
A-C) Immunocytochemical analysis of HCAECs with adenoviral-mediated overexpression of OSGIN 1 + 2. β-catenin (marker of intercellular junctional stability), vinculin (focal adhesions) and phalloidin (Actin), Tubulin and VE-Cadherin (intercellular junctions in Red), visualized by immunofluorescence microscopy. OSGIN1+2 overexpression profoundly affects cell structure, reduces cytoskeletal integrity and focal adhesions with cells detaching even in static culture denoted by arrows (quantification in Figure S11). D-E) Immunofluorescence of SQSTM/p62 (in green) and LAMP1 or HSP70 (in red), demonstrate an accumulation of SQSTM/p62 and LAMP1 positive vesicles, indicative of a block in autophagic flux. F-K) Changes in mRNA expression of key regulators of the chaperone-mediated autophagy pathway by OSGIN1+2 overexpression F) HSPA1A; G) HSPA1B; H) BAG3; I) MAP1LC3B; J) SQSTM1/P62; K) ATG9A (**P<0.01 and ***P<0.001; N=6) protein level quantification (Figure S23).

### Upregulation Of OSGIN1 And OSGIN2 Dysregulate Actin And Tubulin Networks, Focal Adhesions and Autophagy

Stable cell adhesion requires the integrity the cytoskeleton and focal adhesions. Overexpression of OSGIN1 and OSGIN2 triggered profound alterations in cell structure, with a collapse of the actin and tubulin networks (Figure 4A-C), with an overall reduction in staining for actin, tubulin and vinculin (Figure S19). Concomitantly autophagic vesicles accumulated within HCAECs (Figure 4D-E) and expression of genes involved in HSP70/BAG3-controlled chaperone-mediated autophagy pathway (Figure 4F-K).

### OSGIN1 and OSGIN2, Proteostasis and Nrf2 Regulate HCAEC Adhesion

Overexpression of OSGIN1 and OSGIN2 in HCAEC triggered cell detachment (Figure 4A) equivalent to Nrf2 overexpression (Figure 2D), quantified under exposure to shear stress using the orbital shaker model (Figure 5A, 5B). Many detached cells retained membrane integrity as assessed by trypan blue exclusion, indicating a non-apoptotic mechanism. Incubation with low doses of chloroquine (150μM) or bafilomycin (50nM) to inhibit autophagy (Figure 5C, 5D), or moderate doses (300μM, 100nM Figure S20), also triggered cell detachment with similar maintenance of membrane integrity. Co-treatment of endothelial cells with OSGIN1+2 overexpression and either chloroquine or bafilomycin did not enhance cell detachment, or alter cell integrity, suggesting a reduction in autophagic flux by either chloroquine or bafilomycin, or by OSGIN1+2 overexpression can regulate cell adhesion.

**Figure 5:**
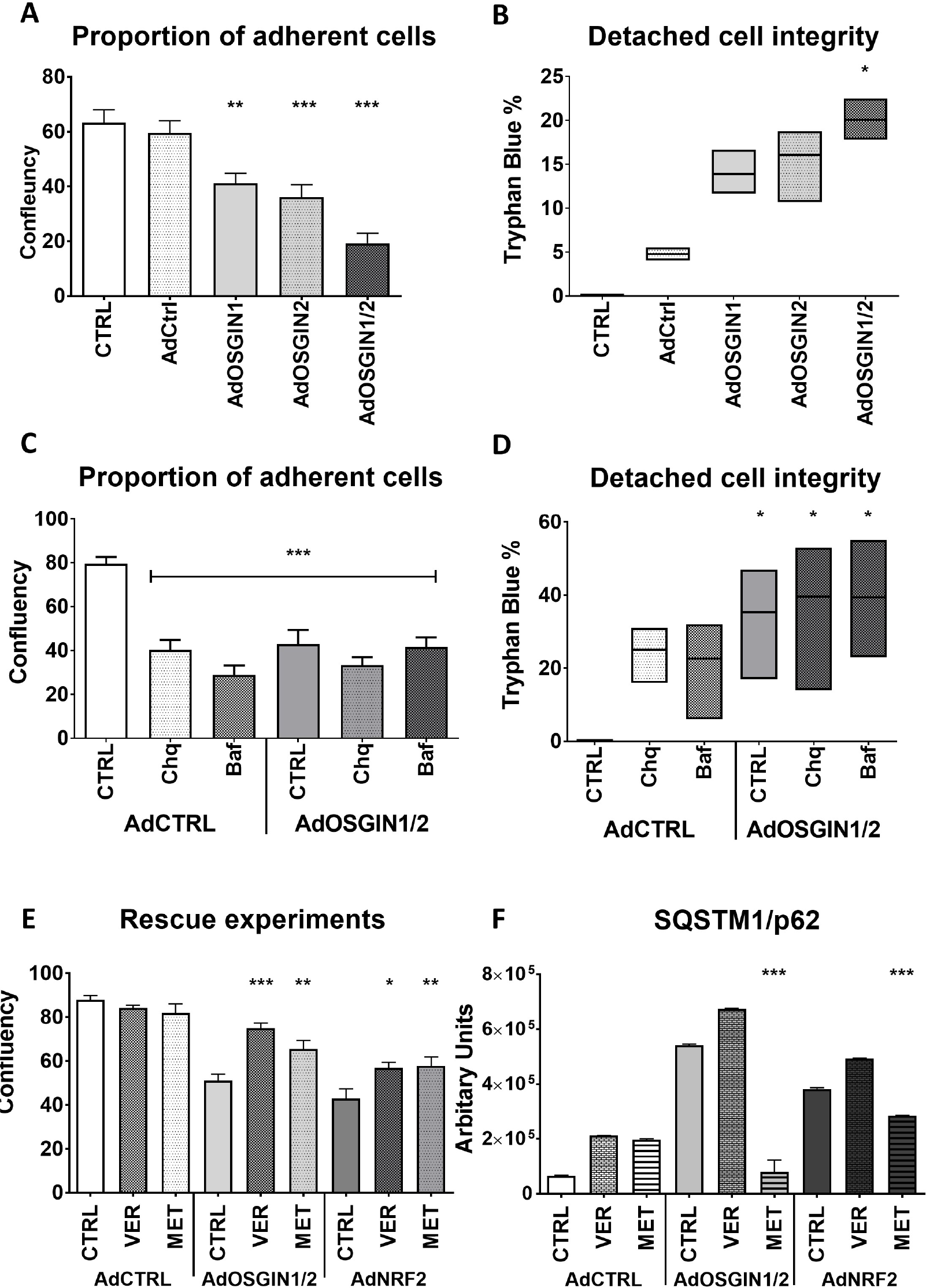
Quantification of endothelial cell detachment using orbital shaker shear stress model. A-B) **A**denoviral overexpression of OSGIN1+2 triggers cell detachment (**P<0.01, ***P<0.001 v AdCTRL; n=3, One-way ANOVA), with detached cells displaying a significant maintenance of cell membrane integrity (*P<0.05 vs AdCTRL; n=4 One-way ANOVA). C-D), chloroquine (150μM), bafilomycin (50nM), or OSGIN1+2 overexpression induced comparable, non-synergistic detachment (**P<0.01 and ***P<0.001 v AdCTRL, n=4, Two-way ANOVA), with a similar maintenance of cell membrane integrity (*P<0.05 v AdCTRL, n=4, One-way ANOVA). E) co-treatment with Ver155008 (15μM), or Metformin (100μM) reduced OSGIN1+2 or Nrf2-mediated cell detachment; (*P<0.05, **P<0.01, ***P<0.001, n=3, Two-way ANOVA). F) Metformin, but not VER-155008 treatment reversed SQSTM1/p62 protein accumulation following AdOSGIN1+2 and AdNRF2 overexpression (***P<0.001, n=3, Two-Way ANOVA).

### HSP70 Inhibition or Activation of AMPK with Metformin Limits Detachment

Based on the observation that OSGIN1+2 overexpression markedly enhanced the HSP-70/BAG3 proteostasis pathway, we dissected a potential role of dysregulated chaperone-mediated autophagy in cell detachment by inhibiting HSP70 nucleotide binding site using VER-155008, or activating AMP kinase with metformin. Either VER-155008 or metformin reduced cell detachment that was induced by either OSGIN1+2 or Nrf2 overexpression and modified the accumulation of autophagic vesicle markers p62 and LAMP1 (Figure 5E, 5F, S18, S20), supporting the hypothesis that OSGIN1+2 dysregulation of chaperone-mediated autophagy mediates Nrf2-mediated cell detachment.

## Discussion

Endothelial erosion of plaques mediates a substantial and possibly growing proportion of ACS [1–4]. While plaque rupture allows communication of blood with the necrotic core initiating the coagulation cascade, plaque erosion exposes the much less thrombogenic sub-endothelial matrix, predominantly attracting platelet adhesion and ‘white’ thrombus formation, a process with a possibly slower thrombus accumulation and progression to clinical events [10–12]. This situation may permit a management strategy that avoids PCI and stent deployment in patients with ACS due to plaque erosion[43], a proposition currently being tested in Erosion II (NCT03062826). In addition, two studies reported that patients who had plaque rupture are at 1.72 to 2.5-fold greater risk of major adverse cardiac events (MACE) within 2 years, compared to those who had a plaque erosion [44, 45], highlighting a potentially distinct clinical outcome for these two mechanisms of ACS. Yet, the need for OCT to discriminate between plaque rupture and erosion mandates a similar approach to their management in routine clinical practice. Further development of less invasive diagnostic biomarkers could advance the ability to distinguish these two aetiologies of ACS to guide more individualized treatment strategies [46].

The data presented here advances understanding of the pathophysiology of plaque erosion. Firstly, this study describes the haemodynamic environments permissive for plaque erosion. Reconstruction of vessel geometry from 17 instances of OCT-defined erosion identified the predominant haemodynamic feature as highly elevated flow. This observation agrees with those of Dai *et. al.* [21] who observed 96% of thrombi either proximal to, or within the minimal lumen area (point of maximum stenosis), where the flow is predicted to be elevated and laminar, in 209 cases of OCT-defined erosion, a result that is supported by another independent study [47]. If causal, this association could result from the profound influence of the haemodynamic environment in regulating endothelial function and dysfunction [25–29], with the response to elevated flow eliciting a distinct pattern of gene expression [17, 37]. We and others have previously demonstrated that different flow patterns (and in our case pathologically-relevant elevated flow) changes endothelial behaviour and in particular the response to noxious stimuli that induce endothelial dysfunction [17, 37, 38]. In addition, the CFD simulations and subsequent statistical post-processing indicted that several OCT-defined erosions occurred in regions with modest or no stenosis, where the predominant flow feature was oscillatory shear stress (defined through OSI) [48, 49]. Low time averaged wall shear stress or elevated OSI values strongly correlate with the focal predilection sites for atherosclerosis [32, 50–52] and tend to activate endothelial cells, priming them for inflammatory activation and apoptosis [28, 34, 53]. Inducing endothelial apoptosis experimentally can initiate thrombosis [54] supporting a role for apoptosis triggering endothelial erosion. An independent case report corroborates our observation that erosions can occur under conditions of oscillatory shear stress [55]. OCT-defined erosion requires imaging an intact fibrous cap, precluding assessment of cases with high residual thrombus burden and accurate assessment of the proportion of plaque erosion cases that occur in elevated or oscillatory flow environments.

Secondly, we identified a hitherto unknown mechanism that contributes to endothelial detachment in vitro and potentially over stenotic plaques. Exposure of HCAEC to elevated flow, cigarette smoke extract and TNFα in vitro triggered cell detachment, aggravated by pharmacological activation of Nrf2. Augmented OSGIN1 and OSGIN2 expression accompanied Nrf2 activation in vitro, and also occurred in the aortas of mice exposed to cigarette smoke in vivo. Further, overexpression of either Nrf2 or OSGIN1+2 is sufficient to trigger endothelial cell detachment. Further dissection of the effects of OSGIN1 and OSGIN2 in HCAEC revealed that their overexpression inhibits proliferation and dramatically effects cell behaviour, increasing markers of senescence, altering cell size and shape, dysregulating the cytoskeleton and focal adhesions. The significantly increased expression of the HSPA1A, HSPA1B (HSP70) and BAG3, genes related to proteostasis, potentially contributes to this cellular response, as indicated by the accumulation of p62-labelled vesicles and the observation that low dose chloroquine or bafilomycin that inhibits autophagic flux also triggered detachment of HCAEC.

Finally, we demonstrated that inhibition of HSPA1A/HSPA1B nucleotide binding site with VER155008 reduced cell detachment suggesting a role of chaperone-mediated autophagy in regulating cell adhesion. This conclusion derived support from the observation that AMP kinase activation by metformin, which promotes macroautophagy potentially bypassing the blockade in chaperone-mediated autophagy, normalised p62 protein levels, suggesting a clearance of autophagic vesicles, and stabilised the attachment to substrate of HCAEC overexpressing either Nrf2 or OSGIN1+2. The HSP70/BAG3 complex participates in a range of disease states [56], represents a novel axis for therapeutic intervention, and highlights the necessity to dissect further the role of dysregulated autophagy in the modulation of endothelial adhesion. The ability of metformin to partially rescue the deficit on adhesion may have relevance to the observed reduction in ACS in diabetic patients treated with metformin [57, 58]. Future work should investigate this possible connection.

The transcriptional programme regulated by Nrf2 includes many genes with an important role in protection from free radical and oxidative stress [19]. Laminar shear stress modestly increases Nrf2 activity in the endothelium, contributing to the atheroprotective signalling induced by laminar flow [59–63]. Cigarette smoke contains high levels of free radical, reactive oxygen and nitrogen species [64] and activates Nrf2 in the lung [65]. We have previously shown that elevated flow and inflammation act together to trigger the highest level of many Nrf2-regulated genes in HCAEC, indicating a potential flow-regulated synergism between oxidant stress and inflammation increasing Nrf2 activity [17]. Therefore, haemodynamic and environmental activators of Nrf2 may impair adhesion of endothelial cells overlying stenotic plaques. In addition to cigarette smoke, phytochemicals (e.g. sulforaphane and isoliquiritigenin used in this study), pollutants including diesel particulates PM<10, which can pass through the lung [66], hydrogen sulphide [67] and additional sources of free radicals and reactive oxygen species (or suppressors of anti-oxidant defences) may potentiate Nrf2-dependent gene transcription, all of which might have with implications for endothelial adhesion. Global deletion of Nrf2 (Nrf2-/-) in hypercholesterolemic mice, reduces rather than increases atherosclerosis, despite the predicted reduction in antioxidant defence [68, 69] and inflammation [62]. While this has been explained previously by effects on lipid metabolism [68], a reduction in expression of scavenger receptor CD36 reducing foam cell formation [69], from NLRP3 inflammasome co-activation by cholesterol crystals [70], our results described here suggest that Nrf2-/- mice might harbour a greater capacity for endothelial adhesion and repair. These observations raise a cautionary note on modulation of Nrf2 as an adjunctive therapy to prolong lifespan [71, 72]. The multiple levels of control that exist to regulate Nrf2 activity strengthen this concern [19], by indicating the necessity for tight regulation of the Nrf2-dependent transcriptional programme.

Our data also suggest that interferon α signalling may potentiate the gene expression pattern elicited by OSGIN1+2 overexpression, as numerous regulated genes had IRF3/7 binding sites in their promoters. In this context, the large increase of HSP70 observed downstream of Nrf2 or OSGIN1+2 holds interest. Extracellular HSP70 can mediate a protective signalling function in endothelial cells [73], mediated through Trif-dependent TLR4 signalling [74], which activates IRF3/7. Further *in vitro* experiments and measurements in patient samples should probe the contribution of interferon α in endothelial adhesion and the implicit potential for viral or bacterial infection to accentuate the signalling pathways identified in this study.

The sub-endothelial matrix of eroded plaques contains abundant hyaluronan and its binding partner versican [5]. The recent observation that OCT-defined plaque erosion patients have increased HYAL2 (the enzyme responsible for degrading high-molecular-weight hyaluronan to its proinflammatory 20-kDa isoform) and the hyaluronan receptor CD44v6 adds weight to a potential causal relationship [9]. Hyaluronan and versican may regulate vascular inflammation in atherosclerosis [75, 76]. Hyaluronan (and biglycan) can bind to and activate TLR2 and TLR4 [77, 78]. TLR2 signalling may predispose to endothelial desquamation as exposure to disturbed blood flow and engagement of TLR2 (potentially by hyaluronan fragments) promoted endothelial apoptosis and desquamation, which was enhanced by neutrophil NET formation [4, 35, 36, 79].

However, exposure to laminar shear inhibits apoptosis in endothelial cells overlying atherosclerotic plaques [34]. Furthermore, our experiments define a mechanism of endothelial detachment without any signature of apoptosis, and with detached cells maintaining significant membrane integrity, consistent with two independent mechanisms erosion potentially reflecting different haemodynamic environments. Further studies should address the differential effects on cell death pathways of hyaluronan and HSP70 signalling through TLR2 and TLR4 under various flow conditions. In addition, the relative adhesive quality of a sub-endothelial matrix containing a higher proportion of hyaluronan and versican and possible contributions of extracellular matrix-degrading hydrolases requires elucidation, especially under elevated flow conditions [4, 38, 80]. These observations put a spotlight on the role of TLR2 and TLR4 and their ligands hyaluronan and HSP70, in modulating endothelial erosion in both haemodynamic environments.

## Conclusions

We demonstrated here that endothelial erosion from plaques frequently occurs in regions of elevated flow, which we previously demonstrated alters endothelial cell behaviour. Mimicking the inflammation and oxidative stress that might occur during exposure to cigarette smoking in a high flow environment strongly upregulated Nrf2 and this led to endothelial detachment that was not mediated by apoptosis. Furthermore, overexpression of Nrf2 upregulated genes OSGIN1 and OSGIN2 caused endothelial detachment that was associated with dysregulated HSP70/BAG3-mediated proteostasis. These findings highlight novel mechanisms that could be responsible for ACS and hence attractive for pharmacological intervention.

## Acknowledgments

The work was supported by British Heart Foundation grants (PG/11/44/28972, FS/12/77/29887 and CH95/001), Internal strategic funding from Manchester Metropolitan University, the National Health Research Institute (UK) Bristol Biomedical Research Unit in Cardiovascular Medicine and the European Commission through MOVE-AGE, an Erasmus Mundus Joint Doctorate program (2011-2015).

