## supplementary methods and data for "A pivotal role for Nrf2 in endothelial detachment-implications for endothelial erosion of stenotic plaques"

##### Arterial geometry reconstruction

The lumen geometries of LAD (left anterior descending artery), LCX (left circumflex artery) and RCA (right coronary artery) arteries (n=17) were reconstructed by combining the high lumen profile accuracy extracted from optical coherence tomography imaging with the vessel curvature obtained from bi-plane angiography.

##### Data acquisition

17 coronary arteries were studied [LAD, n = 7; LCX, n = 5; RCA, n =5] derived from 17 patients who underwent bi-plane angiography and intravascular optical coherence tomography (OCT) prior to medical intervention. For OCT imaging, a catheter was inserted into the affected patient coronary artery and OCT frames of a portion of that artery were obtained. Two institutes contributed data to this study. 8 sets of patient data from University Hospitals Bristol, UK and 9 from Policlinico A. Gemelli (Italy).

##### Lumen geometry reconstruction

The geometry was constructed by systematically going through 5 stages.

1. **3D centreline extraction.** The medical imaging software QAngio XA 3D by Medis (Leiden, Netherlands) was used to export the bi-plane angiography imaging data into a solid geometry. The solid geometry was then imported into Vascular Modelling Toolkit (VMTK) where the Cartesian co-ordinates of the artery centreline were extracted.

2. **OCT images segmentation.** The Cartesian co-ordinates of the OCT frames were exported from QCU-CMS by Medis (Leiden, Netherlands) into a large array of data. This array of data was then processed and used to build geometrical representations of each frame in SOLIDWORKS (SOLIDWORKS; Dassault Systèmes SOLIDWORKS Corp., Waltham, MA). The circumference of each frame was smoothened with a tolerance of 0.04 mm to eliminate un-physiological lumps present in the OCT data.

3. **Registration.** Manually interpreting the bi-plane angiography images enabled side branch centreline positions to be defined and built into the geometrical model. OCT frames were then stacked perpendicularly with the lumen centroid of each frame intersecting the 3D centreline [1-3]. The distance between the OCT frames was defined by the catheter pullback speed [4]. This stacking procedure is similar to the ANGUS technique used in similar studies in the literature [4-7]. A sampling rate of approximately 1 to 4 OCT images was applied where there is no stenosis, whilst for the stenosis region, the sampling rate was approximately 1 to 2, similar to methods used in the literature [3]. OCT frames at branch bifurcation sites were then rotationally orientated by aligning landmarks on the OCT frames that indicate a bifurcation site with the peripheral branches. The interim frames were then orientated using an interpolation technique. Both the registration and subsequent processes were conducted using SOLIDWORKS (SOLIDWORKS; Dassault Systèmes SOLIDWORKS Corp., Waltham, MA).

4. **Surfacing.** A lofting technique was then used between each OCT frame sequentially to create the final lumen surface. Peripheral branch diameters were extracted from the relative OCT frames and the branch walls were extruded in the model with fillets of 0.25 mm in radius applied at the bifurcation sites. Flow extensions were affixed to the inlet (one diameter in length) and outlets (seven diameters in length) to ensure physiologically accurate flow profiles within the artery [6, 8, 9] as shown by the blue surfaces in Figures S3 & S4. Inlet and outlet diameters for each case are shown in Table S1.

| | Diameter (mm) | | | | | | | |
|---|---|---|---|---|---|---|---|---|
| | Inlet | Outlet 1 | Outlet 2 | Outlet 3 | Outlet 4 | Outlet 5 | Outlet 6 | Outlet 7 |
| Case 1 | 3.48 | 1.6 | 1 | 1.6 | 0.2 | 2.3 | - | - |
| Case 2 | 2.76 | 0.6 | 0.9 | 0.8 | 0.4 | 1.1 | 1.3 | 2.05 |
| Case 3 | 2.24 | 2.2 | 0.9 | 0.9 | 1.5 | 2.87 | - | - |
| Case 4 | 2.41 | 0.8 | 1 | 0.7 | 1.61 | - | - | - |
| Case 5 | 3.66 | 2 | 0.5 | 0.8 | 0.7 | 2.91 | - | - |
| Case 6 | 4.55 | 2.3 | 0.4 | 0.2 | 1.7 | 1.6 | 0.9 | 2.77 |
| Case 7 | 3.9 | 1.7 | 1.5 | 0.6 | 1.2 | 0.7 | 3.1 | - |
| Case 8 | 3.66 | 1 | 1 | 0.5 | 2.7 | - | - | - |
| Case 9 | 3.79 | 1.6 | 0.7 | 1.8 | 1.6 | 0.8 | 2.79 | - |
| Case 10 | 3.71 | 0.7 | 0.8 | 0.3 | 1 | 2.6 | - | - |
| Case 11 | 3.34 | 1.2 | 0.9 | 1.9 | 1.6 | 3.02 | - | - |
| Case 12 | 4.04 | 1.7 | 1 | 1.5 | 2.46 | - | - | - |
| Case 13 | 3.33 | 0.6 | 0.7 | 1.2 | 1.1 | 2.26 | - | - |
| Case 14 | 4.75 | 3.8 | 2.1 | 1.9 | 2.4 | 2.89 | - | - |
| Case 15 | 3.32 | 1.4 | 0.6 | 0.8 | 1.4 | 1.67 | - | - |
| Case 16 | 3.31 | 1.6 | 1.3 | 1.7 | 0.7 | 2.36 | - | - |
| Case 17 | 3.67 | 1.6 | 1.4 | 1.6 | 0.7 | 2.61 | - | - |

Table S1 – Inlet & outlet diameters of the coronary artery lumen geometries used in this study. Outlet numbers ascend in order as they become more distal from the inlet

5. **Adhered thrombus identification.** The adhered thrombus sites in each OCT frame was noted sequentially and transferred onto the lumen surface. This was used to define where the adhered thrombus has formed on the lumen wall and was later used to extract haemodynamic metrics associated with atherosclerotic plaque erosion at this site. The thrombus location sites are highlighted in green in Figure S4. For information regarding the severity of stenosis for each case, refer to Table S2.

| | Non-diseased | Under thrombus | Percentage Stenosis | |
|---|---|---|---|---|
| | Average diameter (mm) | | Diameter (%) | Area (%) |
| Case 1 | 2.9 | 1.4 | 51.9 | 76.9 |
| Case 2 | 2.9 | 2.3 | 20.7 | 37.2 |
| Case 3 | 3.1 | 2.1 | 32.4 | 54.2 |
| Case 4 | 2.8 | 2.0 | 28.2 | 48.4 |
| Case 5 | 3.3 | 2.3 | 30.0 | 51.0 |
| Case 6 | 3.6 | 2.5 | 29.0 | 49.6 |
| Case 7 | 3.9 | 1.3 | 66.2 | 88.6 |
| Case 8 | 3.7 | 1.4 | 62.3 | 85.8 |
| Case 9 | 2.3 | 1.6 | 30.9 | 52.3 |
| Case 10 | 3.7 | 1.4 | 61.6 | 85.3 |
| Case 11 | 3.3 | 2.3 | 30.9 | 52.3 |
| Case 12 | 3.2 | 1.9 | 41.1 | 65.3 |
| Case 13 | 2.7 | 1.2 | 57.4 | 81.9 |
| Case 14 | 2.9 | 2.6 | 9.00 | 17.2 |
| Case 15 | 2.5 | 1.4 | 43.3 | 67.8 |
| Case 16 | 2.6 | 2.2 | 16.3 | 30.0 |
| Case 17 | 2.7 | 3.6 | -32.8 | -76.5 |
| Minimum | 2.3 | 1.2 | -32.8 | -76.5 |
| Quartile 1 | 2.7 | 1.4 | 28.2 | 48.4 |
| Median | 2.9 | 2.0 | 30.9 | 52.3 |
| Quartile 3 | 3.4 | 2.3 | 46.8 | 71.3 |
| Maximum | 3.9 | 3.6 | 66.2 | 88.6 |

Table S2 – Average diameter of areas of interest and the percentage of the stenosis.

**Omitted data.** 3 cases were omitted from the initial study of 20 patients, (10 from Bristol, 10 from Italy), due to insufficient data available for reconstructing the geometries.
Omitted data 1 (Italy): No OCT frames were generated proximal to the thrombus site. This is the most influential data that affects the flow local to the adhered thrombus site.
Omitted data 2 (BHI): Location and shape of the lesion could not be accurately identified in the OCT images as the catheter mostly obstructs the view of the lesion site.
Omitted data 3 (BHI): OCT data only covered the cannula wall and did not capture the coronary lumen profile.

**Computational Domain**

In order to ensure the accuracy of the simulations, a series of 10 steady-state computations with various mesh refinement levels (ranging from 0.7 to 14.5 million elements) were conducted to test mesh convergence (see Figure S1). Case 9 was used for the mesh convergency study as it was deemed to represent the general population of the cases studied by including common characteristics, such as the adhered thrombus site being in close proximity to a stenosis, having a relatively large number of outlets and the stenosis was a sufficient distance distal from the inlet to allow for flow to properly develop. The geometries were meshed using ANSYS-Meshing (Version 19.0). The mesh was based on a finite volume hybrid mesh consisting of tetrahedral elements within the core region and prism layers (3 elements thick) near the wall to allow for large spatial velocity gradients. Velocity representing flow of 5 cc/s was applied to the inlet based on the maximum velocity from the pulsatile waveform of the LCX artery of a 36 year old male subject during light exercise [10], as shown in Figure S3. Flow ratios are then applied to the outlets, except for the most distal, which had a zero pressure condition applied, details are given in Table S3. The velocity within the stenosis, as well as the overall volume were monitored. The maximum change in velocity was 0.5 % from 5.5 to 6.3 million elements. Velocity iso-surfaces as well as wall shear stress contours were qualitatively assessed. The mesh settings used in the 6.3 million elements study were deemed adequate for this application, and is therefore used for all subsequent simulations. The final mesh chosen for Case 9 is shown in Figure S2. It is worth noting that the mesh densities used in this work are approximately triple that used in a similar study by Timmins *et al.* [9], which further supports the confidence in the mesh settings used.

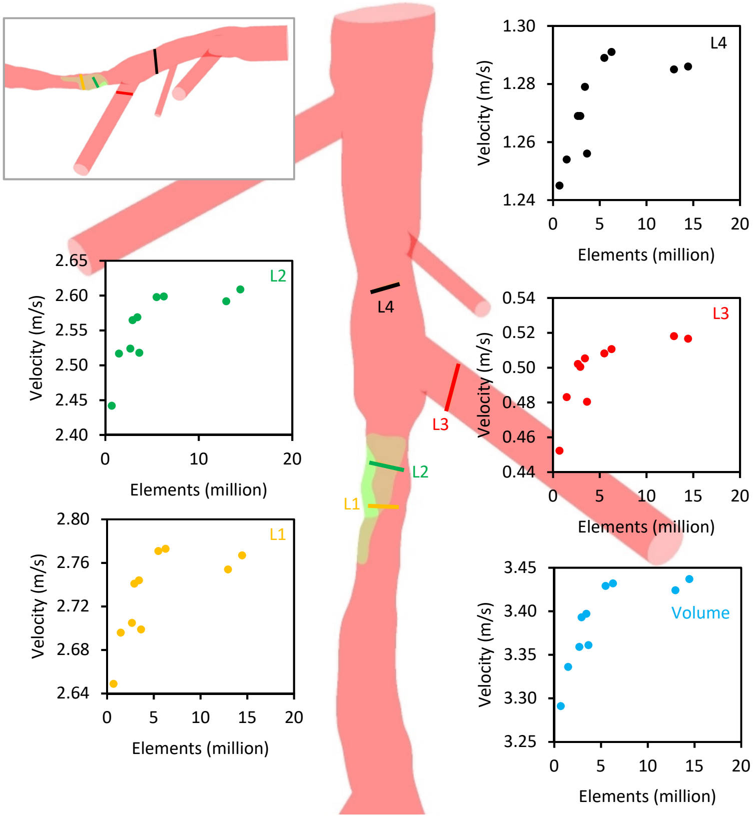

Figure S1 - Mesh convergence history plots. Velocity was monitored at 4 monitoring line locations, as well as the maximum volume velocity. Two lines (L1 & L2) are located within the stenosis to monitor the flow at the region of most interest. Monitoring line (L3) was placed within the branch proximal and in close proximity to the stenosis and adhered thrombus. Monitoring line (L4) is placed proximal to the aforementioned branch and within the LCX. These monitoring locations were chosen as they would capture the changes in the flow environment within the areas of most interest.

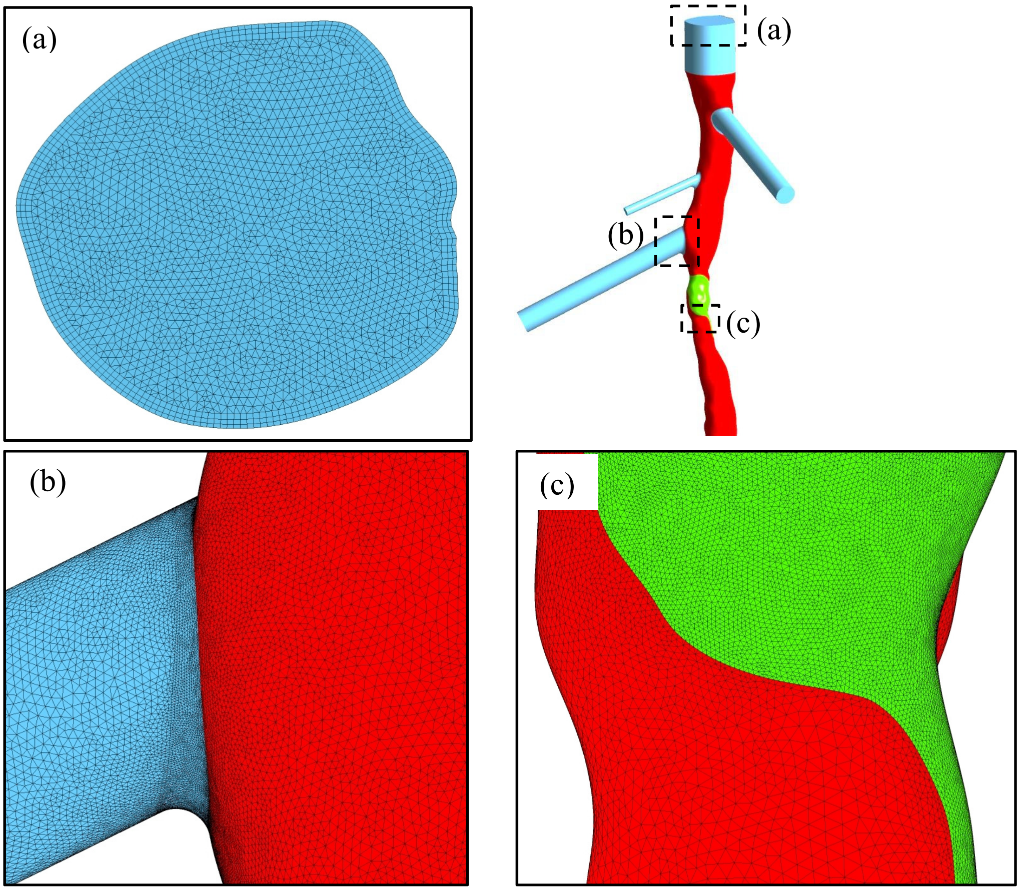

Figure S2 – Example of the chosen mesh setting used for all simulations. This mesh represents the optimal performing mesh taken from the mesh independency study (Case 9), in this case, the mesh has 6.3 million elements. (a) Cross-sectional view at the inlet, here the prism layers are visible on the lumen walls. (b) & (c) Refined face sizing (0.02 mm) at the bifurcation and adhered thrombus regions.

**Numerical Procedure**

The CFD solver ANSYS-CFX (Version 19.0) was used for the simulations. Blood was defined as a Newtonian and incompressible fluid with dynamic viscosity of 0.004 $Pa \cdot s^{-1}$ and density of 1060 $kg \cdot m^{-3}$ [11]. Blood flow through stenosed arteries is generally turbulent in nature [12], therefore, the shear stress transport model [13] was employed as the turbulence model in these simulations as it is considered the best suited turbulence model for capturing the turbulent transition in coronary arteries [12, 14, 15]. Simulations were run for 4 cardiac cycles (4 s) with a time step size of 1.25 ms and 10 coefficient loops per time step. This proved adequate in order to allow the boundary flow rate and pressure waveforms to converge, similar to previous studies in the literature [16]. The results were recorded during the final cardiac cycle.

**Boundary Conditions**

For all cases, no-slip boundary conditions were applied to all walls [17]. A rigid wall model was also assumed, which has been shown to be a valid assumption [18]. For every case, the respective time-dependent coronary velocity profiles (see Figure S3) were prescribed at the inlet based on a 36 year old healthy male subject, obtained from Kim et al. [10]. The inlet flow rates were assumed to remain constant for each coronary artery group. This was an assumption made due to a lack of patient specific flow data, that the coronary flow rate is unaffected by age, gender and body mass. At all the outlets, except for the most distal, an outlet velocity was prescribed. These outlet velocities were scaled versions of the inlet profile to satisfy Doriot's fit (Eq. 1).

$$\dot{m} = d^{2.27} \quad (1)$$

where $\dot{m}$ is the mass flow rate and $d$ is the vessel diameter. At the remaining, most distal, outflow tract, a traction free boundary condition was applied. This method is an accurate way of prescribing outlet conditions within coronary arteries [17, 19]. The velocities at the inlet and outlets were assigned as a velocity normal to the boundary. The boundary conditions for each case are summarised in Table S3.

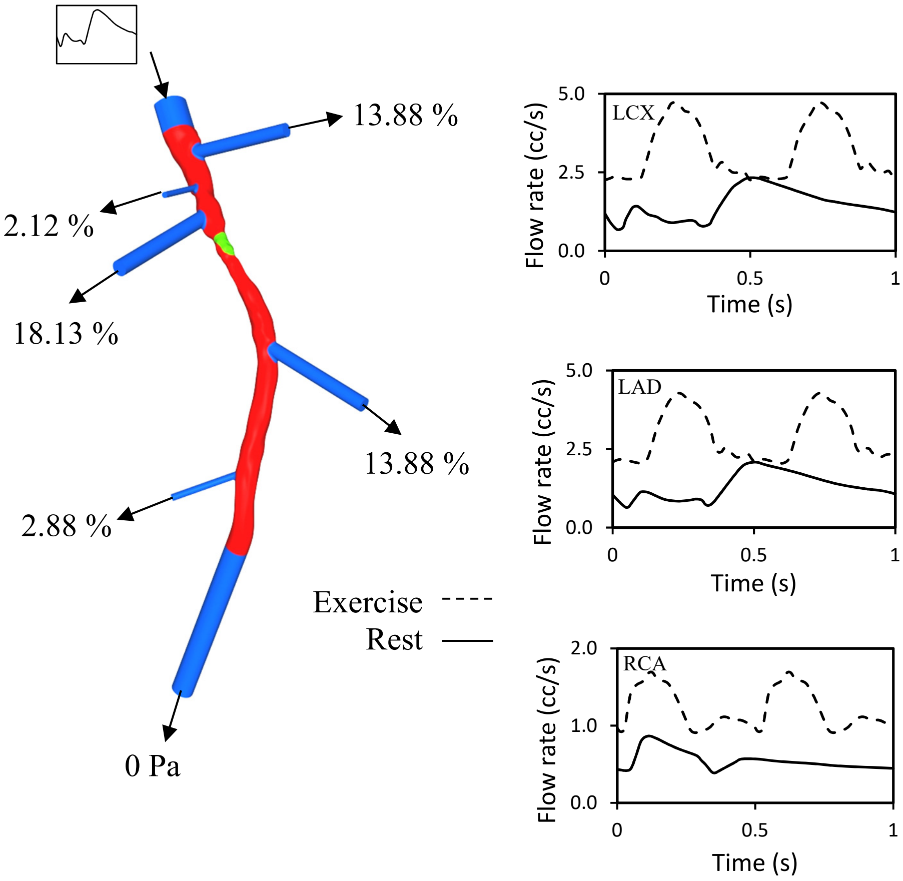

Figure S3 - Boundary conditions applied at the inlet and outlets. The LAD, LCX and RCA flow profiles were taken from Kim et al. [10] and used for the simulations. These profiles were from a patient at both rest and light exercise. Case 9 is used here as an example.

| | Boundary condition | | | | | | | |
|---|---|---|---|---|---|---|---|---|
| | Inlet | Outlet 1 | Outlet 2 | Outlet 3 | Outlet 4 | Outlet 5 | Outlet 6 | Outlet 7 |
| Case 1 | $\dot{m}_{RCA}(t)$ | 21.60 % | 7.43 % | 21.60 % | 0.19 % | 0 Pa | - | - |
| Case 2 | $\dot{m}_{LAD}(t)$ | 3.14 % | 7.88 % | 6.03 % | 1.25 % | 12.43 % | 18.16 % | 0 Pa |
| Case 3 | $\dot{m}_{LAD}(t)$ | 28.46 % | 3.74 % | 3.74 % | 11.93 % | 0 Pa | - | - |
| Case 4 | $\dot{m}_{LAD}(t)$ | 12.08 % | 20.04 % | 8.92 % | 0 Pa | - | - | - |
| Case 5 | $\dot{m}_{RCA}(t)$ | 27.75 % | 1.19 % | 3.47 % | 2.56 % | 0 Pa | - | - |
| Case 6 | $\dot{m}_{LAD}(t)$ | 27.70 % | 0.52 % | 0.11 % | 13.95 % | 12.15 % | 3.29 % | 0 Pa |
| Case 7 | $\dot{m}_{RCA}(t)$ | 15.76 % | 11.86 % | 1.48 % | 7.15 % | 2.10 % | 0 Pa | - |
| Case 8 | $\dot{m}_{LCX}(t)$ | 8.52 % | 8.52 % | 1.77 % | 0 Pa | - | - | - |
| Case 9 | $\dot{m}_{LCX}(t)$ | 13.88 % | 2.12 % | 18.13 % | 13.88 % | 2.88 % | 0 Pa | - |
| Case 10 | $\dot{m}_{LCX}(t)$ | 4.10 % | 5.55 % | 0.60 % | 9.21 % | 0 Pa | - | - |
| Case 11 | $\dot{m}_{RCA}(t)$ | 6.94 % | 3.61 % | 19.70 % | 13.33 % | 0 Pa | - | - |
| Case 12 | $\dot{m}_{LAD}(t)$ | 22.86 % | 6.85 % | 17.21 % | 0 Pa | - | - | - |
| Case 13 | $\dot{m}_{LCX}(t)$ | 3.17 % | 4.50 % | 15.28 % | 12.54 % | 0 Pa | - | - |
| Case 14 | $\dot{m}_{LAD}(t)$ | 42.46 % | 11.05 % | 8.80 % | 14.96 % | 0 Pa | - | - |
| Case 15 | $\dot{m}_{LCX}(t)$ | 25.48 % | 3.72 % | 7.15 % | 25.48 % | 0 Pa | - | - |
| Case 16 | $\dot{m}_{LAD}(t)$ | 18.70 % | 11.67 % | 21.46 % | 2.86 % | 0 Pa | - | - |
| Case 17 | $\dot{m}_{RCA}(t)$ | 16.87 % | 12.46 % | 16.87 % | 2.58 % | 0 Pa | - | - |

$\dot{m}_{LAD}(t)$, $\dot{m}_{LCX}(t)$ and $\dot{m}_{RCA}(t)$ represent time-dependent mass flow rates of LAD, LCX and RCA arteries respectively. Percentage values represent outflow ratios. The most distal outlet has a zero pressure condition. Outlet numbers ascend in order as they become more distal from the inlet

Table S2 – Summary of boundary conditions applied for each case. For all the outlets except the most distal, the outflow ratio was calculated according to Doriot's fit (Eq. 1).

#### Haemodynamic Metrics

The commercial visualisation tool, Ensight 10.2.3 was used to post-process the results and extract widely used wall shear-based haemodynamic metrics, specifically, Time-Averaged Wall Shear Stress (TAWSS) [20, 21],Oscillatory Shear Index (OSI) [21, 22], Relative Residence Time (RRT) [20, 22, 23] and Time-Averaged Wall Shear Stress Gradient (TAWSSG) [22, 24] according to equations (Eq. 2-5).

$$TAWSS = \frac{1}{T}\int_0^T |\vec{\tau}_w|\, dt \tag{2}$$

$$OSI = \frac{1}{2}\left(1 - \frac{\left|\int_0^T \vec{\tau}_w\, dt\right|}{\int_0^T |\vec{\tau}_w|\, dt}\right) \tag{3}$$

$$RRT = \frac{1}{\frac{1}{T}\left|\int_0^T \vec{\tau}_w\, dt\right|} \tag{4}$$

$$TAWSSG = \frac{1}{T}\int_0^T \sqrt{\left(\frac{\partial \tau_x}{\partial x}\right)^2 + \left(\frac{\partial \tau_y}{\partial y}\right)^2 + \left(\frac{\partial \tau_z}{\partial z}\right)^2}\; dt \tag{5}$$

In the above equations, $\vec{\tau}_w$ is the WSS vector and $T$ is the time period of the flow cycle.

**Tissue culture flow apparatus:**

**Surface coating**
Glass slides were coated with 0.1% gelatin (Gibco, Life Technologies) for >24hr and stored at 4°C.

**Flow System**
HCAECs were seeded on 0.1% gelatin coated slides with a density of 2.5 x $10^5$ and cultured for a minimum of 3 days to ensure complete confluency, production and reorganisation of sub-cellular matrix and maturation of cell-cell junctions. Culture under flow was performed using a parallel plate flow apparatus as described [25-27]. Briefly, confluent monolayers of HCAEC were initially cultured for 24 hours under oscillatory (0±5 dynes/cm$^2$, OSS), normal laminar (15 dynes/cm$^2$, LSS), or elevated laminar shear stress (75 dynes/cm$^2$, ESS) to allow cells to adapt to their shear environment. Three treatments of vehicle (control), TNFα (5ng/ml), CSE (10%), or both (T+C) were administered 16 hours apart using a parallel plate flow apparatus. HCAECs were cultured for a total of 72 hours, with 48 hours of treatment.

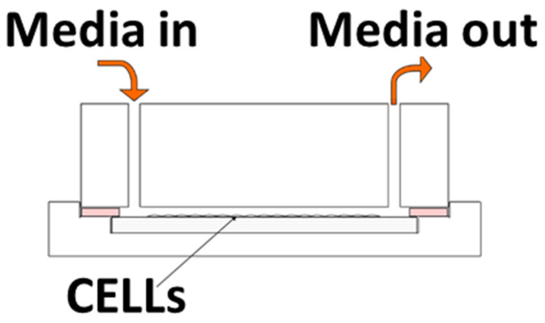

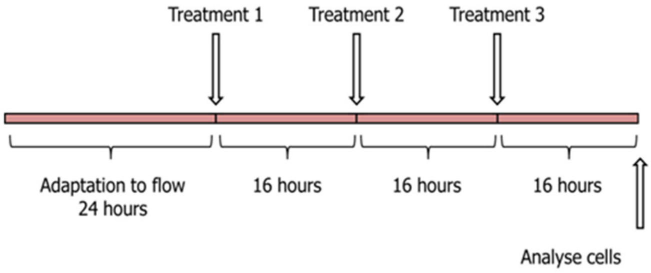

In addition, the orbital shaker model was also used [28], as this allows analysis of cells that have detached. For this system, cells were seeded on day 1 into 6 well plates, transduced with virus day 2 at a total pfu/cell of 400. For single gene transduction, this consisted of 200 pfu/cell active vector (AdOSGIN1, AdOSGIN2 or AdNrf2) and 200pfu/cell of control virus; for duel gene overexpression this consisted of 200pfu/cell of both vectors; 400pfu/cell of Adcontrol –E1/E3 'empty' Ad vector, was used as the control. Virus was removed on day 3 and the cells exposed to shear stress for 48 hours prior to analysis. 3 ml of MV2 fresh media was added and the plates were placed on an orbital shaker at 210 rpm for 48 hours to generate a defined shear stress pattern across the well, with low shear stress of approximately 4 dyn/cm2 in the centre and high shear stress of approximately 13 dyn/cm2 in the periphery. Compounds (Table S4) were added to the HCAECs for 48hrs on commencement of exposure to shear stress.

| **Compounds** | **Work concentration** | **Brand** | **Dissolved in** |
|---|---|---|---|
| Chloroquine | 150 or 300µM | Sigma | MV2 media |
| Bafilomycin | 50nM or 100nM | Sigma | DMSO |
| Metformin | 100µM | Sigma | MV2 media |
| VER-155008 | 15µM | Sigma | DMSO |

Table S4 Chemical compounds added during orbital shaker experiment

**Construction of Adenoviral vectors**

OSGIN1 and OSGIN2 were cloned into pCpG-free MCS (Invivogen). The coding regions were amplified by PCR using primers detailed in Table S8 below using KOD proofreading DNA-polymerase, introducing a 5' BglII site and 3' NheI site. Once inserted into the respective sites in pC-G-free MCS, the whole expression cassette was shuttled into pDC511 (Microbix Biosystems, Canada) for adenoviral vector production as previously described [26].

**Cytotoxicity, viability and apoptosis analysis**

Apotox-Glo triplex assay (Promega, Madison, USA) was performed in the HCAECs line, according to manufacturer's instructions, and 40 hours after AdOSGIN1 and AdOSGIN2 infection. To this end, cells were plated into 96-well plates at a density of 5 x $10^4$ cells per well. Results were normalized to DMSO treated control cells. Experiments were performed five times with eight replicates per condition.

**Caspase-3 activity assay**

Caspase-3 activity was measured using a caspase-3 activity assay kit (Promokine kit). This assay is based on a fluorometric immunosorbent enzyme assay performed in a multititer plate, according to the manufacturer's instructions. Briefly, a 96-well multititer plate was coated with human caspase-3 monoclonal antibodies and blocked to prevent nonspecific binding. Then, cellular lysates were incubated in the coated wells for 1.5 h at 37°C. After a washing step, the caspase-3-specific substrate Ac-Asp-Glu-Val-Asp-7-amino-4-(trifluoromethyl) coumarin (DEVD-AFC) was added and cleaved by the active enzymes to generate free fluorescent AFC. Fluorescence was measured at λ max = 505 nm after 3 h of incubation with the substrate and normalized by the protein content of each sample. Each treatment was done in 5 wells.

**Bicinchoninic-assay Assay kit**

Total protein of HCAECs was measured via bicinchoninic acid-assay (BCA), according to the manufacturer's instructions, using Pierce BCA Protein Assay (Thermo Scientific, Rockford, USA).

**Western blotting**

40 hr after adenovirus vector infection, cells were lysed in SDS lysis buffer [2% SDS; 50 mM Tris pH 6.8; 10% glycerol]. Protein concentration in the lysate was quantified using BCA assay kit. Between 12 and 20 µg of protein were loaded per lane on denaturing SDS–polyacrylamide gels and blotted onto Nitrocellulose or PVDF membranes. See Table S5 for further information. All experiments were conducted in triplicate.

| | Primary ab | Blocking Solution | Secondary ab |
|---|---|---|---|
| OSGIN1<br>(Biorbyt orb100666) | Rabbit-antiOSGIN1 (1:1000)<br>O/N 4°C | 1% BSA | Monoclonal Secondary HRP-anti rabbit (1:1000-1h) in TBSTween |
| OSGIN2<br>(Biorbyt orb185683) | Rabbit-antiOSGIN2 (1:500)<br>O/N 4°C | 1% BSA | Monoclonal Secondary HRP-anti rabbit (1:1000-1h) in TBSTween |
| PARP cleavage<br>(Cell Signaling D64E10) | Rabbit anti-PARP (1:1000)<br>O/N 4°C | 0,5% BSA | Monoclonal Secondary HRP-anti rabbit (1:2000-1h) in TBSTween |
| SQSTM1/p62<br>(Abcam ab56416) | Mouse anti-p62 (1:1000)<br>O/N 4°C | 3% Milk | Monoclonal Secondary HRP-anti mouse (1:5000-1h) in TBSTween |
| HSP70<br>(Abcam ab45133) | Rabbit anti-HSP70 (1:1000)<br>O/N 4°C | 5% Milk | Monoclonal Secondary HRP-anti rabbit (1:5000-1h) in TBSTween |

| | | | |
|---|---|---|---|
| LAMP1 (Abcam ab24170) | Rabbit anti-LAMP1 (1:1000) O/N 4°C | 1,5% Milk | Polyclonal Secondary HRP-anti rabbit (1:500-1h) + Monoclonal Secondary HRP-anti rabbit (1:500-1h) in TBSTween |

Table S5. Primary Antibodies, dilutions and methodology used in Western Blotting

**Bromo-deoxyuridine (BrdU) incorporation assay**

16 hr later on the morning of day 5, BrdU (10 µM, Sigma) was added to the media and the cells were left for a further 4 hr. Cells were washed (ice-cold PBS) and fixed in 70% ethanol. The incorporated BrdU was detected by immunocytochemistry using a mouse anti-BrdU primary antibody (1/500, Sigma B8434), biotinylated goat anti-mouse secondary (1/250, Sigma) and Extravidin-HRP (1/250, Sigma E2886), and visualised with diaminobenzidine staining. Counterstaining was made by hematoxylin (Mayer's Hematoxylin Sigma) for 2 minutes. Multiple images of the wells were taken in the centre of each well for assessment of BrdU staining. The percentage of cells positive for BrdU staining was calculated.

**Immunocytochemistry**

HCAECs were exposed to AdOSGIN1 and AdOSGIN2 infection before being fixed in cold 4% paraformaldehyde for 10 minutes. Cells were permeabilised with 0.1% triton, blocked in 20% goat serum and probed with rabbit anti-OSGIN1 (1:100, Biorbyt) and rabbit anti-OSGIN2 (1:75, Biorbyt) , and rabbit anti-VE-Cadherin (Cell Signaling) followed by goat anti rabbit Alexa Fluor 488 (1/200, Invitrogen). Further staining were carried out with mouse anti-Vinculin (1/400, Sigma), mouse anti-βcathenin (1/200, BD Transduction Laboratories), mouse -mab137 (1/100, Sigma) followed by goat anti mouse Alexa Fluor 594 (1/200, Invitrogen). In combination with the previous staining, anti-Phalloidin (1/250, Sigma), and anti-Tubulin (1/1000, Abcam) were added (antibody titration Table S6).

| **Antibody** | **Type** | **Dilution** | **Company** |
|---|---|---|---|
| OSGIN1 | Rabbit | 1:100 | (Biorbyt orb100666) |
| OSGIN2 | Rabbit | 1:75 | (Biorbyt orb185683) |
| VE-Cadherin | Rabbit | 1:400 | (Cell Signalling D87F2) |
| β-Catenin | Mouse | 1:100 | (BD Transduction Laboratories 610153) |
| Vinculin | Mouse | 1:400 | (Sigma V4505) |
| Mab113 | Mouse | 1:100 | (Abcam ab92824) |
| Tubulin | Already conjugated (Green) | 1:1000 | (Abcam ab64503) |
| Phalloidin | Already conjugated (Red) | 1:250 | (Sigma P1951) |
| HSP70 | Rabbit | 1:400 | (Abcam ab45133) |
| LAMP1 | Rabbit | 1:100 | (Abcam ab24170) |
| SQSTM1/p62 | mouse | 1:200 | (Abcam ab56416) |

Secondary Antibody description

| | | | |
|---|---|---|---|
| Alexa fluor488 | Anti mouse or rabbit | 1:200 | Invitrogen |
| Alexa fluor647 | Anti mouse or rabbit | 1:200 | Invitrogen |

Table S6. Primary Antibodies used in Immunocytochemical analysis

**Immunofluorescence on Mouse Aortas**
8µM frozen sections of aortas from mice exposed to cigarette smoke (as described[29]) for 3 months, or control mice, were fixed in ice cold acetone before immunofluorescence was performed using the antibodies described above.

**Flow cytometric analysis**
For cell cycle analysis, cells ($1\times10^5$) were washed twice with ice-cold PBS, treated with trypsin, and fixed in cold 70% ethanol at 4°C for at least 24 h, washed twice in PBS, and incubated in 25 µg/ml of RNase for 30 min at 37°C. Before analysis, cells were stained with 25 µg/ml of propidium iodide (PI) at room temperature for 30 min. Analyses were performed using a FACScan flow cytometer (FACS Calibur, BD Transduction Laboratories). Data obtained from the cell cycle distributions were analyzed using Modfit software.

**Senescent-associated β-galactosidase staining**
Paraformaldehyde fixed HCAECs were rinsed with ice-cold PBS pH 6.0 and submerged in senescent-associated β-galactosidase (SA-β-Gal) staining solution [1 mg/ml 5-bromo-4-chloro-3-indolyl β-d-galactopyranoside (X-gal), 5 mM $K_4Fe(CN)_6$, 5 mM $K_3Fe(CN)_6$, 150 mM $NaCl_2$ and 2 mM $MgCl_2$ titrated with 1M $NaH_2PO_4$ to pH 6.0] and incubated overnight at 37°C. After staining, HCAECs were rinsed with ice-cold PBS pH 6.0 pictures were taken with a Leica M165 FC stereomicroscope.

**RNA extraction and reverse transcriptase-polymerase chain reaction (RT-PCR)**
Total RNA was extracted using the RNeasy Mini Kit (Norgen) according to the manufacturer's protocol. 250ng RNA were reverse transcribed into cDNA with random primer by qiagen reverse transcriptase (RT) (Qiagen). 1µg cDNA product (relative to RNA amount) was amplified by standard PCR with Taq DNA polymerase (Sensifast, SybrGreen, LOW-ROX Kit, Bioline) and primers.

**Real-time PCR**
Primers were designed using NCBI software. For each gene, SYBR Green was used in place of a labelled probe (primers sequences Table 2). GAPDH RNA was used as the internal control for each gene and the target gene was amplified in duplex in PCR mixtures (10 µl final volume) containing 4 µl Sybr® Green PCR Master Mix, cDNA template 1 µl, optimised primers 2 µl and 3 µl of $H_2O$. PCR thermal cycle parameters were: 5 minutes 95°C 30 seconds between at 65°C to 70°C depending on primers optimisation, 30 seconds at 72°C 40 cycles of 95°C for 15 seconds and 60°C for 1 minute. Reactions were performed, and fluorescence was monitored in a Applied Biosystem detector (Applied Biosystem). Relative mRNA expression level was defined as the ratio of target gene expression level to GAPDH mRNA expression. Primers sequences can be found in Table S7.

| | |
|---|---|
| p21 (SW878F/879R) | F CTCAGGGTCGAAAACGGCGG<br>R GTGGGCGGATTAGGGCTTCCT |
| p16 (SW926F/927R) | F CGAGCTCGGCCCTGGAG<br>R TCGGGCGCTGCCCATCAT |
| HSPA1A (SW355F/356R) | F TGAGGAGCTGCTGCGACAGT<br>R GGCTGGAAACGGAACACTGG |
| HSPA1B (SW357F/358R) | F TGTTGAGTTTCCGGCGTTCC<br>R AACACCCCCACGCAGGAGTA |
| BAG3 (SW966F/967R ) | F GCGGGGCATGCCAGAAACCA<br>R CTGGCCGGGTAACGTTCTGCT |
| ATG7 (SW894F/895R) | F GGACTGGCCGTGATTGCAGGA<br>R ATCCGATCGTCACTGCTGCTGG |

| | |
|---|---|
| ATG9A (SW938F/939R) | F AGAGGCGCTACGGTGGCATC<br>R GCCTTGATGCCGACTGCCCA |
| MAP1LC3B (SW934F/935R) | F CGCCCAGATCCCTGCACCAT<br>R AGCATTGAGCTGTAAGCGCCTTCT |
| SQSTM1/p62 (SW936F/937R) | F GGACGGGGACTTGGTTGCCTT<br>R CGGGTTCCTACCACAGGCCC |
| GABARAPL1/ATG8 (SW940F/941R) | F CGGACAGGGTCCCCGTGATTG<br>R AGCACTGGTGGGAGGGATGGT |
| GAPDH (SW180F/181R) | F CGGATTTGGTCGTATTGGGCG<br>R GCCTTCTCCATGGTGGTGAAGAC |

Table S7 Primer sequences used for real-time PCR

**Statistical analysis**

Data are presented as means ± standard error of the mean (SEM) of 3 independent experiments. One-way ANOVA was used to determine the difference between three or more groups. Two-way ANOVA was used to evaluate 3 or more groups on different conditions. P-values <0.05 were considered to indicate statistically significant differences. Graph pad prism software has been used for statistical analysis (GraphPad Software, La Jolla, CA).

**RNASeq data analysis**

Strand-specific RNA-seq libraries were prepared using the Illumina workflow with the TruSeq® Stranded mRNA Sample Preparation Kit. Paired-end reads of 65bp were generated from each sample on the Illumina platform of HiSeq4000. The fastq files generated were analysed with FastQC [30], any low quality reads and contaminated barcodes were trimmed with Trimmomactic [31]. All libraries were aligned to the hg38 assembly of human genome using STAR-2.5.3a [32] and only the unique alignments were reported for each read. The mapped reads were also counted with STAR at gene level against gencode.v25.annotation.gtf. R was used for all the statistical analysis of data [33]. The counts data was normalized and donor effect removed using the R package RUVSeq [34]. Differentially expressed genes were detected with the R package of DESeq2 [35] between groups of experimental data sets. The cluster analysis was carried out on the DE genes identified with DESeq2 using a padj cut off of 0.05 with gplots [36].The predicted upstream regulators and altered canonical pathways were generated through the use of IPA (QIAGEN Inc., https://www.qiagenbioinformatics.com/products/ingenuity-pathway-analysis) [37].

**A pivotal role for Nrf2 in endothelial detachment– implications for endothelial erosion of stenotic plaques.** Sandro Satta, *et. al.*

### SUPPLEMENTARY DATA

* The median value for the respective metric.

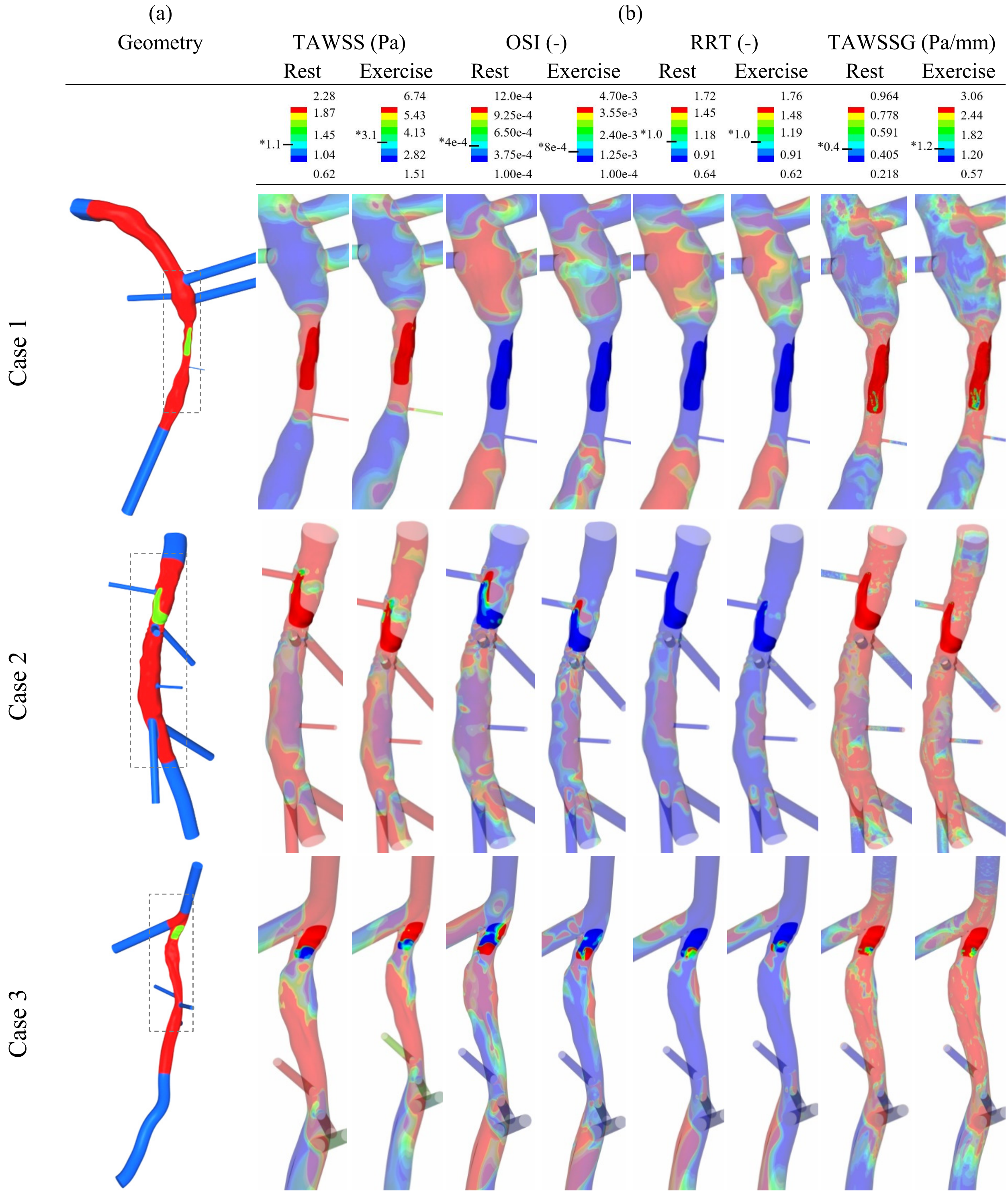

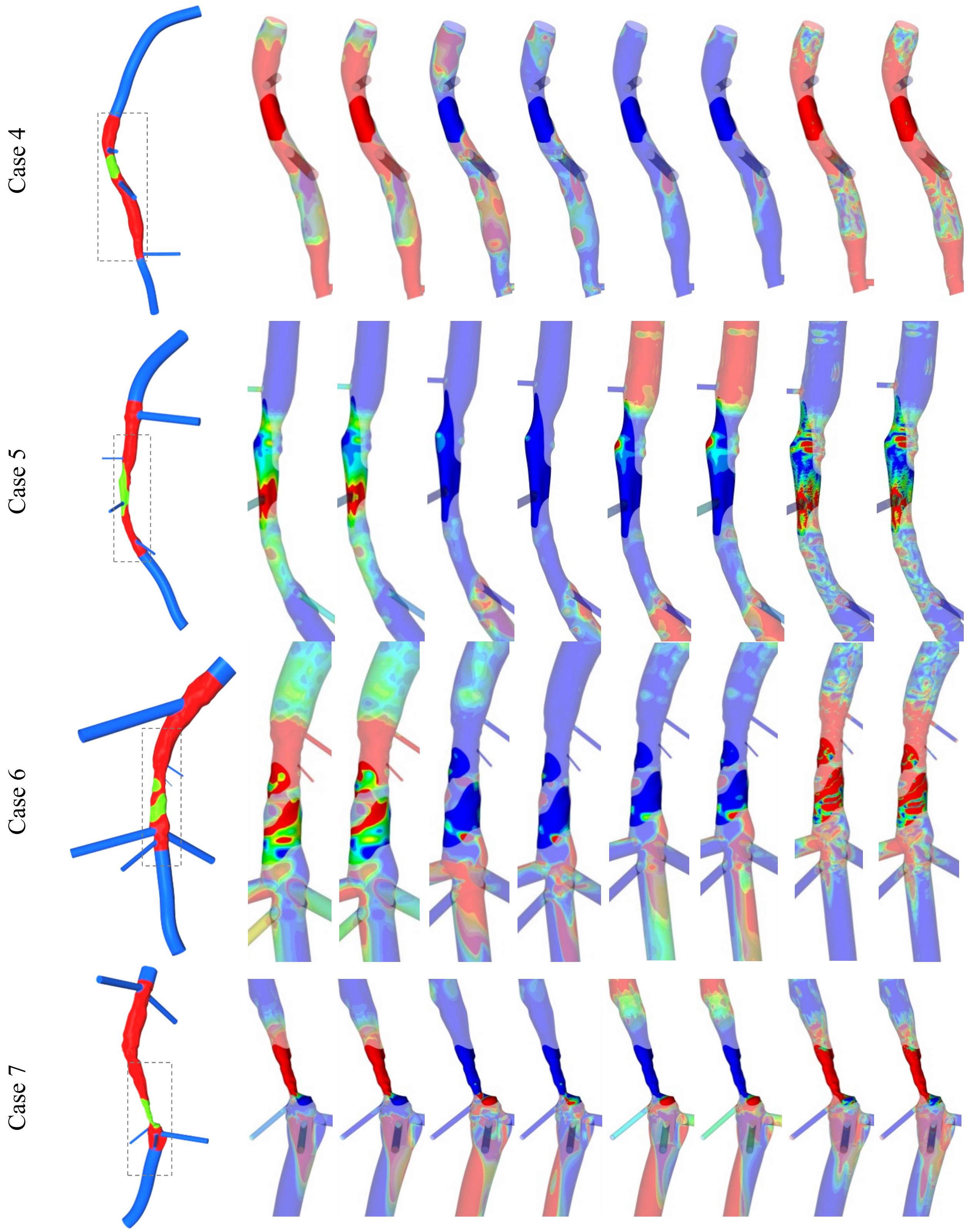
Case 4
Case 5
Case 6
Case 7

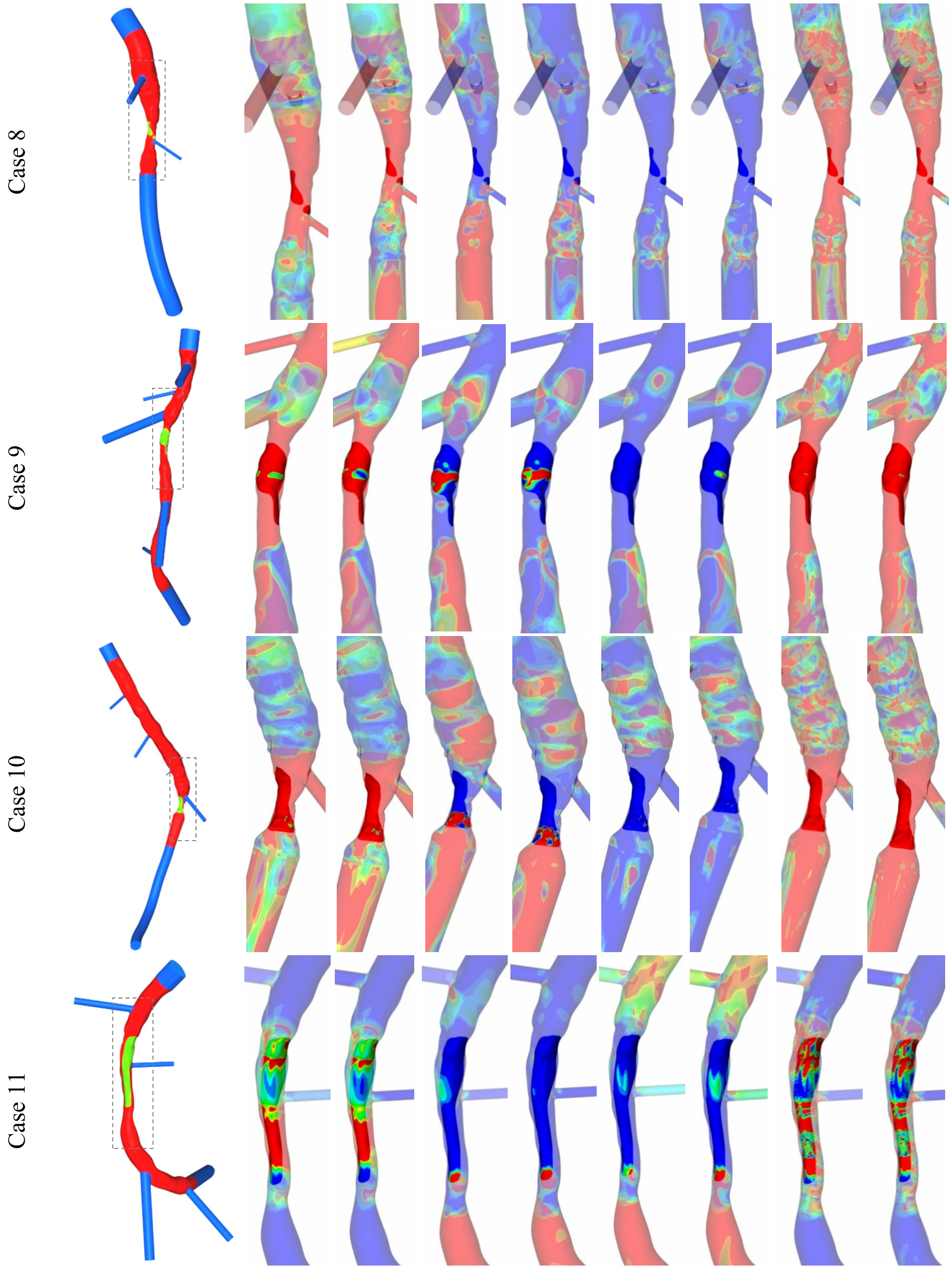
Case 8
Case 9
Case 10
Case 11

**A pivotal role for Nrf2 in endothelial detachment– implications for endothelial erosion of stenotic plaques.** Sandro Satta, *et. al.*

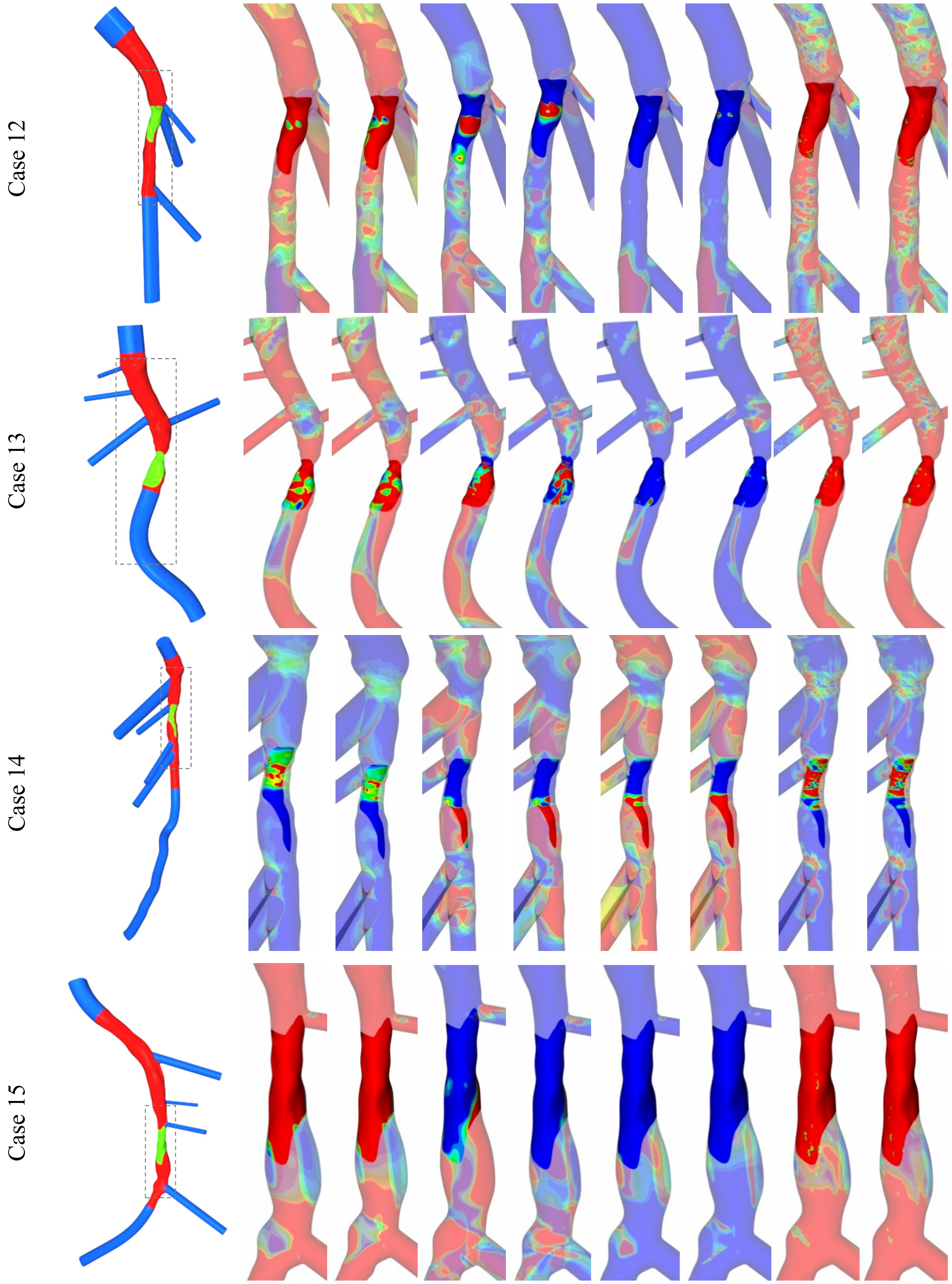

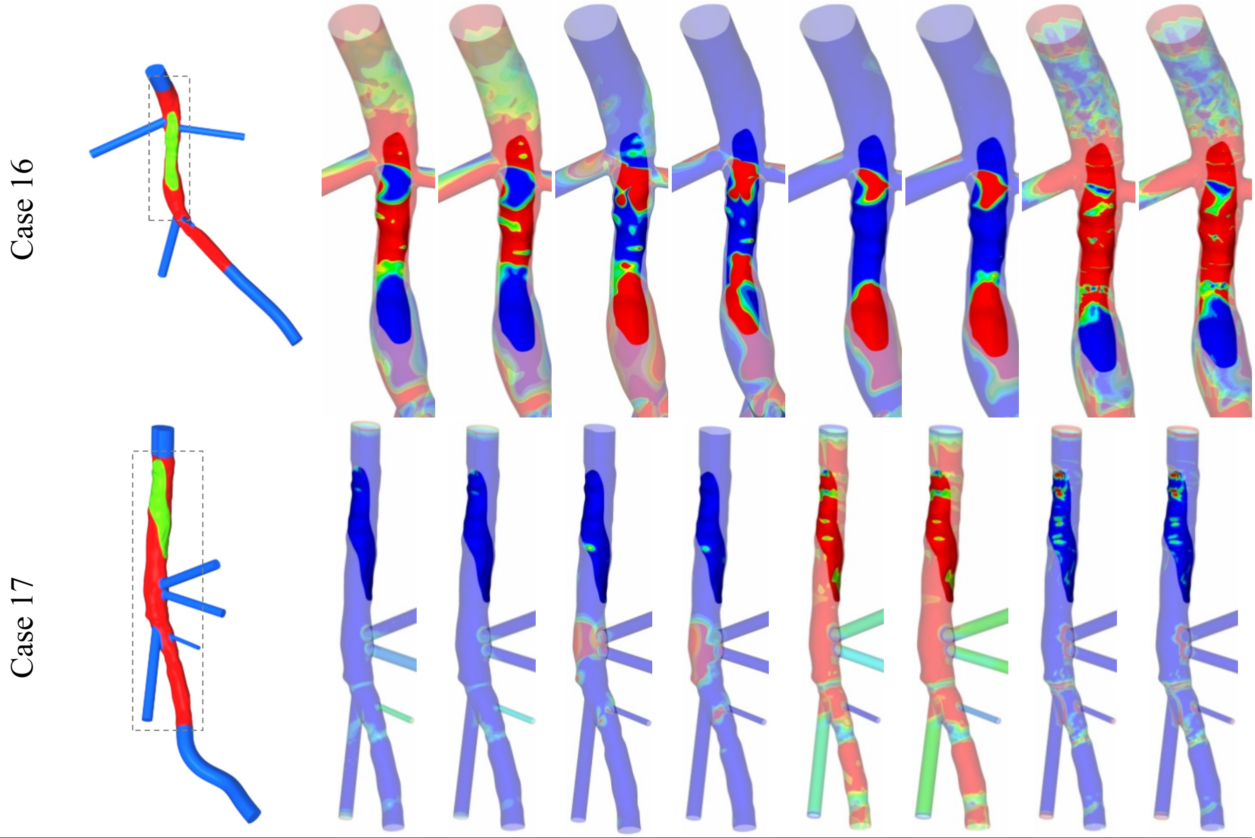

Figure S4. (a) Reconstructed lumen geometries of the LAD, LCX and RCA arteries. Red sections are reconstructed using a hybrid OCT/bi-plane angiography method to produce a lumen profile to a high accuracy. The blue sections represent the geometry reconstructed using bi-plane angiography. The adhered thrombus is defined by the green surface, extracted from OCT data. (b) Haemodynamic metrics extracted from CFD simulations. Time-Averaged Wall Shear Stress (TAWSS), Oscillatory Shear Index (OSI), Relative Residence Time (RRT) and Time-Averaged Wall Shear Stress Gradient (TAWSSG). Both 'rest' and 'exercise' flow rate conditions were simulated. The thrombus is the opaque portion of the metrics, whilst the remainder of the lumen is semi-transparent. Flow is from top to bottom for all images. Minimum and maximum values for the legends are the lower and upper quartiles of the respective metrics averaged across the rest and exercise cases separately, as shown in Table S4 & Table S5. RRT ranges are normalised in respect to the averaged median RRT at the 'non-diseased' location, with the median values being 1.11 and 0.43 for rest and exercise respectively.

**A pivotal role for Nrf2 in endothelial detachment– implications for endothelial erosion of stenotic plaques.** Sandro Satta, *et. al.*

| **Rest** | TAWSS (Pa) | | | TAWSSG (Pa/mm) | | $WSS_{Max}$ (Pa) | |
|---|---|---|---|---|---|---|---|
| | Non-diseased | Thrombus | Thrombus/ Non-diseased | Non-diseased | Thrombus | Non-diseased | Thrombus |
| Case 1 | 0.75 | 4.79 | 2.68 | 0.28 | 3.99 | 0.59 | 3.79 |
| Case 2 | 2.41 | 11.90 | 2.30 | 1.58 | 22.00 | 1.57 | 8.10 |
| Case 3 | 1.03 | 4.64 | 2.18 | 0.32 | 7.62 | 0.64 | 3.16 |
| Case 4 | 2.10 | 9.25 | 2.14 | 0.99 | 3.95 | 1.38 | 6.33 |
| Case 5 | 0.46 | 1.82 | 1.98 | 0.22 | 0.71 | 0.38 | 1.44 |
| Case 6 | 1.10 | 2.31 | 1.07 | 0.41 | 2.09 | 0.70 | 3.09 |
| Case 7 | 0.26 | 7.66 | 4.86 | 0.08 | 10.90 | 0.22 | 5.92 |
| Case 8 | 1.47 | 28.60 | 4.28 | 0.55 | 39.20 | 0.91 | 39.29 |
| Case 9 | 2.59 | 10.70 | 2.05 | 0.96 | 18.40 | 1.64 | 7.40 |
| Case 10 | 1.01 | 23.00 | 4.51 | 0.38 | 58.20 | 0.58 | 15.83 |
| Case 11 | 0.62 | 2.14 | 1.80 | 0.10 | 1.03 | 0.50 | 1.70 |
| Case 12 | 2.28 | 7.29 | 1.68 | 0.65 | 8.90 | 1.50 | 4.96 |
| Case 13 | 4.16 | 9.68 | 1.22 | 2.33 | 38.40 | 2.73 | 6.81 |
| Case 14 | 0.48 | 1.08 | 1.18 | 0.06 | 0.87 | 0.32 | 0.69 |
| Case 15 | 4.58 | 9.51 | 1.05 | 1.04 | 8.20 | 3.05 | 6.50 |
| Case 16 | 1.34 | 2.35 | 0.81 | 0.32 | 2.88 | 0.90 | 1.58 |
| Case 17 | 0.48 | 0.46 | -0.04 | 0.11 | 0.27 | 0.39 | 0.38 |
| Minimum | 0.26 | 0.46 | -0.04 | 0.06 | 0.27 | 0.22 | 0.38 |
| Quartile 1 | 0.62 | 2.31 | 1.18 | 0.22 | 2.09 | 0.49 | 1.70 |
| Median | 1.10 | 7.29 | 1.98 | 0.38 | 7.62 | 0.70 | 4.95 |
| Quartile 3 | 2.28 | 9.68 | 2.15 | 0.96 | 18.36 | 1.50 | 6.81 |
| Maximum | 4.58 | 28.57 | 4.86 | 2.33 | 58.22 | 3.05 | 39.29 |
| **Exercise** | TAWSS (Pa) | | | TAWSSG (Pa/mm) | | $WSS_{Max}$ (Pa) | |
| | Non-diseased | Thrombus | Thrombus/ Non-diseased | Non-diseased | Thrombus | Non-diseased | Thrombus |
| Case 1 | 1.93 | 12.00 | 2.65 | 0.87 | 10.40 | 5.27 | 9.98 |
| Case 2 | 7.89 | 33.40 | 2.08 | 5.93 | 69.40 | 1.40 | 1.59 |
| Case 3 | 3.11 | 12.30 | 1.99 | 1.16 | 21.90 | 13.40 | 54.50 |
| Case 4 | 6.74 | 25.80 | 1.94 | 3.77 | 12.70 | 9.92 | 32.54 |
| Case 5 | 1.08 | 4.59 | 2.10 | 0.57 | 1.93 | 11.44 | 41.51 |
| Case 6 | 2.93 | 5.63 | 0.94 | 1.24 | 6.62 | 0.86 | 30.72 |
| Case 7 | 0.59 | 20.40 | 5.11 | 0.21 | 32.30 | 19.00 | 43.90 |
| Case 8 | 3.85 | 76.50 | 4.31 | 1.65 | 118.00 | 4.73 | 8.61 |
| Case 9 | 8.35 | 27.30 | 1.71 | 4.13 | 55.20 | 2.25 | 7.94 |
| Case 10 | 2.34 | 62.60 | 4.74 | 1.03 | 171.00 | 5.25 | 20.30 |
| Case 11 | 1.51 | 5.33 | 1.82 | 0.29 | 2.96 | 0.59 | 3.79 |
| Case 12 | 6.16 | 20.00 | 1.70 | 1.92 | 28.90 | 15.18 | 44.03 |
| Case 13 | 11.20 | 28.10 | 1.33 | 7.20 | 119.00 | 1.57 | 6.85 |
| Case 14 | 1.03 | 2.57 | 1.31 | 0.20 | 2.34 | 3.63 | 104.42 |
| Case 15 | 11.80 | 26.20 | 1.15 | 3.06 | 26.00 | 1.58 | 4.04 |
| Case 16 | 3.33 | 6.14 | 0.88 | 0.96 | 9.02 | 6.25 | 125.07 |
| Case 17 | 1.00 | 1.07 | 0.11 | 0.28 | 0.71 | 18.27 | 48.17 |
| Minimum | 0.59 | 1.07 | 0.11 | 0.20 | 0.71 | 0.59 | 1.59 |
| Quartile 1 | 1.51 | 5.63 | 1.31 | 0.57 | 6.62 | 1.58 | 7.94 |
| Median | 3.11 | 19.96 | 1.82 | 1.16 | 21.86 | 5.25 | 30.72 |
| Quartile 3 | 6.74 | 27.29 | 2.02 | 3.06 | 55.19 | 11.44 | 44.03 |
| Maximum | 11.81 | 76.51 | 5.11 | 7.20 | 170.93 | 19.00 | 125.07 |

Table S8 - Area-averaged values of TAWSS, TAWSSG and $WSS_{Max}$ (maximum flow during the cardiac cycle) at the site defined as being normal flow (non-diseased) and at the adhered thrombus location.

| **Rest** | OSI (-) | | | RRT | |
|---|---|---|---|---|---|
| | Non-diseased | Thrombus | Thrombus/ Non-diseased | Non-diseased (1/Pa) | Thrombus (-) |
| Case 1 | 4.90E-04 | 7.76E-06 | -5.98 | 1.70E+00 | -8.74E-01 |
| Case 2 | 2.73E-02 | 1.09E-03 | -4.65 | 1.45E+00 | -8.71E-01 |
| Case 3 | 1.67E-02 | 1.15E-02 | -0.55 | 1.91E+00 | -7.02E-01 |
| Case 4 | 6.10E-03 | 6.40E-05 | -6.57 | 7.13E-01 | -8.31E-01 |
| Case 5 | 4.96E-05 | 3.84E-05 | -0.37 | 2.31E+00 | -6.99E-01 |
| Case 6 | 1.26E-03 | 1.59E-04 | -2.98 | 1.11E+00 | -5.35E-01 |
| Case 7 | 4.02E-04 | 1.29E-03 | 1.69 | 4.06E+00 | -8.10E-01 |
| Case 8 | 5.89E-04 | 2.29E-04 | -1.36 | 7.55E-01 | -9.26E-01 |
| Case 9 | 1.17E-03 | 1.69E-03 | 0.53 | 4.75E-01 | -6.42E-01 |
| Case 10 | 2.07E-04 | 1.03E-02 | 5.64 | 1.06E+00 | -8.82E-01 |
| Case 11 | 3.58E-05 | 1.64E-04 | 2.20 | 1.64E+00 | -6.45E-01 |
| Case 12 | 5.97E-05 | 1.22E-03 | 4.35 | 4.54E-01 | -5.46E-01 |
| Case 13 | 1.32E-04 | 9.79E-03 | 6.21 | 2.66E-01 | 7.28E-01 |
| Case 14 | 1.51E-04 | 1.71E-02 | 6.83 | 2.18E+00 | 3.66E-01 |
| Case 15 | 1.20E-05 | 1.32E-02 | 10.10 | 2.35E-01 | 6.25E-01 |
| Case 16 | 7.56E-04 | 3.07E-02 | 5.34 | 8.26E-01 | 1.58E+00 |
| Case 17 | 1.64E-05 | 6.24E-05 | 1.92 | 2.21E+00 | 9.14E-02 |
| Minimum | 0.00E+00 | 0.00E+00 | -6.57 | 2.40E-01 | -9.30E-01 |
| Quartile 1 | 1.00E-04 | 2.00E-04 | -1.36 | 7.10E-01 | -8.30E-01 |
| Median | 4.00E-04 | 1.20E-03 | 1.69 | 1.11E+00 | -6.50E-01 |
| Quartile 3 | 1.20E-03 | 1.03E-02 | 5.42 | 1.91E+00 | 9.00E-02 |
| Maximum | 2.73E-02 | 3.07E-02 | 10.10 | 4.06E+00 | 1.58E+00 |
| **Exercise** | OSI (-) | | | RRT | |
| | Non-diseased | Thrombus | Thrombus/ Non-diseased | Non-diseased (1/Pa) | Thrombus (-) |
| Case 1 | 7.84E-04 | 1.21E-05 | -6.02 | 7.57E-01 | -8.87E-01 |
| Case 2 | 1.10E-02 | 2.13E-03 | -2.37 | 3.81E-01 | -7.85E-01 |
| Case 3 | 8.18E-03 | 1.13E-02 | 0.47 | 5.56E-01 | -5.45E-01 |
| Case 4 | 4.71E-03 | 4.67E-05 | -6.66 | 2.62E-01 | -8.30E-01 |
| Case 5 | 1.41E-04 | 6.38E-05 | -1.15 | 1.03E+00 | -7.02E-01 |
| Case 6 | 9.21E-04 | 4.95E-04 | -0.90 | 4.26E-01 | -4.42E-01 |
| Case 7 | 1.38E-03 | 5.64E-04 | -1.29 | 1.95E+00 | -8.55E-01 |
| Case 8 | 2.84E-04 | 7.89E-03 | 4.79 | 3.07E-01 | -8.70E-01 |
| Case 9 | 3.55E-03 | 5.13E-03 | 0.53 | 1.66E-01 | -4.59E-01 |
| Case 10 | 2.28E-04 | 1.09E-02 | 5.57 | 4.82E-01 | -9.15E-01 |
| Case 11 | 4.15E-05 | 6.36E-04 | 3.94 | 6.73E-01 | -6.07E-01 |
| Case 12 | 2.97E-05 | 3.07E-03 | 6.69 | 1.68E-01 | -4.01E-01 |
| Case 13 | 8.23E-05 | 7.98E-03 | 6.60 | 1.03E-01 | 5.49E-01 |
| Case 14 | 1.28E-02 | 2.85E-02 | 1.16 | 1.33E+00 | 2.33E-01 |
| Case 15 | 8.35E-06 | 9.53E-03 | 10.16 | 9.59E-02 | 1.40E+00 |
| Case 16 | 4.82E-03 | 1.36E-02 | 1.49 | 3.78E-01 | 1.52E+00 |
| Case 17 | 4.46E-05 | 1.26E-04 | 1.50 | 1.11E+00 | -2.09E-02 |
| Minimum | 0.00E+00 | 0.00E+00 | -6.66 | 1.00E-01 | -9.10E-01 |
| Quartile 1 | 1.00E-04 | 5.00E-04 | -1.15 | 2.60E-01 | -8.30E-01 |
| Median | 8.00E-04 | 3.10E-03 | 1.16 | 4.30E-01 | -5.40E-01 |
| Quartile 3 | 4.70E-03 | 9.50E-03 | 4.99 | 7.60E-01 | -2.00E-02 |
| Maximum | 1.28E-02 | 2.85E-02 | 10.16 | 1.95E+00 | 1.52E+00 |

Table S9 - Area-averaged values of OSI and RRT at the site defined as being normal flow (non-diseased) and at the adhered thrombus location.

#### Limitations

The limited number of angiography images available of the coronary tree at various angles limits the accuracy of incorporating the side branches into the lumen geometries during the reconstruction process to an extent. A sensitivity test to quantify the effects the angles of the branches have on the haemodynamic metrics around the adhered thrombus location would provide insight into the impact of this limitation.

The reconstruction process presented in this work was semi-automated, requiring user input at various stages of the process. In order to make the reconstruction technique useful for processing a large cohort of data, the process should be designed such that it requires minor manual user input.

It is currently challenging to incorporate biochemical models into CFD simulations [38], therefore, the biochemical factors that are involved with cardiovascular diseases have been ignored in this work.

Due to limited patient-specific flow data, flow data of the RCA, LCA and LCX arteries for one patient were applied to all of the cases studied.

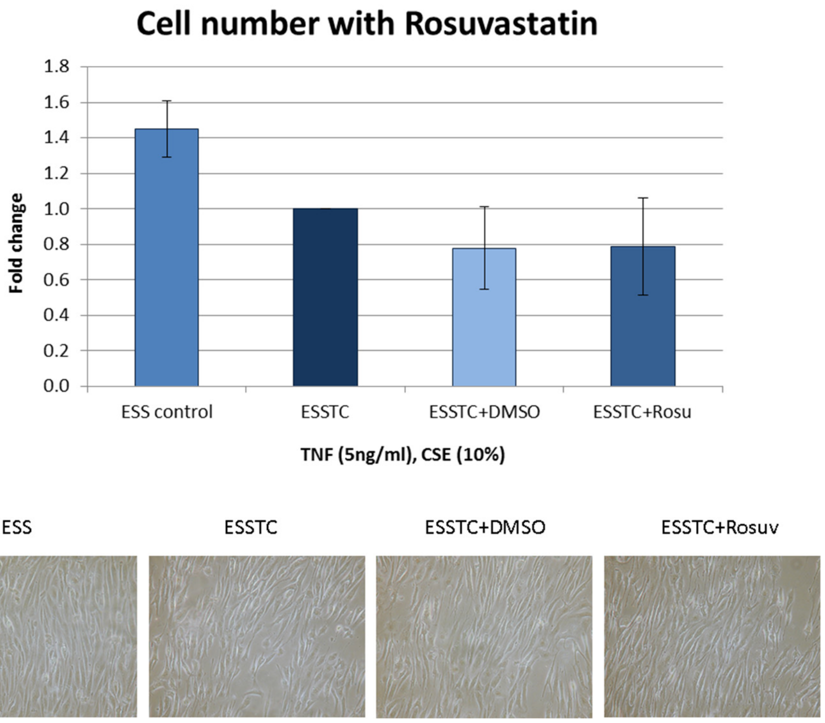

Figure S5: Cell Adhesion of HCAECs exposed to a combination of elevated shear stress, TNFα and CSE together (ESSTC), with additional 3μM Rosuvastatin treatment, vs untreated and DMSO controls at ESS for 72 hours. Cell number was quantified using picogreen assay, expressed as mean fold change against ESSTC ± S.E. n=3.

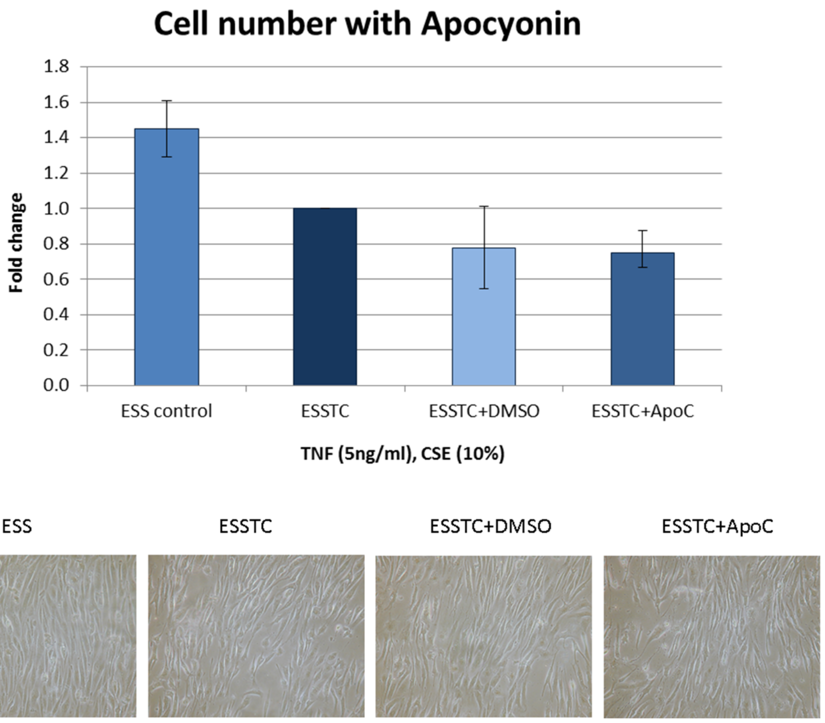

Figure S6: Cell Adhesion of HCAECs exposed to a combination of elevated shear stress, TNFα and CSE together (ESSTC), with additional 200uM Apocynin treatment, vs untreated and DMSO controls at ESS for 72 hours. Cell number was quantified using picogreen assay, expressed as mean fold change against control ± S.E. n=3.

**A pivotal role for Nrf2 in endothelial detachment– implications for endothelial erosion of stenotic plaques.** Sandro Satta, *et. al.*

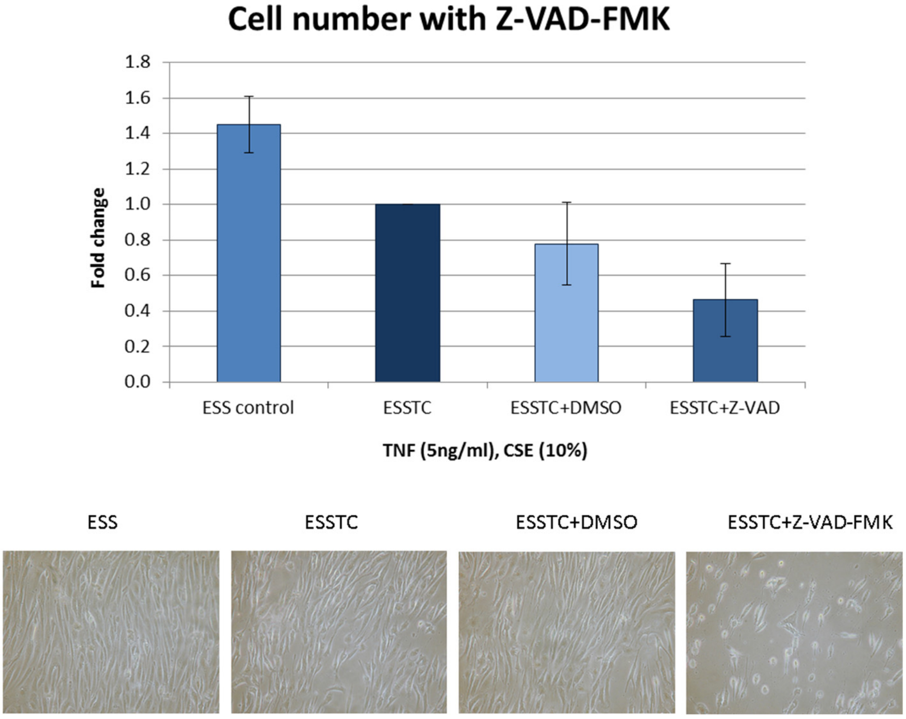

Figure S7: Cell Adhesion of HCAECs exposed to a combination of elevated shear stress, TNFα and CSE together (ESSTC), with additional Z-VAD-FMK treatment (20µM), vs untreated and DMSO controls at ESS for 72 hours. Cell number was quantified using picogreen assay, expressed as mean fold change against control ± S.E. n=3.

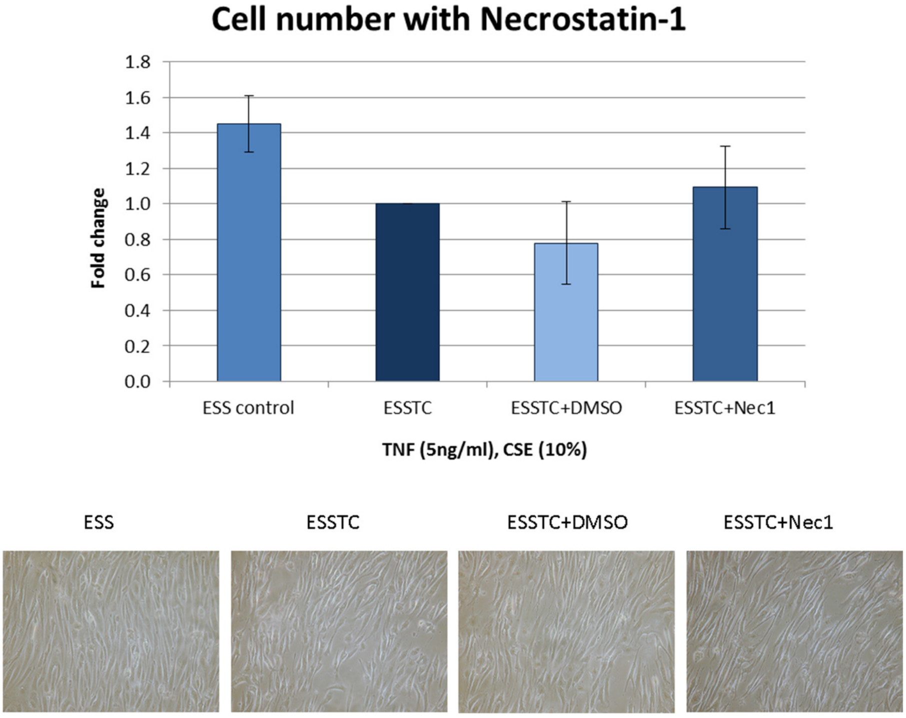

Figure S8: Cell Adhesion of HCAECs exposed to a combination of elevated shear stress, TNFα and CSE together (ESSTC), with additional Necrostatin-1 treatment (10µM), vs untreated and DMSO controls at ESS for 72 hours. Cell number was quantified using picogreen assay, expressed as mean fold change against control ± S.E. n=3.

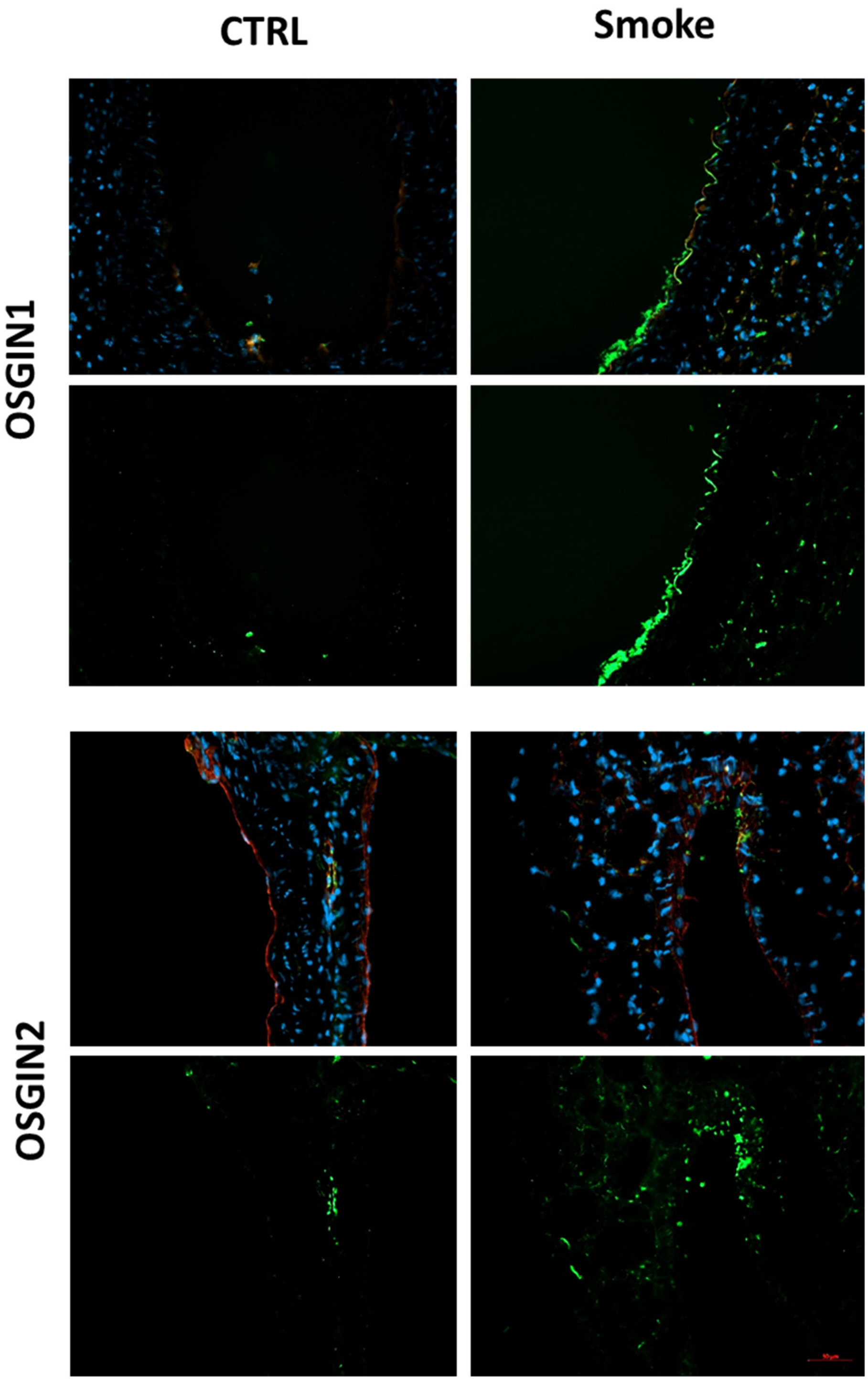

Figure S9. Immunofluorescent analysis of OSGIN1 and OSGIN2 expression in the Aortas from mice exposed to cigarette smoke for 3 months v control. Top tow images for each antibody with DAPI counterstain and endothelial marker (rhodamine-labelled GSL1, 1/100 Vector Labs for OSGIN1 and goat anti-CD31, 10µg/ml Bio-Techne AF3628). Bottom two images for each antibody are just the green channel allowing assessment of OSGIN1 or OSGIN2 staining. OSGIN1 was predominantly localised to the endothelium, while OSGIN2 was found throughout the aorta. Images closest to the median values are presented.

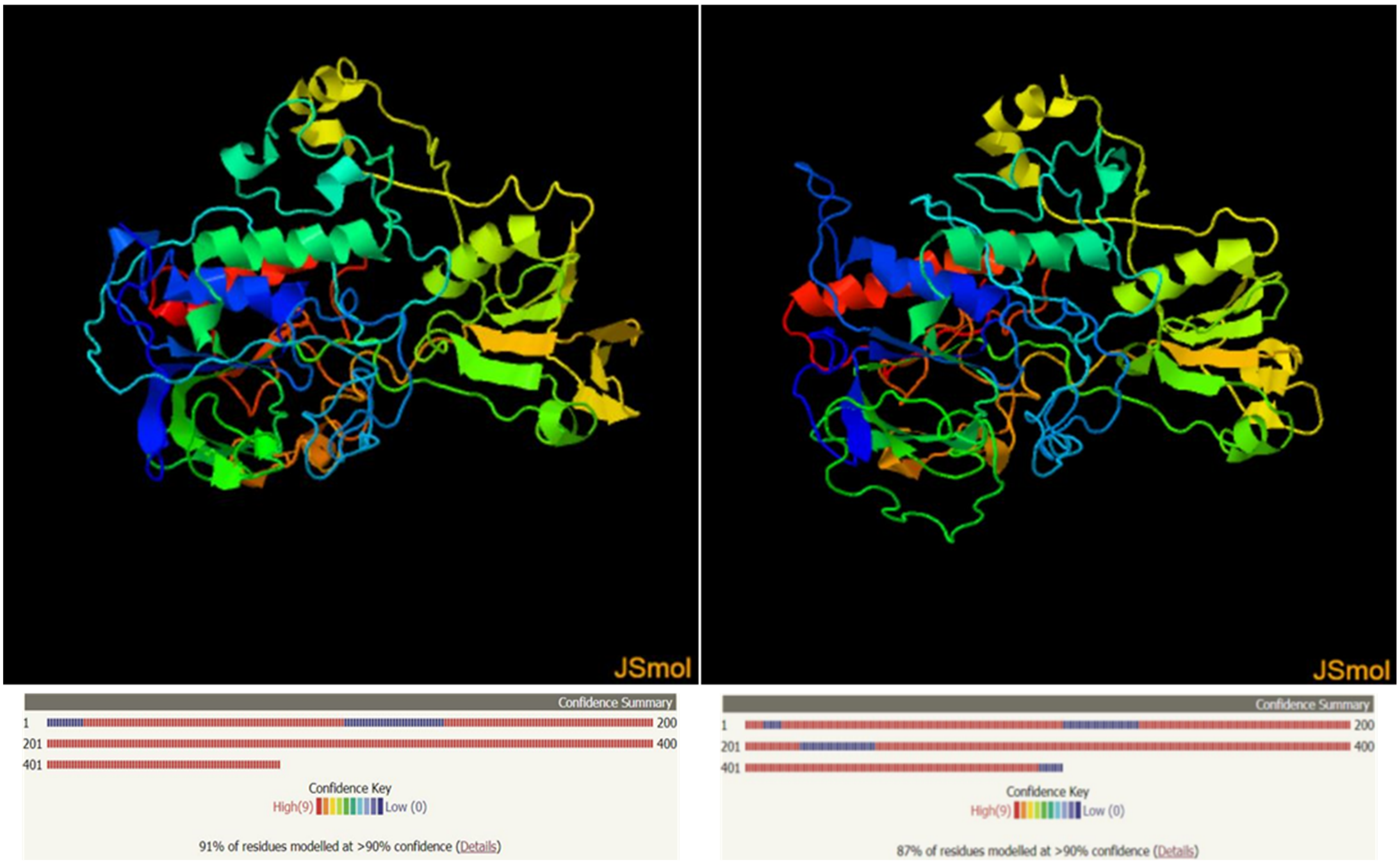

Figure S10. Predicted structure of OSGIN 1 (a) and OSGIN2 (B) generated by RaptorX (http://raptorx.uchicago.edu/StructurePrediction/predict/), demonstrating very similar predicted secondary structure.

```
Q9UJX0 OSGI1_HUMAN      1    MGKWRPRGCCRGNMQCRQEVPATLTSSELFSTRNQPQPQPQPLLADAPVPWAVASRMCLT    60
Q9Y236 OSGI2_HUMAN      1    ------------------------------------------------------------    0

Q9UJX0 OSGI1_HUMAN     61    PGQGCGHQGQDEGPLPAPSPPPAMSSSRKDHLGASSSEPLPVIIVGNGPSGICLSYLLSG    120
Q9Y236 OSGI2_HUMAN      1    -----------------------MPLVEETSLLEDSSVTFPVVIIGNGPSGICLSYMLSG    37
                                                    *   .:  *  .**  :**:*:***********:***

Q9UJX0 OSGI1_HUMAN    121    YTPYTKPDAIHPHPLLQRKLTEAPGVSILDQDLDYLSEGLEGRSQSPVALLFDALLRPDT    180
Q9Y236 OSGI2_HUMAN     38    YRPYLSSEAIHPNTILNSKLEEARHLSIVDQDLEYLSEGLEGRSSNPVAVLFDTLLHPDA    97
                             * **  .  :****:  :*:  ** **   :**:****:**********..***:***:**:**:

Q9UJX0 OSGI1_HUMAN    181    DFGGNMKSVLTWKHRKEHAIPHVVLGRNLPGGAWHSIEGSMVILSQGQWMGLPDLEVKDW    240
Q9Y236 OSGI2_HUMAN     98    DFGYDYPSVLHWKLEQHHYIPHVVLGKGPPGGAWHNMEGSMLTISFGSWMELPGLKFKDW    157
                             ***  :   *** **   .:.* *******:.  ******.:****:  :* *.** **.*:.***

Q9UJX0 OSGI1_HUMAN    241    MQKKRRGLRNSRATAGDIAHYYRDYVVKKGLGHNFVSGAVVTAVEWGTPDPSSCGAQDSS    300
Q9Y236 OSGI2_HUMAN    158    VSSKRRSLKGDRVMPEEIARYYKHYVKVMGLQKNFRENTYITSVSRLYRDQDDDDIQDRD    217
                             :..***.*:..*.    :**:**:.**    **  :**  ..:  :*:*.      *  .. .  ** .

Q9UJX0 OSGI1_HUMAN    301    -----PLFQVSGFLT-----------RNQAQQPFSLWARNVVLATGTFDSPARLGIPGEA    344
Q9Y236 OSGI2_HUMAN    218    ISTKHLQIEKSNFIKRNWEIRGYQRIADGSHVPFCLFAENVALATGTLDSPAHLEIEGED    277
                                  :: *.*:.                 : ::  **.*:*.**.*****:****:* * **

Q9UJX0 OSGI1_HUMAN    345    LPFIHHELSALEAATRVGAVTPASDPVLIIGAGLSAADAVLYARHYNIPVIHAFRRAVDD    404
Q9Y236 OSGI2_HUMAN    278    FPFVFHSMPEFGAAINKGKLRGKVDPVLIVGSGLTAADAVLCAYNSNIPVIHVFRRRVTD    337
                             :**:.*.:   : **  .  *  :      *****:*:**:******  *  :  ******.*** * *

Q9UJX0 OSGI1_HUMAN    405    PGLVFNQLPKMLYPEYHKVHQMMREQSIL-SPSPYEGYRSLPRHQLLCFKEDCQAVFQDL    463
Q9Y236 OSGI2_HUMAN    338    PSLIFKQLPKKLYPEYHKVYHMMCTQSYSVDSNLLSDYTSFPEHRVLSFKSDMKCVLQSV    397
                             *.*:*:****  ********::**    **         .  .    ..* *:*.*::*.**.*  :.*:*.:

Q9UJX0 OSGI1_HUMAN    464    EGVEKVFGVSLVLVLIGSHPDLSFLPGAGADFAVDPDQPLSAKRNPIDVDPFTYQSTRQE    523
Q9Y236 OSGI2_HUMAN    398    SGLKKIFKLSAAVVLIGSHPNLSFLKDQGCYLGHKSSQPITCKGNPVEIDTYTYECIKEA    457
                             .*::*:*  :*   .:*******:****  .  *.  :.  .  .**::.*  **:::*  :**:.  ::

Q9UJX0 OSGI1_HUMAN    524    GLYAMGPLAGDNFVRFVQGGALAVASSLLRKETRKPP-----------    560
Q9Y236 OSGI2_HUMAN    458    NLFALGPLVGDNFVRFLKGGALGVTRCLATRQKKKHLFVERGGGDGIA    505
                             .*:*:***.*******::****.*:  .*    ::.:*
```

| | |
|---|---|
| Date of job execution | Sep 4, 2018 |
| Job identifier | A2018090483C3DD8CE55183C76102DC5D3A26728B27CAA1L (jobs are stored for 7 days) |
| Running time | 133.9 seconds |
| Identical positions | 240 |
| Identity | 40.816% |
| Similar positions | 143 |
| Program | CLUSTALO |

Figure S11. CLUSTALO protein sequence alignment of *Homo sapiens* (Human) OSGIN1 (Q9UJX0) and OSGIN2 (Q9Y236).

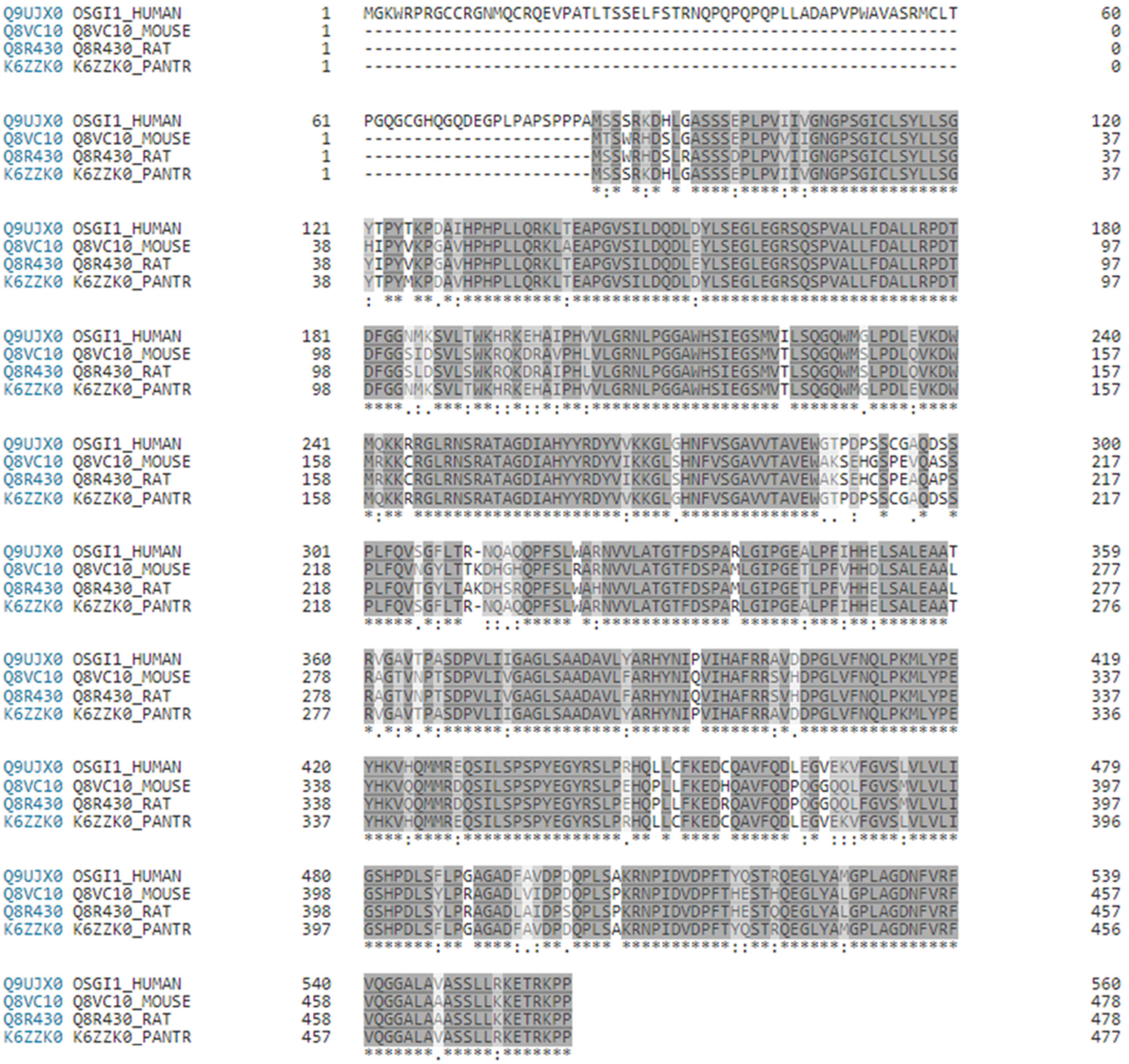

| | |
|---|---|
| Date of job execution | Sep 4, 2018 |
| Job identifier | A2018090483C3DD8CE55183C76102DC5D3A26728B27CC2DW (jobs are stored for 7 days) |
| Running time | 134.9 seconds |
| Identical positions | 385 |
| Identity | 68.627% |
| Similar positions | 66 |
| Program | clustalo |
| Default parameters | Default parameters: The default transition matrix is Gonnet, gap opening penalty is 6 bits, gap extension is 1 bit. Clustal-Omega uses the HHalign algorithm and its default settings as its core alignment engine. The algorithm is described in Söding, J. (2005) 'Protein homology detection by HMM–HMM comparison'. Bioinformatics 21, 951-960. |

Figure S12. CLUSTALO protein sequence alignment of *Homo sapiens* (Human, Q9UJX0), *Mus musculus* (Mouse, Q8VC10), *Rattus norvegicus* (Rat, Q8R430) and *Pan troglodytes* (Chimpanzee, K6ZZK0) OSGIN1.

```
Q9Y236 OSGI2_HUMAN     1    --------------------------------------------MPLVEETSLLEDSSVT    16
Q3TEE9 Q3TEE9_MOUSE    1    MPVWCCRCSLAGHFRNYSDTETEGEIFNSFVQYFGDNLGPKVKGMPLVEETSLLEDSSVT    60
D3ZB49 D3ZB49_RAT      1    MPVWCCRCSLAGHFRNYSDTETEGEIFNSLVQYLGDNLGPKVKGMPLVEETSLLEDSSVT    60
H2QWE6 H2QWE6_PANTR    1    MPVWCCRCSLAGHFRNYSDTETEGEIFNSLVQYFGDNLGRKVKAMPLVEETSLLEDSSVT    60
                                                                        ****************

Q9Y236 OSGI2_HUMAN     17   FPVVIIGNGPSGICLSYMLSGYRPYLSSEAIHPNTILNSKLEEARHLSIVDQDLEYLSEG    76
Q3TEE9 Q3TEE9_MOUSE    61   LPMVIIGNGPSGICLSYMLSGYRPYLSSEAIHPNTILHSKLEEARHLSIVDQDLEYLSEG    120
D3ZB49 D3ZB49_RAT      61   LPMVIIGNGPSGICLSYMLSGYRPYLSSEAIHPNAILHSKLEEARHLSIVDQDLEYLSEG    120
H2QWE6 H2QWE6_PANTR    61   FPVVIIGNGPSGICLSYMLSGYRPYLSSEAIHPNTILNSKLEEARHLSIVDQDLEYLSEG    120
                            :*:*******************************:**:**********************

Q9Y236 OSGI2_HUMAN     77   LEGRSSNPVAVLFDTLLHPDADFGYDYPSVLHWKLEQHHYIPHVVLGKGPPGGAWHNMEG    136
Q3TEE9 Q3TEE9_MOUSE    121  LEGRSSNPVAVLFDTLLHPDADFGYESPSILHWKLEQHHYIPHLVLGKGPPGGAWHNMEG    180
D3ZB49 D3ZB49_RAT      121  LEGRSSNPVAVLFDTLLHPDADFGYDSPSILHWKLEQHHSIPHLVLGKGPPGGAWHNMEG    180
H2QWE6 H2QWE6_PANTR    121  LEGRSSNPVAVLFDTLLHPDADFGYDYPSVLHWKLEQHHYIPHVVLGKGPPGGAWHNMEG    180
                            *************************: **:********* ***:****************

Q9Y236 OSGI2_HUMAN     137  SMLTISFGSWMELPGLKFKDWVSSKRRSLKGDRVMPEEIARYYKHYVKVMGLQKNFRENT    196
Q3TEE9 Q3TEE9_MOUSE    181  SMLTISFGNWMELPGLQFKDWISSKRRNLKGDRVMPEEIARYYKHYVKVMGLQKNFRENT    240
D3ZB49 D3ZB49_RAT      181  SMLTISFGNWMELPGLKFKDWISSKRRNLKGDRVMPEEIARYYKHYVKVMGLQKNFRENT    240
H2QWE6 H2QWE6_PANTR    181  SMLTISFGSWMELPGLKFKDWVSSKRRKLKGDRVMPEEIARYYKHYVKVMGLQKNFRENT    240
                            ********.*******:****:*****.********************************

Q9Y236 OSGI2_HUMAN     197  YITSVSRLYRDQDDDDIQDRDISTKHLQIEKSNFIKRNWEIRGYQRIADGSHVPFCLFAE    256
Q3TEE9 Q3TEE9_MOUSE    241  YITSVSRLYRDQGDNGSQDRDISTKHLQNQKSKFIKRNWEIRGYQRIAGGSHVPFCLFAE    300
D3ZB49 D3ZB49_RAT      241  YITSVSRLYRDQGDNGSQDRDISAKHLQNQKSKFIKRNWEIRGYQRVAGGSHVPFCLFAE    300
H2QWE6 H2QWE6_PANTR    241  YITSVSRLYRDQDDDDSQDRDISTKHLQIEKSNFIKRNWEIRGYQRIADGSHVPFCLFAE    300
                            ************.*:. ******:**** :**:*************:*.***********

Q9Y236 OSGI2_HUMAN     257  NVALATGTLDSPAHLEIEGEDFPFVFHSMPEFGAAINKGKLRGKVDPVLIVGSGLTAADA    316
Q3TEE9 Q3TEE9_MOUSE    301  NVALATGTLDSPAHLEVEGEEFPFVFHSMPEFGAAINKGKLCGRVDPVLIVGSGLTAADA    360
D3ZB49 D3ZB49_RAT      301  NVALATGTLDSPAHLEVEGEEFPFVFHSMPEFGAAINSGKLCGRVDPVLIVGSGLTAADA    360
H2QWE6 H2QWE6_PANTR    301  NVALATGTLDSPAHLEIEGEDFPFVFHSMPEFGAAINKGKLRGKVDPVLIVGSGLTAADA    360
                            ****************:***:****************.*** *:****************

Q9Y236 OSGI2_HUMAN     317  VLCAYNSNIPVIHVFRRRVTDPSLIFKQLPKKLYPEYHKVYHMMCTQSYSVDSNLLSDYT    376
Q3TEE9 Q3TEE9_MOUSE    361  VLCAYNNNIPVIHVFRRRVTDPSLIFKQLPKKLYPEYHKVYHMMCSQSYSADSGPLSDYT    420
D3ZB49 D3ZB49_RAT      361  VLCAYNNNIPVIHVFRRRVTDPSLIFKQLPKKLYPEYHKVYHMMCTQSYSADSDHLSDYT    420
H2QWE6 H2QWE6_PANTR    361  ILCAYNSNIPVIHVFRRRVTDPSLIFKQLPKKLYPEYHKVYHMMCTQSYSVDSNLLSDYT    420
                            :*****.**************************************:****.**. *****

Q9Y236 OSGI2_HUMAN     377  SFPEHRVLSFKSDMKCVLQSVSGLKKIFKLSAAVVLIGSHPNLSFLKDQGCYLGHKSSQP    436
Q3TEE9 Q3TEE9_MOUSE    421  SFPEHRVLSFKADMKCILQSVSGLKKIFKLSAAVILIGSHPNLSFLKEQGCYLGRNSSQP    480
D3ZB49 D3ZB49_RAT      421  SFPEHRVLSFKSDMKCILQSVSGLKKIFKLSAAVILIGSHPNLSFLKEQGCYLGRNSSQP    480
H2QWE6 H2QWE6_PANTR    421  SFPEHHVLSFKSDMKCVLQSVSGLKKIFKLSAAVVLIGSHPNLSFLKDQGCYLGHKSSQP    480
                            *****:*****:****:*****************:************:******::****

Q9Y236 OSGI2_HUMAN     437  ITCKGNPVEIDTYTYECIKEANLFALGPLVGDNFVRFLKGGALGVTRCLATRQKK-KHLF    495
Q3TEE9 Q3TEE9_MOUSE    481  ITCKGNPVEVDAYTYECVKEANLFALGPLVGDNFVRFLKGGALGVTRCLATRQKKKQHLF    540
D3ZB49 D3ZB49_RAT      481  ISCKGNPVEIDAYTYECVKEANLFALGPLVGDNFVRFLKGGALGVTRCLVTRQKKKQHLF    540
H2QWE6 H2QWE6_PANTR    481  ITCKGNPVEIDTYTYECIKEANLFALGPLVGDNFVRFLKGGALGVTRCLATRQKK-KHLF    539
                            *:*******:*:*****:*******************************.***** :***

Q9Y236 OSGI2_HUMAN     496  VERGGGDGIA    505
Q3TEE9 Q3TEE9_MOUSE    541  VQRGGGDGVA    550
D3ZB49 D3ZB49_RAT      541  VQRGGGDGVA    550
H2QWE6 H2QWE6_PANTR    540  VERGGGDGIA    549
                            *:******:*
```

| | |
|---|---|
| Date of job execution | Sep 4, 2018 |
| Job identifier | A20180904F725F458AC8690F874DD868E4ED79B880B95B4W (jobs are stored for 7 days) |
| Running time | 135.9 seconds |
| Identical positions | 456 |
| Identity | 82.909% |
| Similar positions | 43 |
| Program | clustalo |
| Default parameters | Default parameters: The default transition matrix is Gonnet, gap opening penalty is 6 bits, gap extension is 1 bit. Clustal-Omega uses the HHalign algorithm and its default settings as its core alignment engine. The algorithm is described in Söding, J. (2005) 'Protein homology detection by HMM-HMM comparison'. Bioinformatics 21, 951-960. |

Figure S13. CLUSTALO protein sequence alignment of *Homo sapiens* (Human, Q9Y236), *Mus musculus* (Mouse, Q3TEE9), *Rattus norvegicus* (Rat, D3ZB49) and *Pan troglodytes* (Chimpanzee, H2QWE6) OSGIN2.

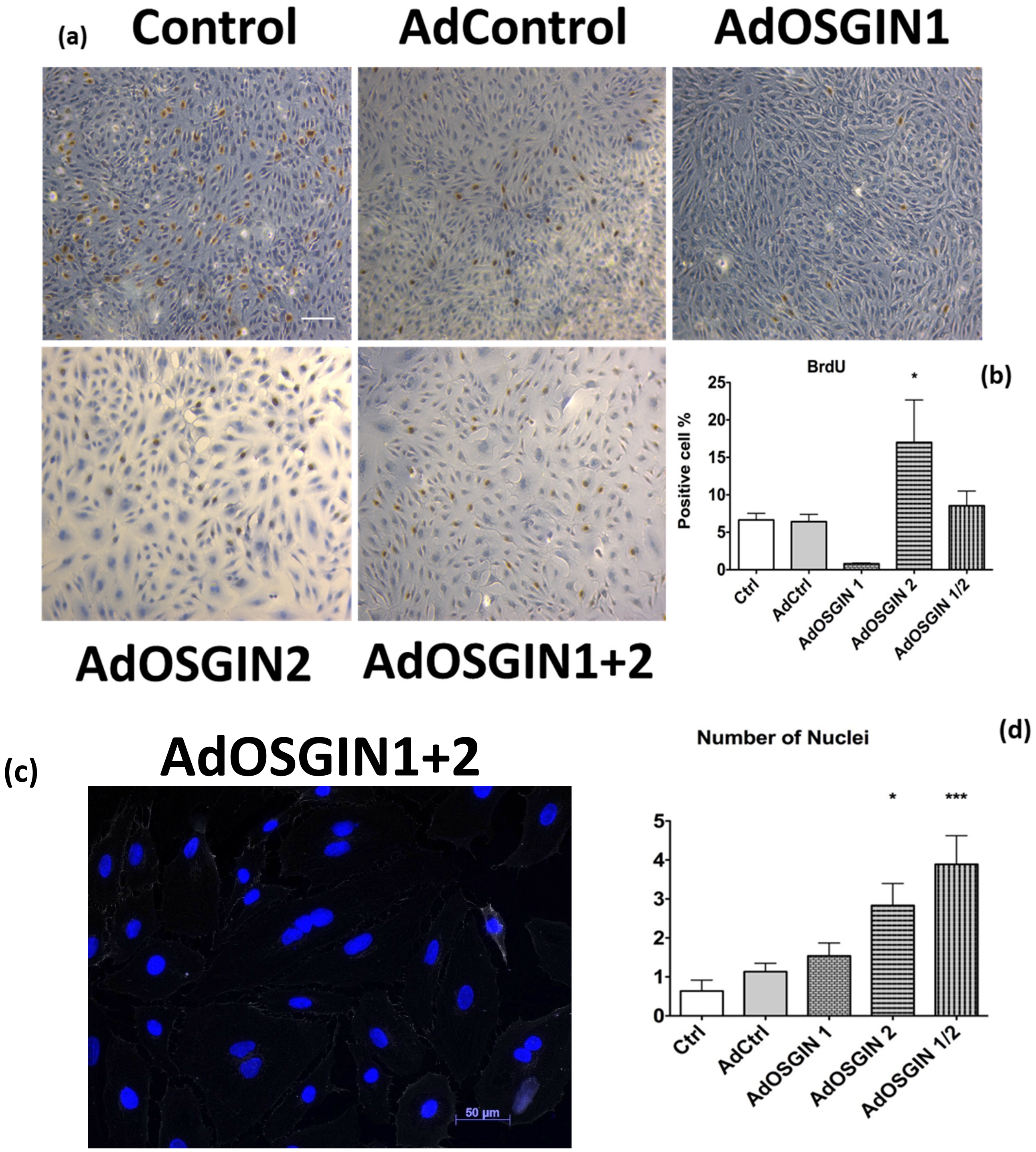

Figure S14. A) Proliferation of HCAECs was analysed using the BrdU incorporation assay. HCAECs transfected with ad OSGIN2 show a lower number of cells compared with the other conditions. Furthermore, the cells are bigger in size and have a high rate of replication. HCAECs transfected with ad OSGIN1/2 show a cell size similar to the ad OSGIN2 overexpression conditions, but higher numbers of cells. B) The graph shows the effect of OSGIN1, OSGIN2 and OSGIN1/2 ad transfection on cell proliferation. * $P<0.05$, in comparison with HCAECs on AdOSGIN2 condition VS AdCtrl overexpression. C) AdOSGIN1/2 condition showed numerous multinucleated cells (highlighted by the arrows) but not BrdU positive. D) Number of nuclei were counted and compared against AdCtrl. AdOSGIN2 and AdOSGIN1/2 showed up to five times more multi nucleated cells *$P<0.05$, and ***$P<0.001$.

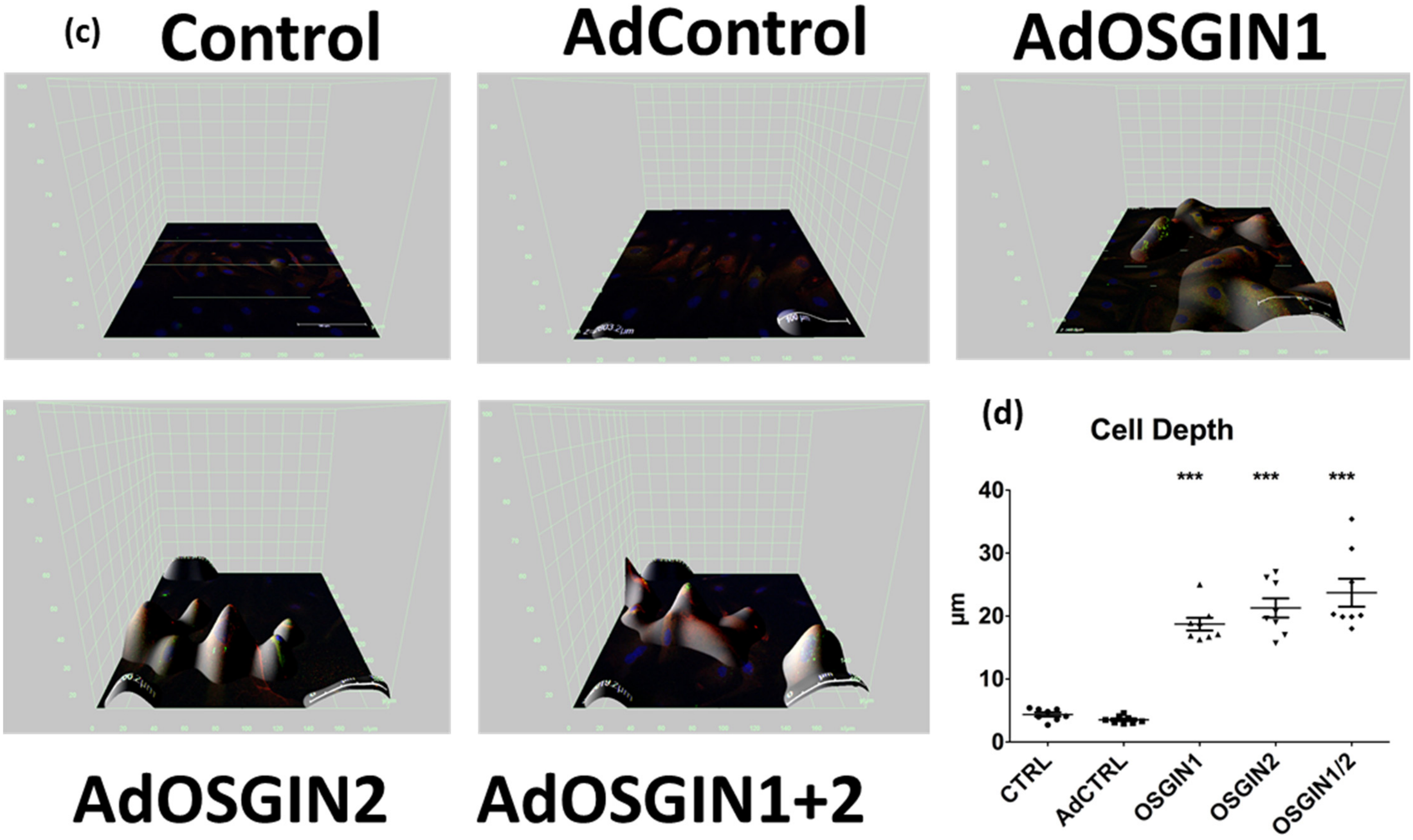

Figure S15: 3D cell model was created using Leica software confocal image stack in combination with Image j: image to stack from 50 layers of confocal images. Cell depth was measured with an increment of cell height up to 25µm (n=3; P<0.001).

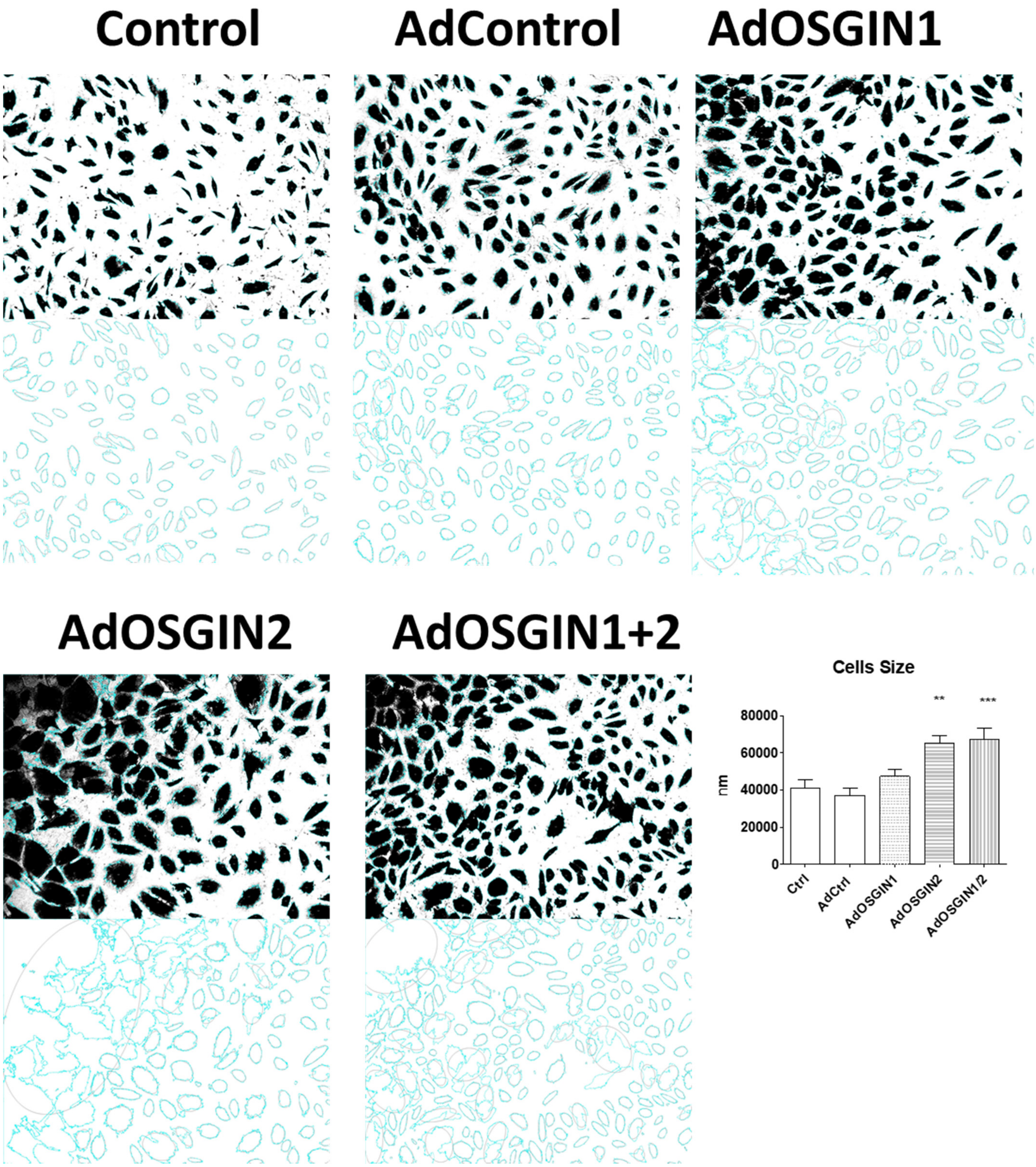

Figure S16: EC size analysis was evaluated through Image J custom particles analyser and multi cell outliner (ellipse fit correction). Five random pictures were analysed, and each cell area was automatically measured. AdOSGIN2 and AdOSGIN1+2 condition showed an increment in cell size with almost double the size compared to the Ctrl and AdCtrl. **$P<0.01$ AdOSGIN2 and ***$P<0.001$ AdOSGIN1+2 vs AdCtrl.

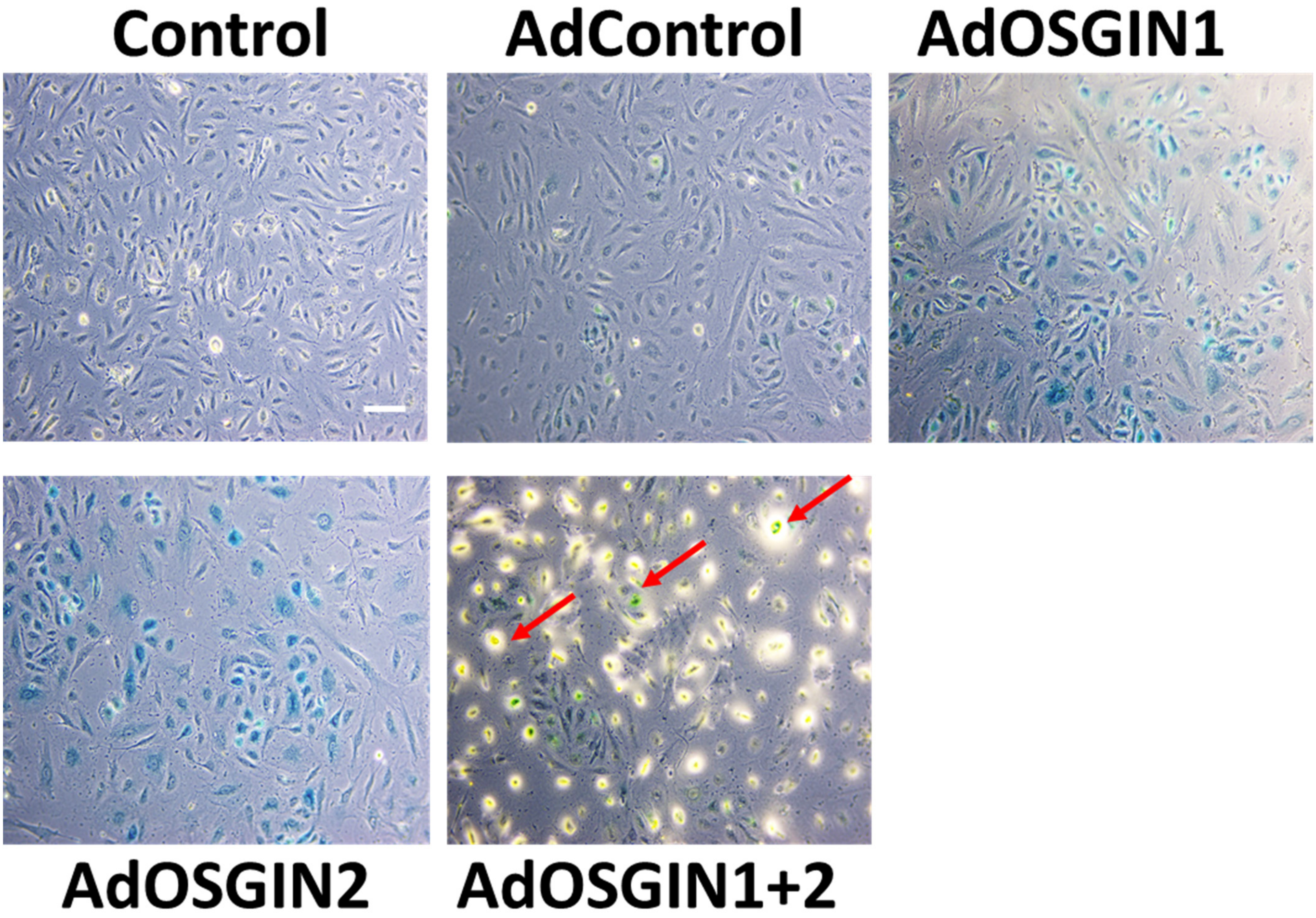

Figure S17. Senescence-associated beta-galactosidase (SA-β-galactosidase) senescence staining of HAECs transfected with adenoviral (OSGIN1, OSGIN2 and OSGIN1/2). Overexpression by ad of AdOSGIN1 and AdOSGIN2 induces senescent phenotypes in HCAECs. AdOSGIN1/2 overexpression shows green/blue senescence cells (red arrows) detached or in progress of detaching. SA-β-galactosidase assays were performed 40h after adenovirus transfection.

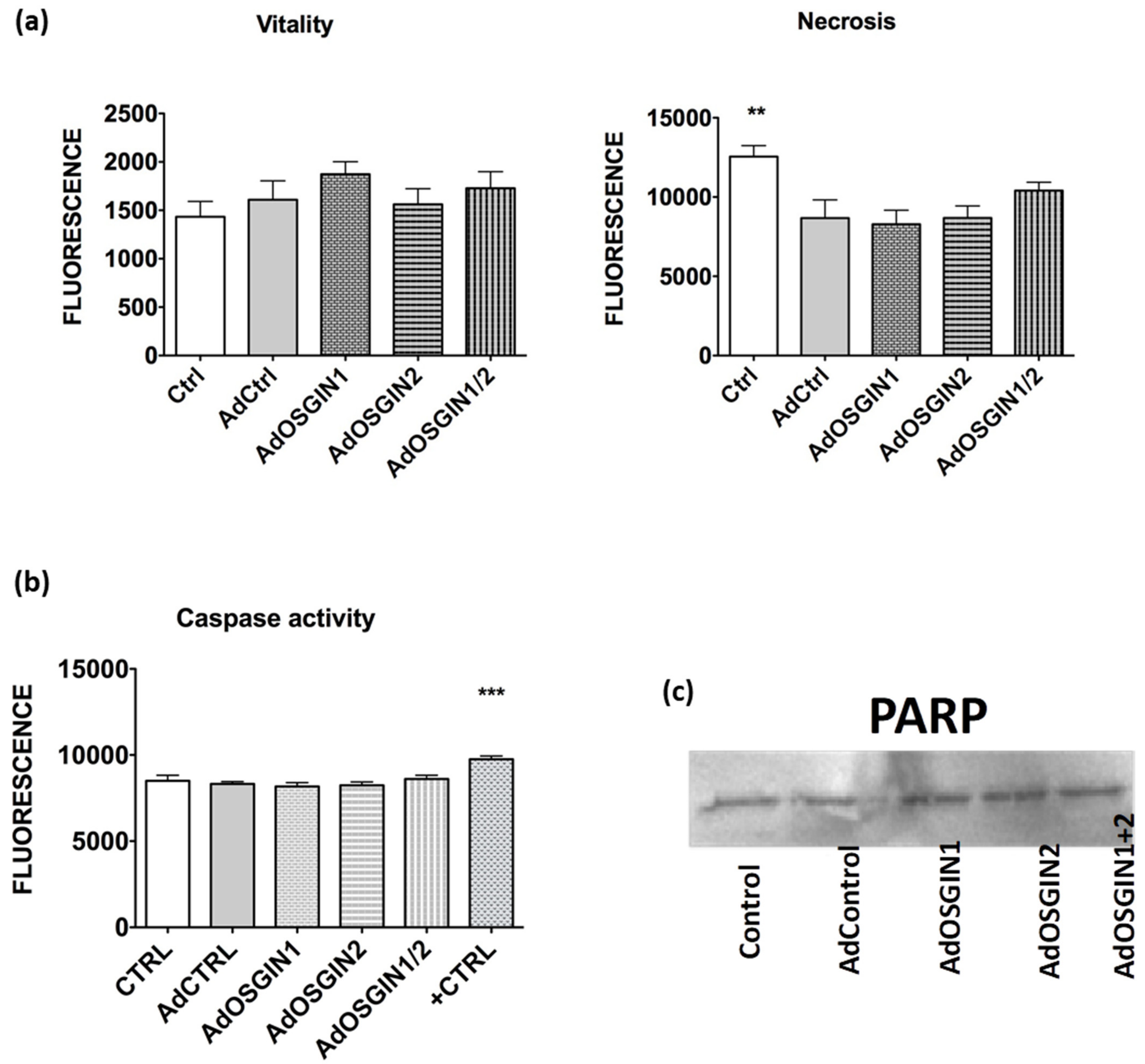

Figure S18 **a)** Cell viability assay for HCAECs. The cell viability was measured post transfection at 24h (adenovirus overexpression of OSGIN1, OSGIN2 and both together) and did not cause a significant decrease in cell viability. Overexpression of OSGIN1, OSGIN2 and both together did not cause a significant increase or decrease in necrosis. Viability and Necrosis were measured using promega Apotox Glo assay kit. **b)** Caspase 3/7 activity of HCAECs transfected with adenovirus overexpressing OSGIN1, OSGIN 2 and both together, compared to AdCtrl transfections and positive control (0.2mM H2O2). The caspase 3/7 activity was measured using Caspase-Glo 3/7 assay. Data is presented as mean ± SEM (n = 3). *** $p < 0.001$. **c)** Total Parp antibody in western blotting showed no parp cleavage by caspase-3. No second band was reported in any sample suggesting no apoptosis activity.

**A pivotal role for Nrf2 in endothelial detachment– implications for endothelial erosion of stenotic plaques.** Sandro Satta, *et. al.*

Supplementary Tables S10 Cluster 1, Genes and top 10 canonical pathways and regulators.

| **Genes** | |
|---|---|
| AHSA1 | HSPB1 |
| ALDH3A2 | HSPB8 |
| ATF3 | HSPD1 |
| B3GAT3 | HSPE1 |
| BAG3 | HSPH1 |
| BANF1 | IER5L |
| C3orf52 | IL7R |
| CACYBP | JMJD6 |
| CBARP | LGALS8 |
| CHKA | LUC7L3 |
| CHORDC1 | MICB |
| CPNE8 | MLKL |
| CRYAB | MRPL18 |
| CRYZ | MSX1 |
| DEDD2 | NAPG |
| DNAJA1 | NDRG1 |
| DNAJB1 | OSGIN1 |
| DNAJB6 | P4HA2 |
| EVI2B | PATL1 |
| FKBP4 | PAXBP1 |
| FTL | PLAUR |
| GDF15 | PLEKHB2 |
| HIF3A | PRPF38B |
| HIST2H2BE | PTGES3 |
| HMOX1 | RNF19B |
| HSP90AA1 | RSRP1 |
| HSP90AB1 | SNX3 |
| HSPA1A | SPATS2L |
| HSPA1B | SQSTM1 |
| HSPA4 | STIP1 |
| HSPA6 | TRIM16 |
| HSPA8 | TSPYL2 |

| **Ingenuity Canonical Pathways** |
|---|
| Aldosterone Signaling in Epithelial Cells |
| Protein Ubiquitination Pathway |
| NRF2-mediated Oxidative Stress Response |
| Unfolded protein response |
| Glucocorticoid Receptor Signaling |
| eNOS Signaling |
| Aryl Hydrocarbon Receptor Signaling |
| Huntington's Disease Signaling |
| Xenobiotic Metabolism Signaling |
| Ingenuity Canonical Pathways |

| **Upstream Transcriptional Regulator** |
|---|
| HSF1 |
| FBXW7 |
| PML |
| SP100 |
| NFE2L2 |
| TP53 |
| ETS1 |
| HSF2 |
| HTT |
| NUPR1 |

Supplementary Tables S11 Cluster 2, Genes and top 10 canonical pathways and regulators.

| **Genes** | |
|---|---|
| ANKRD10 | MX1 |
| APOL6 | MX2 |
| BATF2 | OAS1 |
| C19orf66 | OAS2 |
| DDX58 | OASL |
| DDX60 | OSGIN2 |
| DDX60L | PARP14 |
| DHX58 | PARP9 |
| DNAJA4 | PLSCR1 |
| DTX3L | PMAIP1 |
| EIF2AK2 | PPP1R18 |
| EPSTI1 | RSAD2 |
| GBP1 | SAMD9 |
| GCH1 | SAMD9L |
| HERC6 | SERPINH1 |
| IFI35 | SLC15A3 |
| IFI44L | SLC37A1 |
| IFIH1 | SP110 |
| IFIT1 | TAP1 |
| IFIT2 | TRIM21 |
| IFIT3 | TRIM26 |
| IFIT5 | TRIM69 |
| IFITM1 | UBC |
| IRF7 | UBE2L6 |
| ISG15 | XAF1 |
| LY6E | |

| **Ingenuity Canonical Pathways** |
|---|
| Interferon Signaling |
| Activation of IRF by Cytosolic Pattern Recognition Receptors |
| Role of Pattern Recognition Receptors in Recognition of Bacteria and Viruses |
| Role of RIG1-like Receptors in Antiviral Innate Immunity |
| Tetrahydrobiopterin Biosynthesis I |
| Tetrahydrobiopterin Biosynthesis II |
| Retinoic acid Mediated Apoptosis Signaling |
| Toll-like Receptor Signaling |
| Death Receptor Signaling |
| Protein Ubiquitination Pathway |

| **Upstream Transcriptional Regulator** |
|---|
| IRF7 |
| STAT1 |
| IRF1 |
| NKX2-3 |
| IRF3 |
| IRF5 |
| TRIM24 |
| CNOT7 |
| STAT3 |
| STAT2 |

Supplementary Tables S12 Cluster 3, Genes and top 10 canonical pathways and regulators.

| Genes | |
|---|---|
| ADM | PRDX6 |
| ANGPT2 | SBDSP1 |
| F2RL2 | SEC61G |
| GCLM | SELO |
| INA | SLC17A9 |
| LENG9 | SNRPA1 |
| ME1 | TBXAS1 |
| MLLT11 | TKT |
| NQO1 | TMEM156 |
| P4HA1 | TRIM16L |
| PDE4B | UPP1 |
| PLPP2 | XPOT |
| PRDX1 | ZDHHC6 |

| **Ingenuity Canonical Pathways** |
|---|
| NRF2-mediated Oxidative Stress Response |
| Glutathione Biosynthesis |
| Pentose Phosphate Pathway (Non-oxidative Branch) |
| Superoxide Radicals Degradation |
| Prostanoid Biosynthesis |
| Phagosome Maturation |
| Pentose Phosphate Pathway |
| γ-glutamyl Cycle |
| Glutathione Redox Reactions I |
| Gluconeogenesis I |

| **Upstream Transcriptional Regulator** |
|---|
| MAFG |
| BACH1 |
| NFE2L2 |
| KEAP1 |
| MAFK |
| NFE2 |
| PML |
| KLF2 |
| NFE2L1 |
| PDX1 |

Supplementary Tables S13. Cluster 4, Genes and top 10 canonical pathways and regulators.

| **Genes** |
|---|
| ANGPTL4 |
| CTD-2015G9.2 |
| HOXA9 |
| HOXB7 |
| MT-CYB |
| RPSAP58 |
| TNFSF18 |

| **Ingenuity Canonical Pathways** |
|---|
| Oxidative Phosphorylation |
| Mitochondrial Dysfunction |
| Sirtuin Signaling Pathway |

| **Upstream Transcriptional Regulator** |
|---|
| PHF1 |
| COMMD3-BMI1 |
| KMT2A |
| MLLT1 |
| ASB2 |
| NKX2-3 |
| PSIP1 |
| HOXB3 |
| BHLHE41 |
| PLAGL1 |

Supplementary Tables S14. Cluster 5, Genes and top 10 canonical pathways and regulators.

| Genes | |
|---|---|
| ALDH1A3 | MFSD11 |
| AMDHD2 | NUP107 |
| ANAPC5 | NUP160 |
| ANKRD33B | PACS1 |
| AP3M2 | PAN2 |
| ATXN2 | PCMTD2 |
| C22orf29 | PIGQ |
| CAPN11 | PLA2G4C |
| CHD1L | PLD1 |
| CLIP4 | PPCS |
| CPNE5 | RCBTB1 |
| DFNA5 | RFC3 |
| DHRS1 | RNF123 |
| DHRS11 | SESTD1 |
| EEF2K | SLC35E2B |
| EVL | SMARCC2 |
| HAS3 | STK38L |
| HERC2 | SYNE2 |
| ITPR3 | TGFA |
| KRIT1 | UQCRC2 |
| LETMD1 | VPS13C |
| LRRC16A | WDR59 |
| MED12 | ZC3H7A |

| Ingenuity Canonical Pathways |
|---|
| Coenzyme A Biosynthesis |
| N-acetylglucosamine Degradation I |
| Phospholipases |
| Endothelin-1 Signaling |
| N-acetylglucosamine Degradation II |
| Role of BRCA1 in DNA Damage Response |
| Non-Small Cell Lung Cancer Signaling |
| Phospholipase C Signaling |
| Antioxidant Action of Vitamin C |
| Pancreatic Adenocarcinoma Signaling |

| Upstream Transcriptional Regulator |
|---|
| FOXH1 |
| CITED1 |
| Ncoa6 |
| GSX2 |
| BHLHA15 |
| FOS |
| VAV2 |

Supplementary Tables S15. Cluster 6, Genes and top 10 canonical pathways and regulators.

| Genes |
|---|
| BRIP1 |
| ESCO2 |

| Ingenuity Canonical Pathways |
|---|
| Role of BRCA1 in DNA Damage Response |

| Upstream Transcriptional Regulator |
|---|
| CCND1 |
| UXT |
| FOXM1 |
| IRF1 |
| TCF4 |
| TCF3 |
| FOXO1 |

Supplementary Tables S16 Cluster 7, Genes and top 10 canonical pathways and regulators.

| **Genes** | |
|---|---|
| ABCA1 | LAMA2 |
| AEBP1 | LGALS9 |
| AGRN | LYPD1 |
| APOL1 | LYVE1 |
| BNC1 | MELTF |
| CARD11 | MGP |
| CCDC3 | MYOM3 |
| CDH11 | NEAT1 |
| CFH | NEFH |
| CLDN11 | PCDH10 |
| CSF2RB | PCDH17 |
| CTHRC1 | PLCB2 |
| CX3CL1 | PLCD1 |
| DDR2 | PLCL1 |
| DGKA | POSTN |
| DKK2 | PSMB9 |
| DOCK5 | PTPRU |
| DOCK8 | SAT1 |
| DPP4 | SELENBP1 |
| ERAP1 | SEMA7A |
| FAM129A | SLITRK4 |
| FRAS1 | ST6GALNAC1 |
| GATSL3 | STXBP2 |
| HSPA12B | TACSTD2 |
| IGFBP5 | TFAP2A |
| IL18R1 | TRPV2 |
| ITGB3 | TUSC3 |
| KLF4 | VCAM1 |
| KRT19 | |

| **Ingenuity Canonical Pathways** |
|---|
| Adrenomedullin signaling pathway |
| Role of Macrophages, Fibroblasts and Endothelial Cells in Rheumatoid Arthritis |
| Antioxidant Action of Vitamin C |
| P2Y Purigenic Receptor Signaling Pathway |
| Phospholipases |
| Wnt/Ca+ pathway |
| Melatonin Signaling |
| GPCR-Mediated Integration of Enteroendocrine Signaling Exemplified by an L Cell |
| Aldosterone Signaling in Epithelial Cells |
| Agrin Interactions at Neuromuscular Junction |

| **Upstream Transcriptional Regulator** |
|---|
| HTT |
| NKX2-3 |
| HOXA10 |
| CREB1 |
| SMARCA4 |
| MECP2 |
| STAT1 |
| FOXF1 |
| IRF2 |
| ATF4 |

Supplementary Tables S17. Cluster 8, Genes and top 10 canonical pathways and regulators.

| **Genes** | HSPG2 |
|---|---|
| A2M | IGFBP2 |
| ABCA7 | IL17RD |
| ACE | IL33 |
| ACP5 | ITGA10 |
| ADCY4 | ITPR1 |
| ANK1 | KCND1 |
| ANK3 | KCNN4 |
| APLNR | LIMCH1 |
| APOB | LPL |
| APOL4 | LRP1 |
| AQP1 | MAMDC2 |
| AQP3 | MAN1C1 |
| ARHGEF9 | MAN2B2 |
| ATHL1 | MFAP2 |
| ATP2A3 | MMP17 |
| B3GNT9 | MYO18A |
| BTN3A3 | NPR1 |
| C10orf10 | PBX1 |
| C10orf128 | PCMTD1 |
| C10orf54 | PEG10 |
| C14orf132 | PIDD1 |
| C1RL | PIK3R3 |
| CALCOCO1 | PLPP3 |
| CCND2 | PPARGC1B |
| CD24 | PPP1R14A |
| CD40 | PTGIS |
| CECR1 | RAPGEF5 |
| CFI | RP1-152L7.5 |
| CKB | RRAGB |
| CMKLR1 | SCUBE3 |
| COL17A1 | SEMA3G |
| COL1A2 | SGCE |
| COL4A5 | SIAE |
| COLEC12 | SIDT2 |
| DPYSL4 | SLC16A14 |
| DYSF | SLC46A3 |
| EDA2R | SLCO2A1 |
| ENTPD1 | SLCO2B1 |
| EPS8L2 | SNED1 |
| ERV3-1 | SORT1 |
| FBLN2 | STAB1 |
| FBN1 | SULF1 |
| FKBP9 | TMOD1 |
| GAA | TNFRSF14 |
| GAL3ST4 | TNS1 |

| **Ingenuity Canonical Pathways** |
|---|
| Atherosclerosis Signaling |
| Inhibition of Matrix Metalloproteases |
| Adrenomedullin signaling pathway |
| Gap Junction Signaling |
| GP6 Signaling Pathway |
| eNOS Signaling |
| Hepatic Fibrosis / Hepatic Stellate Cell Activation |
| Renin-Angiotensin Signaling |
| Breast Cancer Regulation by Stathmin1 |
| Dopamine-DARPP32 Feedback in cAMP Signaling |

| **Upstream Transcriptional Regulator** |
|---|
| KLF2 |
| TP53 |
| SP1 |
| TCF4 |
| CALR |
| CEBPA |
| BRD7 |
| ETS1 |
| IKZF1 |
| ASB9 |

| | |
|---|---|
| GJA4 | TPCN1 |
| GJA5 | TRIM66 |
| GPR146 | USHBP1 |
| HEG1 | VASH1 |
| HHAT | VCAN |
| HMCN1 | ZNF366 |

**A pivotal role for Nrf2 in endothelial detachment– implications for endothelial erosion of stenotic plaques.** Sandro Satta, *et. al.*

Supplementary Tables S18. AdOSGIN1 vs AdControl, Genes and top 20 canonical pathways and regulators.

| **Genes** | | | | |
|---|---|---|---|---|
| A2M | CMKLR1 | HSPA1B | OSGIN2 | SLITRK4 |
| ABCA1 | COL17A1 | HSPA4 | P4HA1 | SNED1 |
| ABCA7 | COL1A2 | HSPA6 | PACS1 | SNRPA1 |
| ACE | COL4A5 | HSPB8 | PAN2 | SORT1 |
| ACP5 | COLEC12 | HSPD1 | PBX1 | ST6GALNAC1 |
| ADCY4 | CPNE5 | HSPG2 | PCDH10 | STAB1 |
| ADM | CRYAB | IGFBP2 | PCDH17 | STK38L |
| AEBP1 | CSF2RB | IGFBP5 | PCMTD1 | STXBP2 |
| AGRN | CTD-2015G9.2 | IL17RD | PCMTD2 | SULF1 |
| ALDH1A3 | CTHRC1 | IL18R1 | PDE4B | SYNE2 |
| AMDHD2 | CX3CL1 | IL33 | PEG10 | TACSTD2 |
| ANGPT2 | DDR2 | INA | PIDD1 | TBXAS1 |
| ANGPTL4 | DGKA | ITGA10 | PIK3R3 | TFAP2A |
| ANK1 | DHRS1 | ITGB3 | PLA2G4C | TGFA |
| ANK3 | DKK2 | ITPR1 | PLCB2 | TKT |
| ANKRD33B | DOCK5 | ITPR3 | PLCD1 | TMEM156 |
| AP3M2 | DOCK8 | JMJD6 | PLCL1 | TMOD1 |
| APLNR | DPP4 | KCND1 | PLPP2 | TNFRSF14 |
| APOB | DPYSL4 | KCNN4 | PLPP3 | TNFSF18 |
| APOL1 | DYSF | KLF4 | POSTN | TNS1 |
| APOL4 | EDA2R | KRT19 | PPARGC1B | TPCN1 |
| AQP1 | ENTPD1 | LAMA2 | PPP1R14A | TRIM16 |
| AQP3 | EPS8L2 | LENG9 | PRDX1 | TRIM16L |
| ARHGEF9 | ERAP1 | LETMD1 | PRDX6 | TRIM66 |
| ATHL1 | ERV3-1 | LGALS9 | PSMB9 | TRPV2 |
| ATP2A3 | EVL | LIMCH1 | PTGES3 | TUSC3 |
| B3GNT9 | F2RL2 | LPL | PTGIS | UPP1 |
| BNC1 | FAM129A | LRP1 | PTPRU | UQCRC2 |

| **Ingenuity Canonical Pathways** |
|---|
| Aldosterone Signaling in Epithelial Cells |
| Adrenomedullin signaling pathway |
| Atherosclerosis Signaling |
| eNOS Signaling |
| Hepatic Fibrosis / Hepatic Stellate Cell Activation |
| Sperm Motility |
| Cellular Effects of Sildenafil (Viagra) |
| Dopamine-DARPP32 Feedback in cAMP Signaling |
| Gap Junction Signaling |
| Prostanoid Biosynthesis |
| Endothelin-1 Signaling |
| GP6 Signaling Pathway |
| GPCR-Mediated Integration of Enteroendocrine Signaling Exemplified by an L Cell |
| Synaptic Long Term Depression |
| Neuropathic Pain Signaling In Dorsal Horn Neurons |
| Dendritic Cell Maturation |
| Phospholipases |
| Inhibition of Matrix Metalloproteases |
| Thrombin Signaling |
| Protein Ubiquitination Pathway |

| **Upstream Transcriptional Regulator** |
|---|
| TP53 |
| EPAS1 |
| HIF1A |
| KLF2 |
| NKX2-3 |
| NEUROG1 |
| SP1 |
| NFKBIA |
| ETS1 |
| HTT |
| STAT3 |
| NUPR1 |
| TWIST1 |
| MAFG |
| MYB |
| KEAP1 |
| KDM3A |
| BACH1 |
| NFE2L2 |
| HOXA10 |

| | | | | |
|---|---|---|---|---|
| BTN3A3 | FBLN2 | LRRC16A | RAPGEF5 | USHBP1 |
| C10orf10 | FBN1 | LYPD1 | RCBTB1 | VASH1 |
| C10orf128 | FKBP4 | LYVE1 | RNF123 | VCAM1 |
| C10orf54 | FKBP9 | MAMDC2 | RP1-152L7.5 | VCAN |
| C14orf132 | FRAS1 | MAN1C1 | RPSAP58 | VPS13C |
| C1RL | GAA | MAN2B2 | RRAGB | XPOT |
| CACYBP | GAL3ST4 | ME1 | SAT1 | ZNF366 |
| CALCOCO1 | GATSL3 | MELTF | SCUBE3 | |
| CAPN11 | GCLM | MFAP2 | SEC61G | |
| CARD11 | GDF15 | MFSD11 | SELENBP1 | |
| CCDC3 | GJA4 | MGP | SELO | |
| CCND2 | GJA5 | MLLT11 | SEMA3G | |
| CD24 | GPR146 | MMP17 | SEMA7A | |
| CD40 | HEG1 | MT-CYB | SGCE | |
| CDH11 | HHAT | MYO18A | SIAE | |
| CECR1 | HMCN1 | MYOM3 | SIDT2 | |
| CFH | HMOX1 | NEAT1 | SLC16A14 | |
| CFI | HOXA9 | NEFH | SLC17A9 | |
| CHORDC1 | HOXB7 | NPR1 | SLC35E2B | |
| CKB | HSP90AA1 | NQO1 | SLC46A3 | |
| CLDN11 | HSPA12B | NUP160 | SLCO2A1 | |
| CLIP4 | HSPA1A | OSGIN1 | SLCO2B1 | |

Supplementary Tables S19. AdOSGIN2 vs AdControl, Genes and top 20 canonical pathways and regulators.

| Genes |
|---|
| HERC6 |
| HSPA6 |
| IFIT1 |
| IFIT3 |
| MX2 |
| OASL |
| OSGIN2 |
| RSAD2 |
| XAF1 |

| Ingenuity Canonical Pathways |
|---|
| Interferon Signaling |
| Role of Lipids/Lipid Rafts in the Pathogenesis of Influenza |
| Unfolded protein response |
| Aldosterone Signaling in Epithelial Cells |
| eNOS Signaling |
| Huntington's Disease Signaling |
| Protein Ubiquitination Pathway |
| Glucocorticoid Receptor Signaling |

| Upstream Transcriptional Regulator |
|---|
| IRF7 |
| STAT3 |
| STAT1 |
| IRF1 |
| IRF5 |
| IRF3 |
| STAT2 |
| SPI1 |
| TRIM24 |
| NFATC2 |
| IRF9 |
| MSC |
| BRCA1 |
| SP100 |
| IKZF3 |
| SIRT1 |
| CNOT7 |
| SQSTM1 |
| POU2AF1 |
| HOXD10 |

Supplementary Tables S20. AdOSGIN1&2 vs AdControl, Genes and top 20 canonical pathways and regulators.

**A pivotal role for Nrf2 in endothelial detachment– implications for endothelial erosion of stenotic plaques.** Sandro Satta, *et. al.*

| Genes | | | |
|---|---|---|---|
| AHSA1 | GCLM | NUP107 | TRIM16 |
| ALDH3A2 | GDF15 | OAS1 | TRIM16L |
| AMDHD2 | HAS3 | OAS2 | TRIM21 |
| ANAPC5 | HERC2 | OASL | TRIM26 |
| ANGPT2 | HERC6 | OSGIN1 | TRIM69 |
| ANKRD10 | HIF3A | OSGIN2 | TSPYL2 |
| AP3M2 | HIST2H2BE | P4HA1 | UBC |
| APOL6 | HMOX1 | P4HA2 | UBE2L6 |
| ATF3 | HSP90AA1 | PACS1 | UQCRC2 |
| ATXN2 | HSP90AB1 | PARP14 | UTP3 |
| B3GAT3 | HSPA1A | PARP9 | VPS13C |
| BAG3 | HSPA1B | PATL1 | WDR59 |
| BANF1 | HSPA4 | PAXBP1 | XAF1 |
| BATF2 | HSPA6 | PIGQ | ZC3H7A |
| BRIP1 | HSPA8 | PLAUR | ZDHHC6 |
| C19orf66 | HSPB1 | PLD1 | ZFAND2A |
| C22orf29 | HSPB8 | PLEKHB2 | ZNF207 |
| C3orf52 | HSPD1 | PLSCR1 | ZNF622 |
| CACYBP | HSPE1 | PMAIP1 | |
| CBARP | HSPH1 | PPCS | |
| CHD1L | IER5L | PPP1R18 | |
| CHKA | IFI35 | PRDX1 | |
| CHORDC1 | IFI44L | PRPF38B | |
| CLIP4 | IFIH1 | PTGES3 | |
| CPNE8 | IFIT1 | RCBTB1 | |
| CRYAB | IFIT2 | RFC3 | |
| CRYZ | IFIT3 | RNF123 | |
| DDX58 | IFIT5 | RNF19B | |
| DDX60 | IFITM1 | RSAD2 | |

| Ingenuity Canonical Pathways |
|---|
| Aldosterone Signaling in Epithelial Cells |
| Protein Ubiquitination Pathway |
| Interferon Signaling |
| NRF2-mediated Oxidative Stress Response |
| Activation of IRF by Cytosolic Pattern Recognition Receptors |
| Unfolded protein response |
| Role of Pattern Recognition Receptors in Recognition of Bacteria and Viruses |
| Role of RIG1-like Receptors in Antiviral Innate Immunity |
| eNOS Signaling |
| Glucocorticoid Receptor Signaling |
| Aryl Hydrocarbon Receptor Signaling |
| Pathogenesis of Multiple Sclerosis |
| Hypoxia Signaling in the Cardiovascular System |
| Huntington's Disease Signaling |
| Choline Biosynthesis III |
| Primary Immunodeficiency Signaling |
| Xenobiotic Metabolism Signaling |
| Mitotic Roles of Polo-Like Kinase |
| IL-17A Signaling in Gastric Cells |
| Coenzyme A Biosynthesis |

| Upstream Transcriptional Regulator |
|---|
| IRF7 |
| HSF1 |
| NKX2-3 |
| STAT1 |
| IRF5 |
| IRF1 |
| CNOT7 |
| TRIM24 |
| IRF3 |
| STAT2 |
| STAT3 |
| IRF9 |
| SPI1 |
| PML |
| BRCA1 |
| SIRT1 |
| IRF2 |
| NFE2L2 |
| FBXW7 |
| TP53 |

| | | |
|---|---|---|
| DDX60L | IL7R | RSRP1 |
| DEDD2 | IRF7 | SAMD9 |
| DFNA5 | ISG15 | SAMD9L |
| DHRS1 | ITPR3 | SBDSP1 |
| DHRS11 | JMJD6 | SEC61G |
| DHX58 | KRIT1 | SELO |
| DNAJA1 | LGALS8 | SERPINH1 |
| DNAJA4 | LUC7L3 | SESTD1 |
| DNAJB1 | LY6E | SLC15A3 |
| DNAJB6 | MED12 | SLC35E2B |
| DTX3L | MFSD11 | SLC37A1 |
| EEF2K | MICB | SMARCC2 |
| EIF2AK2 | MLKL | SNX3 |
| EPSTI1 | MLLT11 | SP110 |
| ESCO2 | MRPL18 | SPATS2L |
| EVI2B | MSX1 | SQSTM1 |
| EVL | MX1 | STIP1 |
| FKBP4 | MX2 | SYNE2 |
| FTL | NAPG | TAP1 |
| GBP1 | NDRG1 | TGFA |
| GCH1 | NQO1 | TKT |

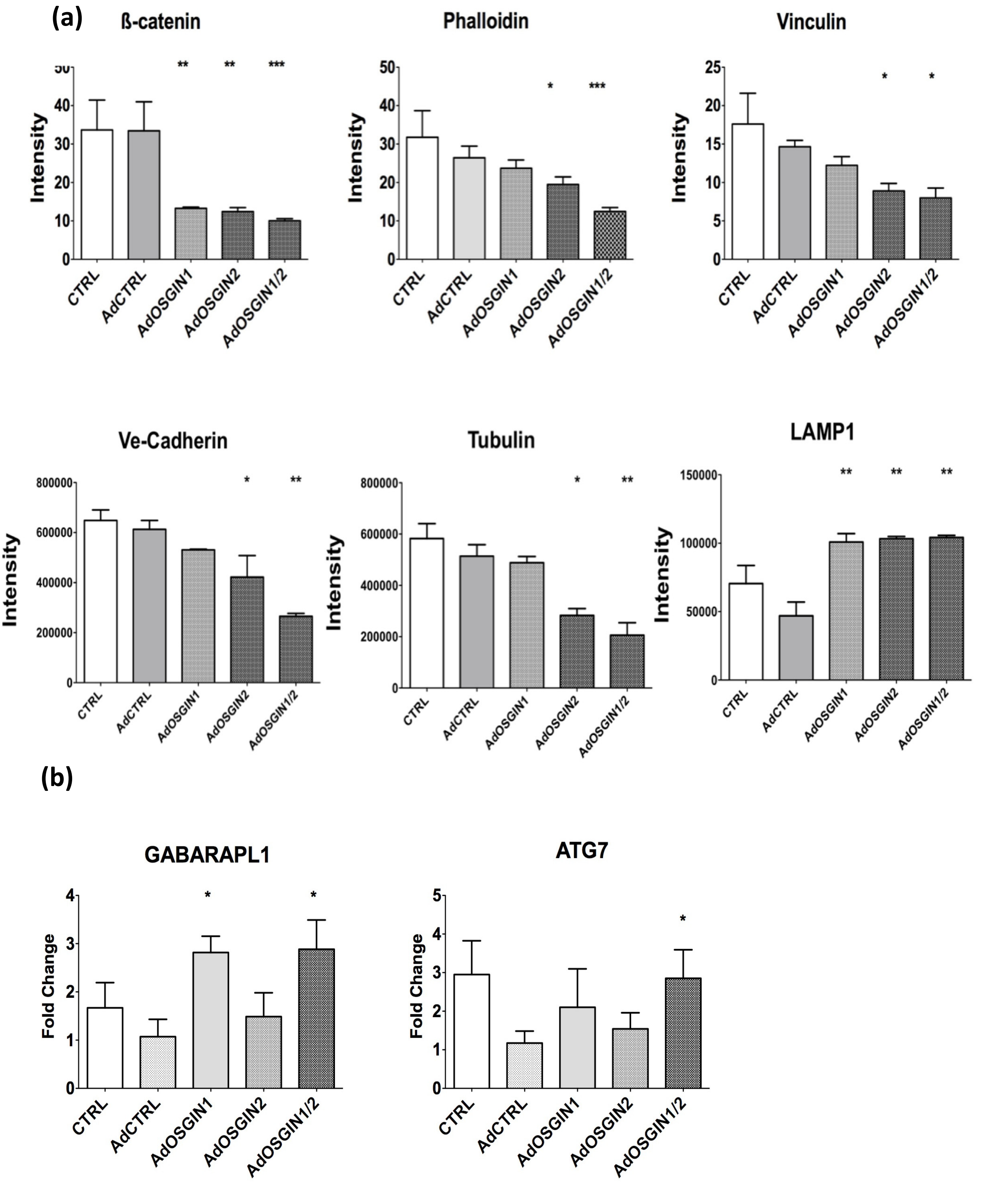

Figure S19: a)Immunocytochemical analysis indicated loss of focal adhesions, stress fibres and tubulin (n=3; P<0.05, P<0.01 and P<0.001), LAMP1 accumulation around nuclei was reported. b) GABARAPL1 (in AdOSGIN1 and AdOSGIN1+2) and ATG7 (AdOSGIN1+2) gene expression level increased compared to AdCTRL *P<0.05.

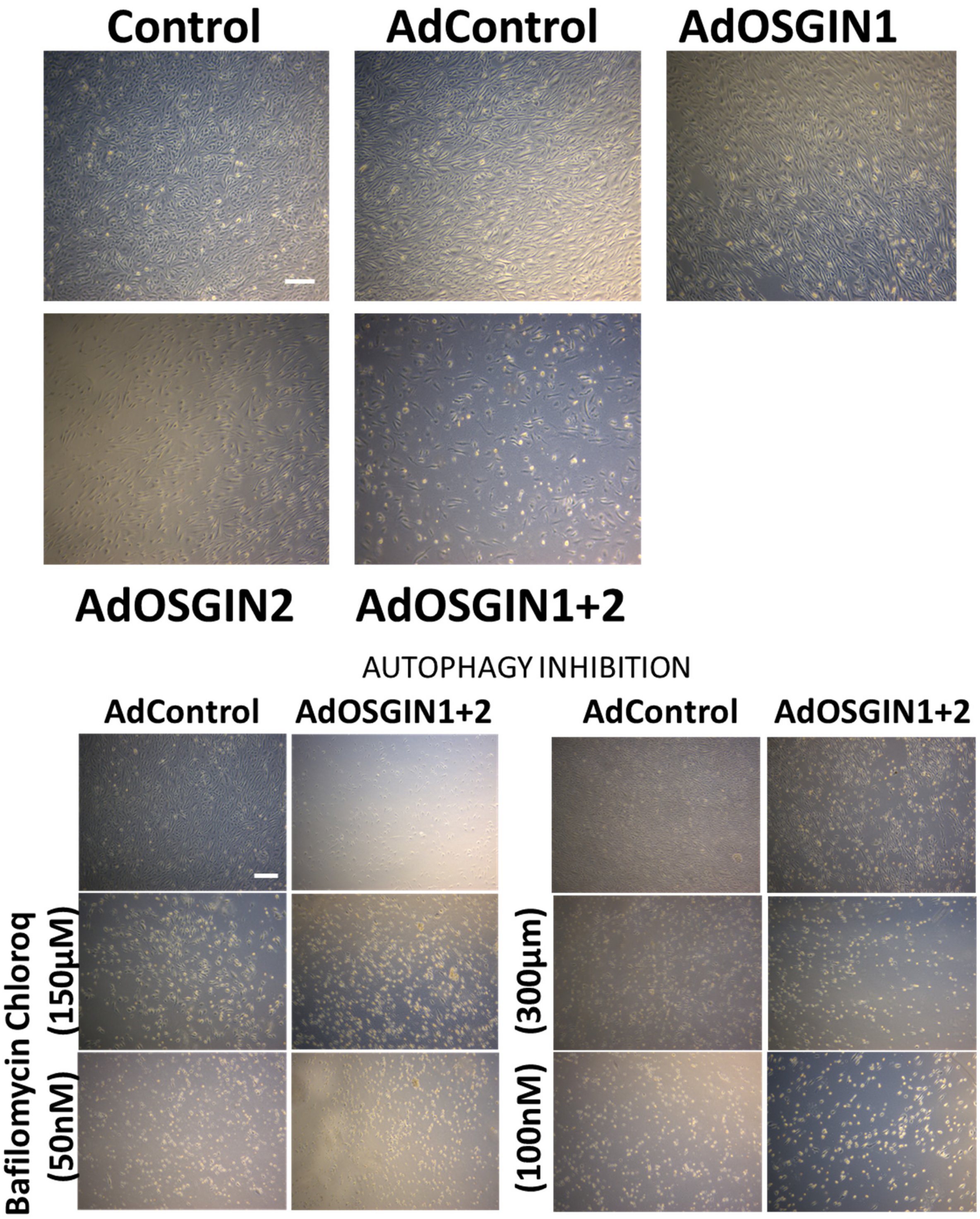

Figure S20. HCAECs were also exposed to variable levels of shear stress using an orbital shaker. Autophagy inhibitors did not rescue cell detachment, (n=3; P<0.05, P<0.01).

### RESCUE EXPERIMENT

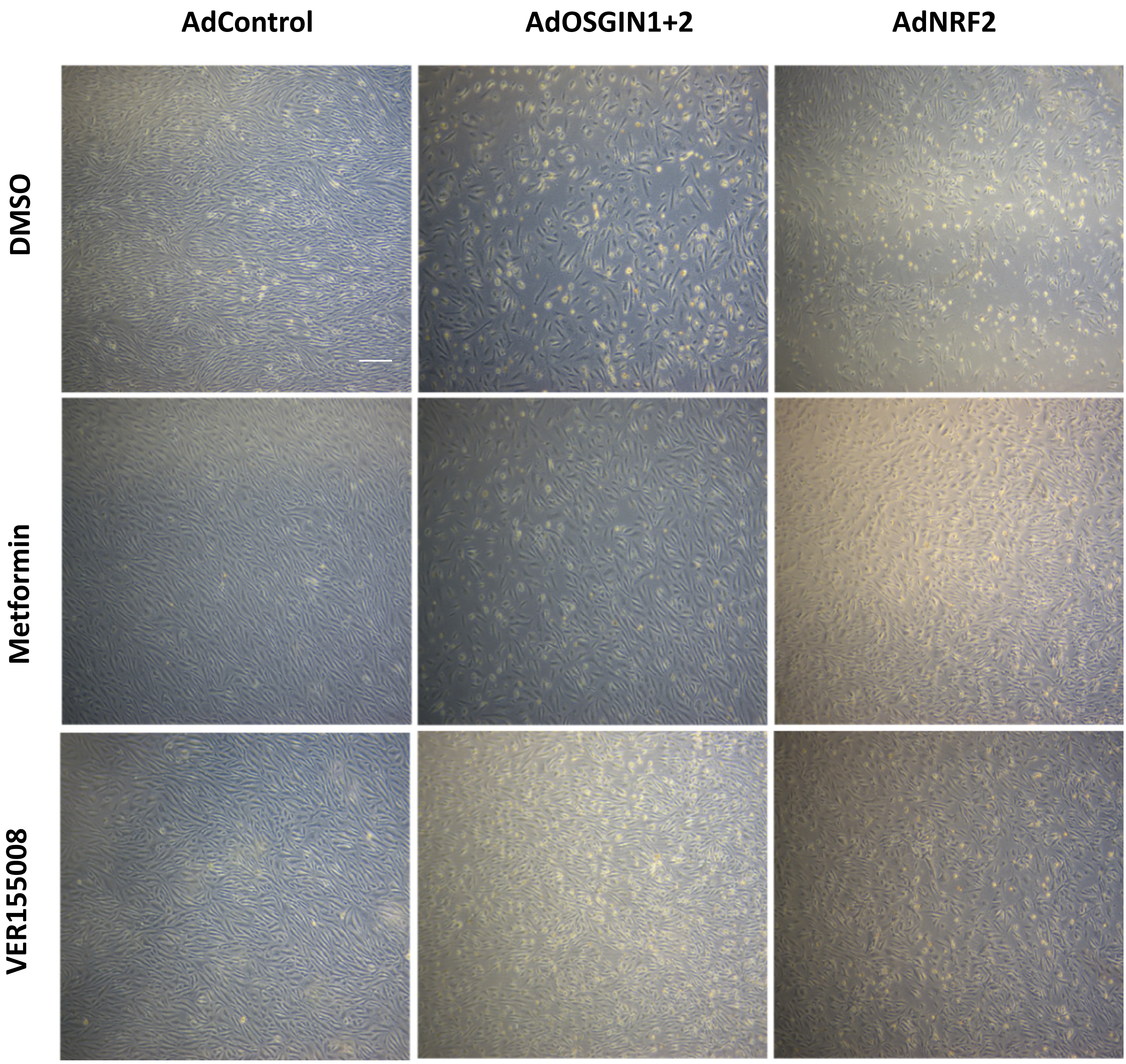

Figure S21: Confluent layer of ECs was seeded on a six-well plate. Adenoviral overexpression (AdCTRL, AdOSGIN1+2, and AdNRF2 was carried out. Following adenoviral transfection, EC were flowed on orbital shaker (210rpm, 3ml) in combination with DMSO, Metformin, and VER-155008) for 72hrs. (See quantification Figure 5E main text)

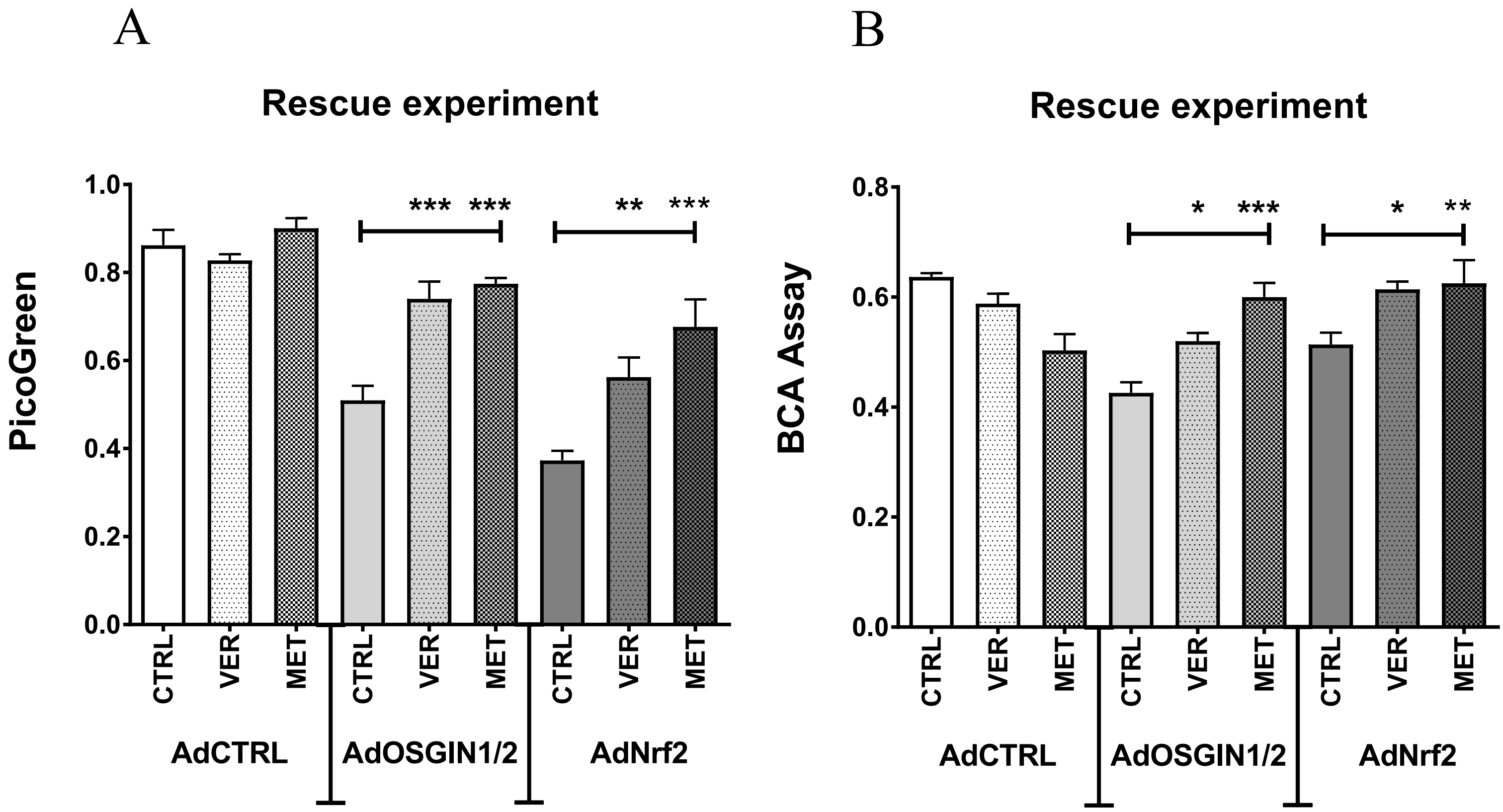

Figure S22. Comparison of different normalisation techniques. Because of the change in cell size, depth ( and therefore assumed volume) and increase in number of nuclei per cell (Figures S14c,d; S15; S16), we calculated cell detachment as % coverage (Figure 5E main text). We repeated the analysis of detachment using by A) DNA quantification [25-27] or B) protein content of the cell lysate. These gave equivalent result as presented in Figure 5E. OSGIN1+2 or Nrf2-mediated cell detachment was reduced by co-treatment with Ver155008 (15µM), or Metformin (100µM); (*P<0.05, **P<0.01, ***P<0.001, n=3, Two-way ANOVA).

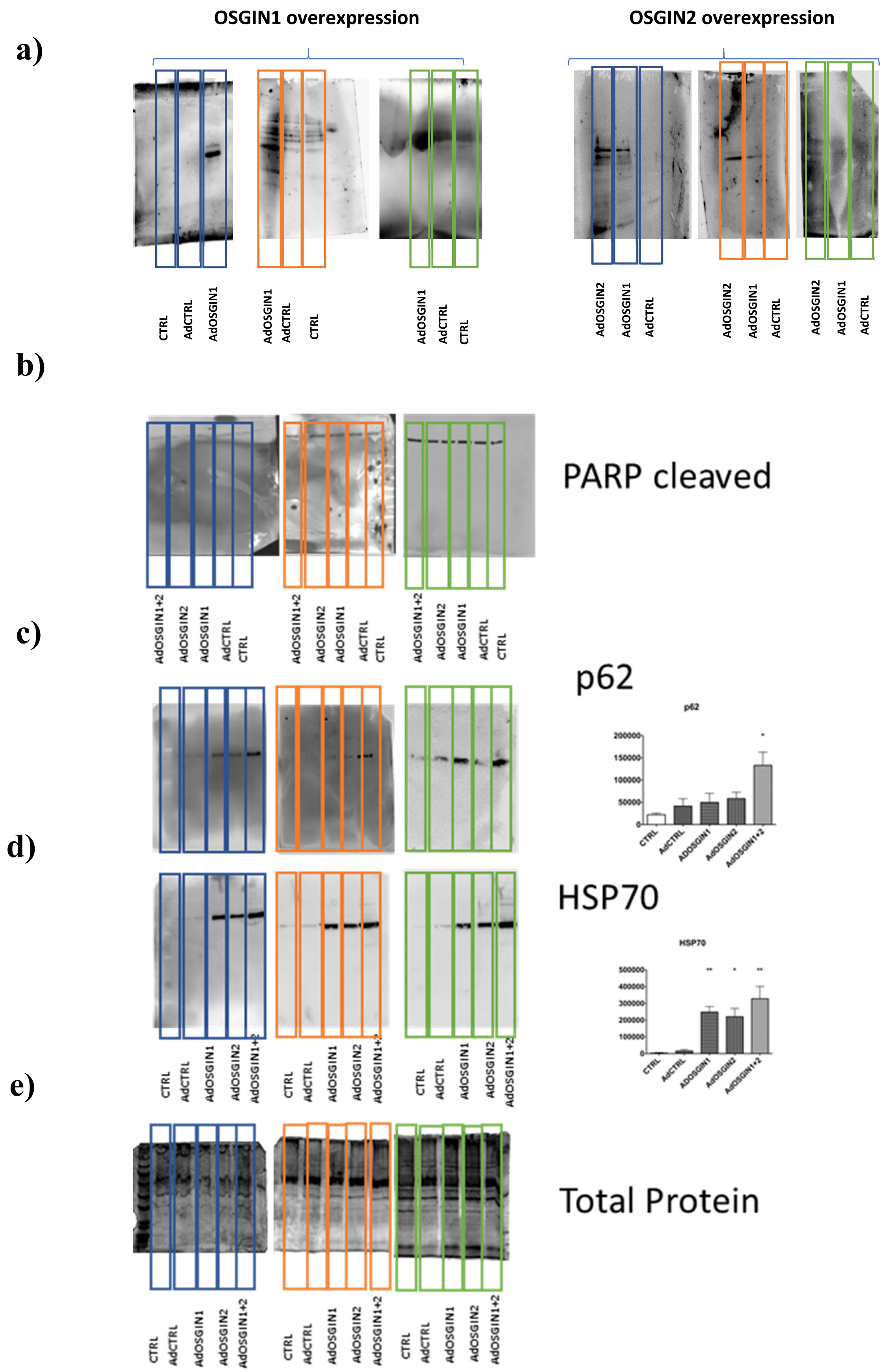

Figure S23: a) Overexpression of OSGINs was evaluated through western blotting. b) PARP cleavage antibody didn't show any cleave of the PARP protein confirming it was no apaptotic pathway related. c) Total protein (e) quantification was used to determine p62 and HSP70 accumulation

**A pivotal role for Nrf2 in endothelial detachment– implications for endothelial erosion of stenotic plaques.** Sandro Satta, *et. al.*

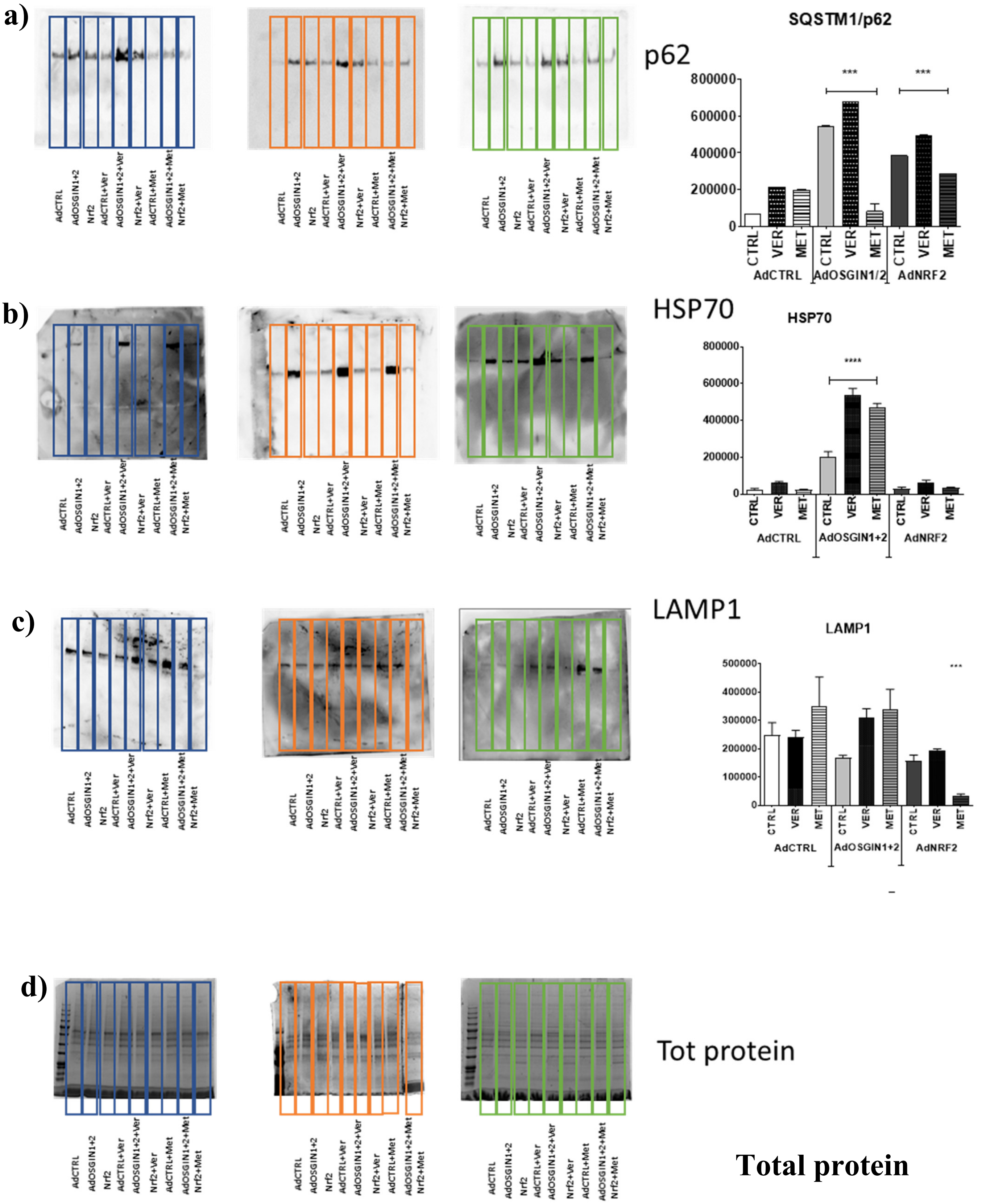

Figure S24: Rescue experiment was carried on orbital shaker using Metformin and Ver155008. Following orbital shaker experiment ECs were lysate and total protein (d) quantification was used to determine (a)p62, (b)HSP70 and (c)LAMP1(western blotting analysis).

**A pivotal role for Nrf2 in endothelial detachment– implications for endothelial erosion of stenotic plaques.** Sandro Satta, *et. al.*
